# Cancer vs. Conversational Artificial Intelligence

**DOI:** 10.1101/2024.12.28.630597

**Authors:** Kevin Kawchak

## Abstract

Solving cancer mechanisms is challenging due to the complexity of the disease integrated with many approaches that researchers take. In this study, information retrieval was performed on 40 oncological papers to obtain authors’ methods regarding the tumor immune microenvironment (TIME) or organ-specific research. 20 TIME summaries were combined and analyzed to yield valuable insights regarding how research based papers compliment information from review papers using Large Language Model (LLM) in-context comparisons, followed by code generation to illustrate each of the authors’ methods in a knowledge graph. Next, the 20 combined organ-specific emerging papers impacting historical papers was obtained to serve as a source of data to update a mechanism by Zhang, Y., et al., which was further translated into code by the LLM. The new signaling pathway incorporated four additional authors’ area of cancer research followed by the benefit they could have on the original Zhang, Y., et al. pathway. The 40 papers in the study represented over 600,000 words which were focused to specific areas totaling approximately 17,000 words represented by detailed and reproducible reports by Clau-3Opus. ChatGPT o1 provided advanced reasoning based on these authors’ methods with extensive correlations and citations. Python or LaTeX code generated by ChatGPT o1 added methods to visualize Conversational AI findings to better understand the intricate nature of cancer research.

## 1 Introduction

LLMs have been increasingly utilized for cancer research in a number of applications. In April 2023, Naik, H., et al. utilized an earlier version of ChatGPT to generate a single patient case report regarding synchronous bilateral breast cancer [1]. In September of the same year, Choi, H., et al. collected data from reports of surgical pathology and ultrasounds from 2,931 breast cancer patients, and extracted the information using ChatGPT 3.5 to efficiently derive clinical factors, obtaining an overall accuracy of 87.7% [2]. On October 16, 2023, Griewing, S., et al. in *Journal of Personalized Medicine* compared the concordance of treatment recommendations from “ChatGPT 3.5 with those of a multidisciplinary tumor board for breast cancer (MTB).” The authors found that “Overall concordance between the LLM and MTB was reached for half of the patient profiles” [3].

**Figure 1:**
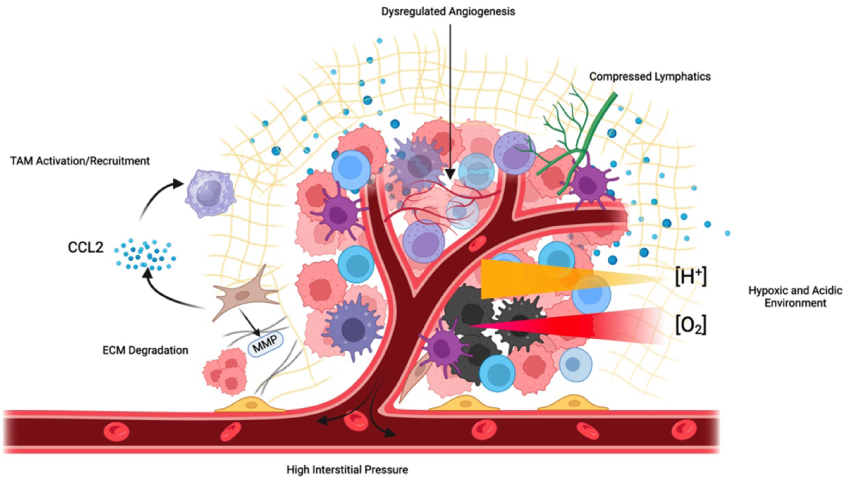
Tumor microenvironment, Garlisi, B., et al. 2024 [4]

Furthermore, in November 2023 Sorin, V., et al. utilized ChatGPT-3.5 or GPT-4 to process breast cancer clinical notes, question-answering based on guidelines, and patients’ management recommendations. “The rate of correct answers varied from 64-98%, with the highest accuracy (88-98%) observed in information extraction and question-answering tasks” [5]. A study published on November 17, 2023 focused on utilizing a variety of LLMs to examine “10 fictional cases of patients with advanced cancer with genetic alterations. Each case was submitted to 4 different LLMs (ChatGPT, Galactica, Perplexity, and BioMedLM) and 1 expert physician”. Results of 4 lung cancer cases and 6 other cancer cases indicated that the LLMs in precision oncology did not yet reach the quality and credibility of human experts [6].

In January 2024, Zack, T., et al. utilized GPT3.5-turbo and GPT4 “to infer disease status, response to treatment, and location of disease for patients with pancreatic adenocarcinoma from clinical radiology reports.” The authors used 200 deidentified radiology reports from pancreatic cancer patients to accurately interpret multiple clinically relevant features, with precision improved markedly with GPT 4 compared to GPT-3.5 [7]. Later in March, Tariq, A., et al. focused on prostate cancer consisting of “more than 1.8 million clinical notes and radiology and pathology reports for 15341 patients” in Mayo Clinic across three sites and outpatient clinics.” The authors’ LLM surpassed GPT-2 in all tasks, and improved upon the BioGPT model in one of the two tasks [8].

Also in March, Sorin, V., et al. performed six breast cancer studies evaluating either ChatGPT-3.5 or GPT-4 to explore “clinical notes analysis, guideline-based question-answering, and patient management recommendations”, with accuracy varying between studies from 50 to 98%. “Higher accuracy was seen in structured tasks like information retrieval” [9]. Iivanainen, S., et al. utilized in-context learning (ICL) and Retrieval Augmented Generation (RAG) with OpenAI’s GPT4 Turbo to field 11 questions regarding small cell lung cancer and 13 questions about non-small cell lung cancer treatment “for responses using ESMO guidelines having oncologists’ consensus, ICL with maximum context and ICL-RAG respectively provided accurate responses for 83.3%, and 79.2% of questions vs. 62.5% for the base GPT4 model. Results were more favorable using National Comprehensive Cancer Network (NCCN guidelines for ICL-RAG at 83.3%, GPT4-T at 75.0% and ICL-MC for 33.3% of questions” [10].

On the same date, Lammert, J., et al. in *Journal of Clinical Oncology* utilized a Retrieval augmented generation (RAG) system that integrates PubMed clinical studies, trial databases and oncological guidelines with LLMs to support targeted treatment recommendations. The authors “used 10 publicly accessible fictional patient cases with 7 tumor types and 59 distinct molecular alterations”, including their MEREDITH (Medical Evidence Retrieval and Data Integration for Tailored Healthcare) LLM system consisting of Google’s Gemini Pro, with a mean semantic textual similarity of LLM responses increasing from 0.69 to 0.76 (p <0.001)” [11]. A third May 29 publication by Bibault, J., et al. performed experiments using a GPT-4-based web application for monitoring breast cancer treatment toxicity. A natural language summary of the patient’s responses was generated using the GPT-4 API for physician review: “The mean time for textual summary generation was 7 (5.7-9.2) seconds. The AI-Symptom Summarization Tool (ASST) mean scores were 4.25 for accuracy and 4 for thoroughness” [12].

On May 30, Sun, C., et al. profiled histopathological and molecular alterations in squamous cervical cancer, and used ChatGPT for interpretation, reasoning, and understanding of multi-modal data of 114 Chinese patients. The authors “implemented an immersive-knowledge prompting (iKLP) strategy to trigger LLMs, which interpreted 17.8%-20.3% of omic alterations known to be associated with cancer.” “With experimental validations, LLM-reasoning showed >2-fold increased confidence for 68.5% of analyzed molecules” [13]. June 18, 2024 Longwell, J., et al. evaluated 8 LLMs, with ChatGPT-4 “correctly answered 125 of 147 questions” of examination-style multiple-choice questions from the American Society of Oncology, the European Society of Medical Oncology, and an original set from the authors. ChatGPT-3.5, and six open-source LLMs with publicly available weights were also used in the study and had lower performance than ChatGPT-4” [14]. Manjunath, P., et al. in July demonstrated that off-the-shelf LLMs could enhance their system’s accessibility through retrieval report summarization and user Q&A interactions” [15]. Later in July, Park, J., et al. employed EHR data and a GenePT model “to leverage NCBI text descriptions of individual genes with Open AI GPT-3.5 to generate gene embeddings for various downstream tasks”, with a “sensitivity increased by 10% (95%CI 7% – 11%) at specificities ranging from 99.0% to 99.9% for predictions made 0-3 months earlier, and by 22% (95%CI-4% – 48%)” [16].

On August 19, 2024, Pan, S., et al. employed a pre-trained single-cell large language model (LLM) to develop an EMT-language model (EMT-LM) for capturing discrete states within the EMT continuum in single cell cancer data. The authors achieved an AUROC of 90% across multiple cancer types, referred to as ‘scMultiNet’ [17]. An additional August publication by Alasker, A., et al. in *BMC Urology* presented a total of 52 questions on general knowledge, diagnosis, treatment, and prevention of PCa were provided to three LLMs. ChatGPT-3.5 “demonstrated superiority over the other LLMs in terms of general knowledge of PCa (p=0.018).” “ChatGPT-4 achieved greater overall comprehensiveness than ChatGPT-3.5 and Bard (p=0.028).”, with Google Bard “generating simpler sentences with the highest FRE score (54.7, p<0.001) and lowest FK reading level (10.2, p<0.001)” [18].

On August 29, Ahmad, N., et al. published CanPrompt to mitigate the accuracy and hallucination concerns to ensure responsible deployment. The CanPrompt strategy utilized “prompt engineering combined with few-shot and in-context learning to significantly enhance model accuracy by generating more relevant answers.” “After applying CanPrompt with models Mistral 7×8b, Falcon 40b, and Llama 3-8b, BERTScore results showed Mistral leading with an accuracy around 84%” [19]. In September of the same year, Li, M., et al. utilized a 7B parameter CancerLLM model “pre-trained on 2,676,642 clinical notes and 515,524 pathology reports covering 17 cancer types, followed by fine-tuning on three cancer-relevant tasks, including cancer phenotypes extraction, and cancer diagnosis generation. CancerLLM achieved state-of-the-art results compared to other existing LLMs, with an average F1 score improvement of 7.61%” [20].

On September 25, 2024, Hao, Y., et al. featured RadOnc-GPT, a specialized Large Language Model (LLM) “powered by GPT-4 that with a focus on radiotherapeutic treatment of prostate cancer with advanced prompt engineering.” The authors implemented patient electronic health records (EHR), with 158 previously recorded in-basket message interactions.” The authors estimated RadOnc-GPT “to save 5.2 minutes per message for nurses and 2.4 minutes for clinicians, from reading the inquiry to sending the response.” [21] Two days later, Hao, Y., et al. published on MedEduChat for prostate cancer patient education, which integrates with patients’ electronic health record data and features a closed-domain, semi-structured, patient-centered approach to address real-world needs. The authors then “integrated MedEduChat with OpenAI’s GPT-4o through the clinic’s Azurehosted endpoint, which is HIPAA compliant.” [22].

On October 1, Gilbert, M. utilized a Mixtral 8×7B LLM by Mistral AI “to automate the extraction of key words and phrases from multidisciplinary head and neck cancer tumor board notes of 50 patients diagnosed and treated in 2021”. The authors “found collectively that the precision, recall, F1 score, and accuracy were 96.9%, 95.7%, 96.3%, and 93.1%, respectively” [23]. On the same day, Khanmohammadi, R., et al. proposed a novel student-teacher LLM architecture that self-improved the key concept abstraction through automatic prompt optimization, and was designed for local use to safeguard patient privacy in oncology. The Mixtral-8×7B student model initially extracted symptoms and treatments from given prompts, “which were then refined by the GPT-4 teacher model.” The performance for multi-symptom notes “demonstrated average improvement in accuracy from 0.24 to 0.43, precision from 0.60 to 0.76, recall from 0.24 to 0.43 and F1 from 0.20 to 0.44.” [24]

On October 3, Kim, K., et al. integrated prognostic capabilities of both CT and pathology images with clinical information for lung cancer, employing a multi-modal integration approach “via multiple instance learning, leveraging large language models (LLMs) to analyze clinical notes and align them with image modalities” [25]. Two days later, Das, R., et al. presented GeneSilico Copilot, “an advanced agent-based architecture that transforms LLMs from simple response synthesizers to clinical reasoning systems” for precision oncology. The authors employed state-of-the-art LLM services, GPT-4 and Claude Opus-3, known for their long context windows” [26]. Li, Y., et al. on October 11 published in *IEEE Journal of Biomedical and Health Informatics* regarding a large language model (LLM)-based Knowledge-aware Attention Network (LKAN) for clinical staging of liver cancer (CSoLC). The LLM and a rule-based algorithm were integrated to generate more diverse and reasonable data, with unlabeled radiology corpus of liver cancer being pre-trained, and attention was improved by incorporating both global and local features. The classification accuracy of LKAN “achieved the best results with 90.3% Accuracy, 90.0% Macro_F1 score, and 90.0%”

Four days later, Yang, S., et al. provided a computational approach called LLM4THP to quickly and effectively detect tumor homing peptides (THPs). LLM4THP utilized LightGBM, XGBoost, Random Forest, and Extremely Randomized Trees, and “Logistic Regression to further refine the identification of sequences as either THP or non-THP.” LLM4THP outperformed other compared methods in terms of ACC, MCC, F1, AUC and AP “with improvement by 2.3–4.61%, 4.63–8.79%, 2.22–3.95%, 1.94% to 3.46 and 2.7–5.91%” on the primary test dataset” [27]. On October 21, Gubanov, M., et al. verified that knowledge graph served as a Retrieval Augmented Generation (RAG) guardrail. Their CancerKG model exhibitsed5 different advanced user interfaces, “each tailored to serve different data modalities better and more convenient for the user.” The authors evaluated CancerKG on real user queries and reported a high normalized discounted cumulative gain score on a large-scale corpora of approximately 44K publications” [28].

Oh, Y., et al. in *Nature Communications* on October 24, 2024 presented a LLM-driven multimodal artificial intelligence (AI), “namely LLMSeg, that utilizes the clinical information and is applicable to the challenging task of 3-dimensional context-aware target volume delineation for radiation oncology.” The authors demonstrated that LLMSeg exhibited markedly improved performance compared to conventional unimodal AI models, particularly exhibiting robust generalization performance and data-efficiency” [29]. On the same day, Yu, J., et al. used LLMs to screen “36,105 EBV-relevant scientific publications and summarize the current literature landscape on various EBV-associated diseases like Burkitt lymphoma (BL), diffuse large B-cell lymphoma (DLBCL), nasopharyngeal carcinoma (NPC), and so on. The accuracy of the GPT-generated summary greatly depended on the concise prompt input with clear instructions, which can enhance the precision of the generated response.” To optimize costs and efficiency, the authors initially utilized GPT-3.5 Turbo followed processing with GPT-4 [30]. Lastly, Lammert, J., et al. on October 30, 2024 published in *JCO Precision Oncology* on utilizing their LLM system MEREDITH to support treatment recommendations in precision oncology. The system was built on Google Gemini Pro, with MEREDITH using retrieval-augmented generation and chain of thought, with concordance between LLM suggestions and expert recommendations of 94.7% for the enhanced system [31].

Thus, from April 2023 to recent studies, LLMs have been utilized in an increasing number of applications for cancer research, as readily apparent with the ten October 2024 articles shown here in journals such as *Nature Communications* and *JCO Precision Oncology*. Authors continue to utilize newer LLMs to increase the performance of their studies as evidenced by the Choi, H., et al. September 2023 inclusion of ChatGPT 3.5, transitioning to the more capable GPT-4o model by Hao, Y., et al. in September 2024. Technologies such as information retrieval implemented by Sorin, V., et al., in-context learning and retrieval augmented generation by Iivanainen, S., et al., and RAG also by Gubanov, M., et al. are examples of similar methods being utilized in this study. In addition, the Khanmohammadi, R., et al. method of incorporating models from different AI software manufacturers was also implemented in this current study. More recent studies showed effective use of a) information retrieval techniques across a number of papers to b) update current research approaches. Note: At time of submission, additional ChatGPT 4o Cancer and “ChatGPT o1” Cancer searches were made, however nothing corresponding to the scope of this paper was immediately found.

**Figure 2:**
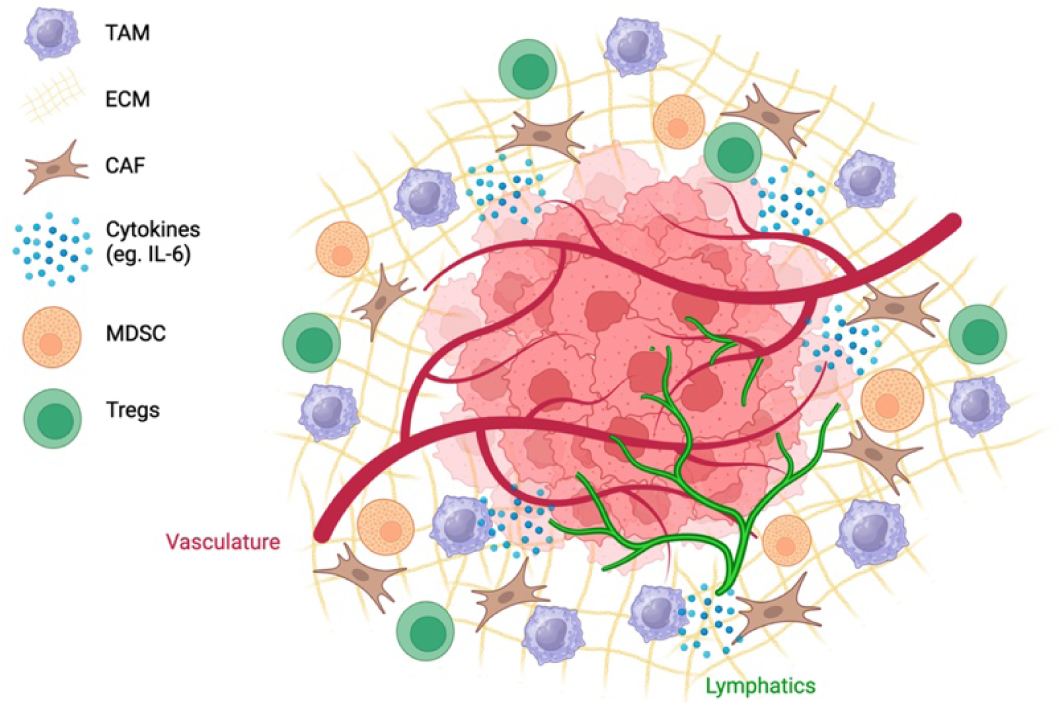
Complex TME with lymphatics, Garlisi, B., et al. 2024 [4]

## 2 Methods

AI software used in this study were all unmodified LLMs with generation times being obtained with a digital stop watch. Clau-3Opus utilized document retrieval of pdfs found only in Table 2 using the ‘paper clip’ option and text inputs, while Lla3.1-405 and ChatGPT o1 processed text inputs. ChatGPT o1 prompts were first optimized, then ran as a chain of prompts in a single conversation. Text that was copied into LLM input fields was also pasted as plain text into Supplementary SIM, SIMp, SOR, SORp, and S40p; with a triplicate reproducibility study shown in Supplementary SREP for a total across files of 61 generations. Generations in the manuscript received white space formatting for readability. The author conducted experiments, analysis, and wrote the manuscript, with ChatGPT 4o being primarily utilized as a research tool. Paper lengths were obtained by copying each pdf into a Google Docs and using the Word Counts feature. JupyterLab [32] was accessed through Anaconda [33] and MacOS 14.5 (23F79) in Google Chrome browser Version 131.0.6778.109 (Official Build) (arm64) for running Python code, while the manuscript software natively processed LaTeX code.

1. Clau-3Opus: Claude website chat interface was accessed through MacOS 14.5 (23F79) and Google Chrome browser Version 131.0.6778.109 (Official Build) (arm64) [34].
2. Lla3.1-405: Fireworks.ai website chat interface was accessed through MacOS 14.5 (23F79) and Google Chrome browser Version 131.0.6778.109 (Official Build) (arm64). Settings: Max Tokens=4096, Temperature=1.0, Top P=1.0, Top K=50, Presence Penalty=0, Frequency Penalty=0 [35].
3. ChatGPT o1: ChatGPT website pro interface was accessed through MacOS 14.5 (23F79) and Google Chrome browser Version 131.0.6778.109 (Official Build) (arm64) [36]. The ChatGPT o1 pro model was not utilized in this study.
4. ChatGPT 4o: ChatGPT website pro interface was accessed through MacOS 14.5 (23F79) and Google Chrome browser Version 131.0.6778.109 (Official Build) (arm64) [37].

## 3 Part I: Information Retrieval and Cancer Insights

### 3.1 Part I Results: Information Retrieval and Cancer Insights

The main outcomes for Part I are as follows: Prompt 1 allowed for information retrieval of the authors’ tumor immune microenvironment methods using 20 separate generations of 20 pdfs with Clau-3Opus. Similarly, all reports returned the requested “Executive Summary,” “Technical Details,” “Key Insights” format based on the authors’ organ-specific cancer methods for Papers 21-40. In Table 2, 609,733 total words across 40 papers were analyzed, with Clau-3Opus generations extracting information totaling 17,329 words. Each generation was verified by searching for at least three paper specific search terms, with some terms being appropriately abbreviated. A portion of Papers 21-40 from journal *BMC* used technical details from the abstract, with less reliance on the rest of the paper. Each of the 40 generations contained at least one quotation from each paper, as requested in Table 1 prompts. The AI model sometimes also included additional quotations, but were typically within the context of original papers. Prompts also requested quotations of numerical data in quotation marks that assisted the AI model’s conclusions, with some reports having more data than others. In one instance, the word “tumour” used inside quotations was modified to “tumor”.

**Table 1:**
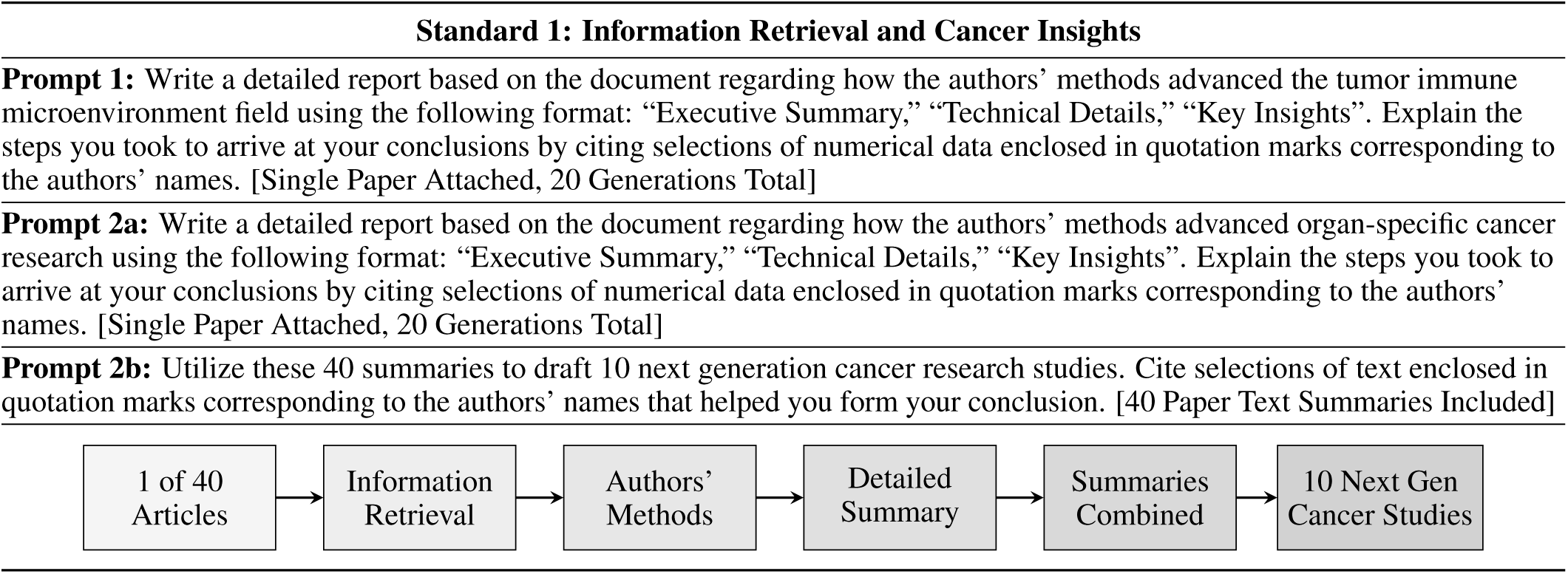
Standard 1 Process Diagram.

**Table 2:**
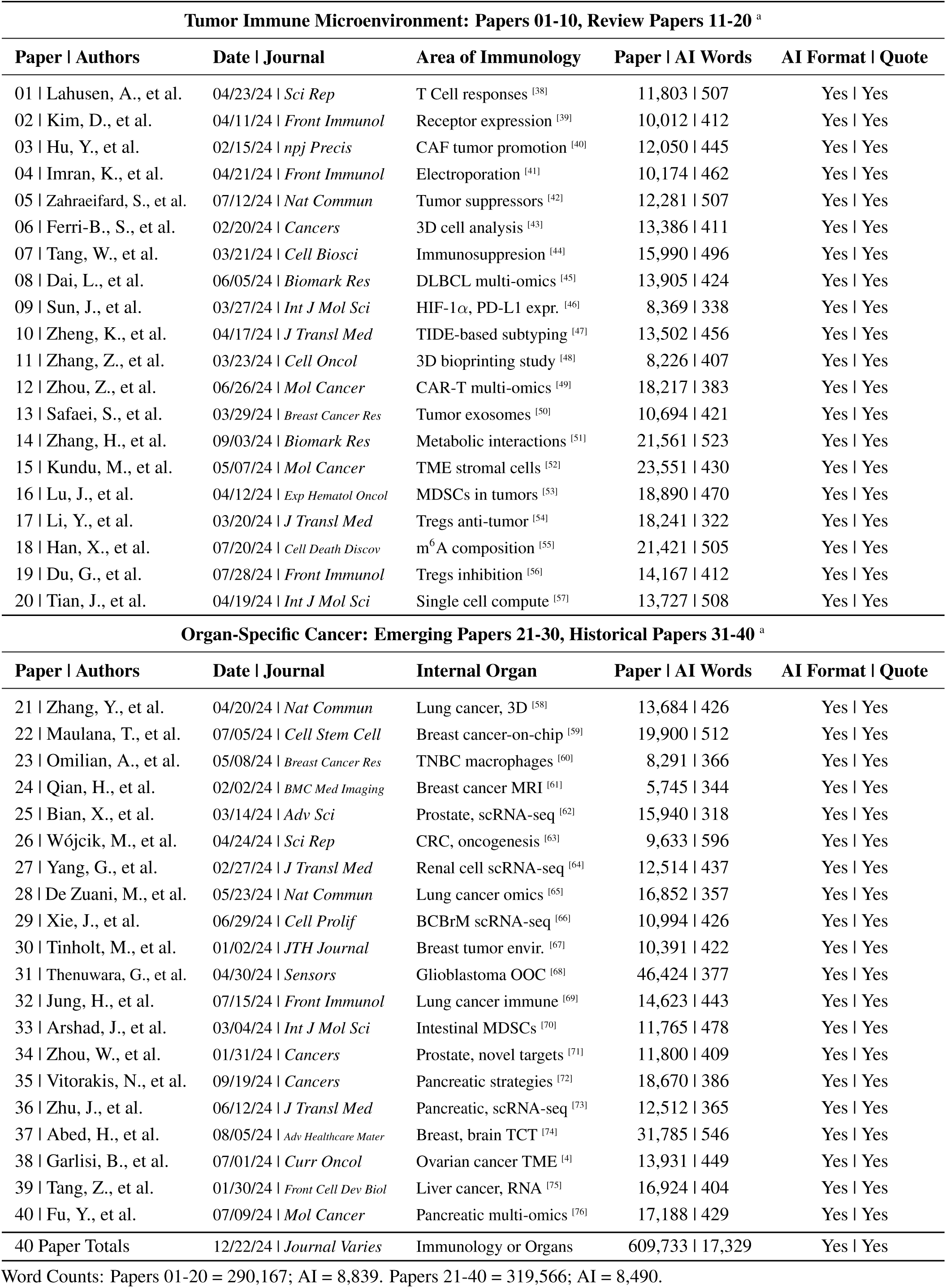
40 Papers Analyzed in Study.

Overall, Clau-3Opus retrieved consistently high quality information regarding how authors’ methods advanced cancer research, while reducing total word counts by over 35x at an average of 433 words per paper with a standard deviation of 62.8 for ChatGPT to process within its 128K context length. Papers were selected according to the PRISMA study located in Appendix Figure 8. The task of processing over 17,000 words across forty summaries to obtain a Top 10 cancer research list was provided to Lla3.1-405, but was not able to cite selections of text enclosed in quotations marks corresponding to the authors’ names, as shown in Table 4. Lla3.1-405 had some repetition between its research question and objective formatting. When focusing on an area from a specific paper, the model tended to support it with other authors’ methods as seen in Study 4: Elucidating the Role of Interleukin Enhancer-Binding Factor 2 (ILF2) in Brain Metastasis supported by single-cell RNA sequencing (Paper 25-27, 29-30, 36) and functional studies using ILF2 knockout/knockdown (Paper 03, 15, 18, 21).

**Table 3:**
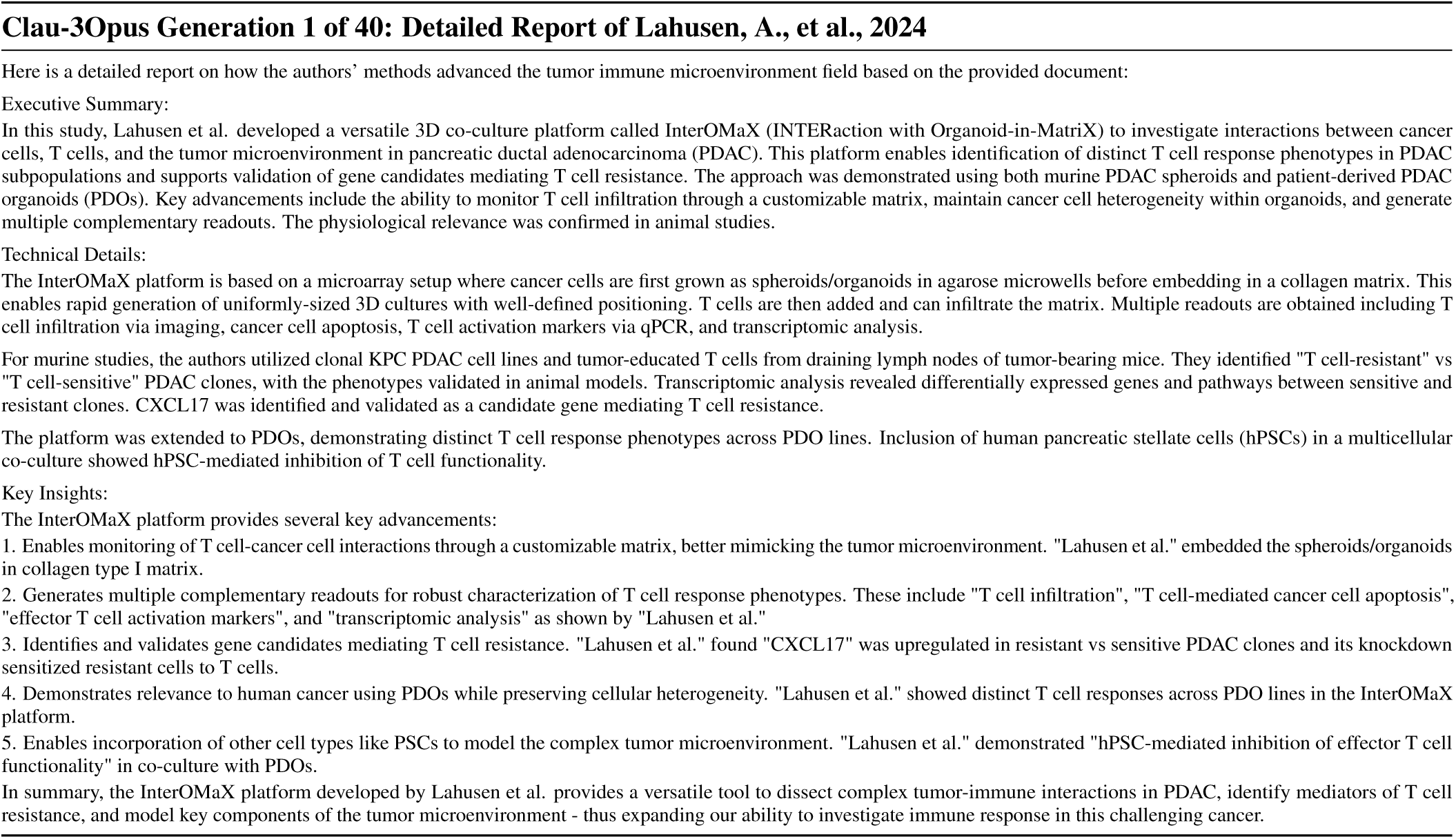
Clau-3Opus Generation 1: 507 words, 46.00s.

**Table 4:**
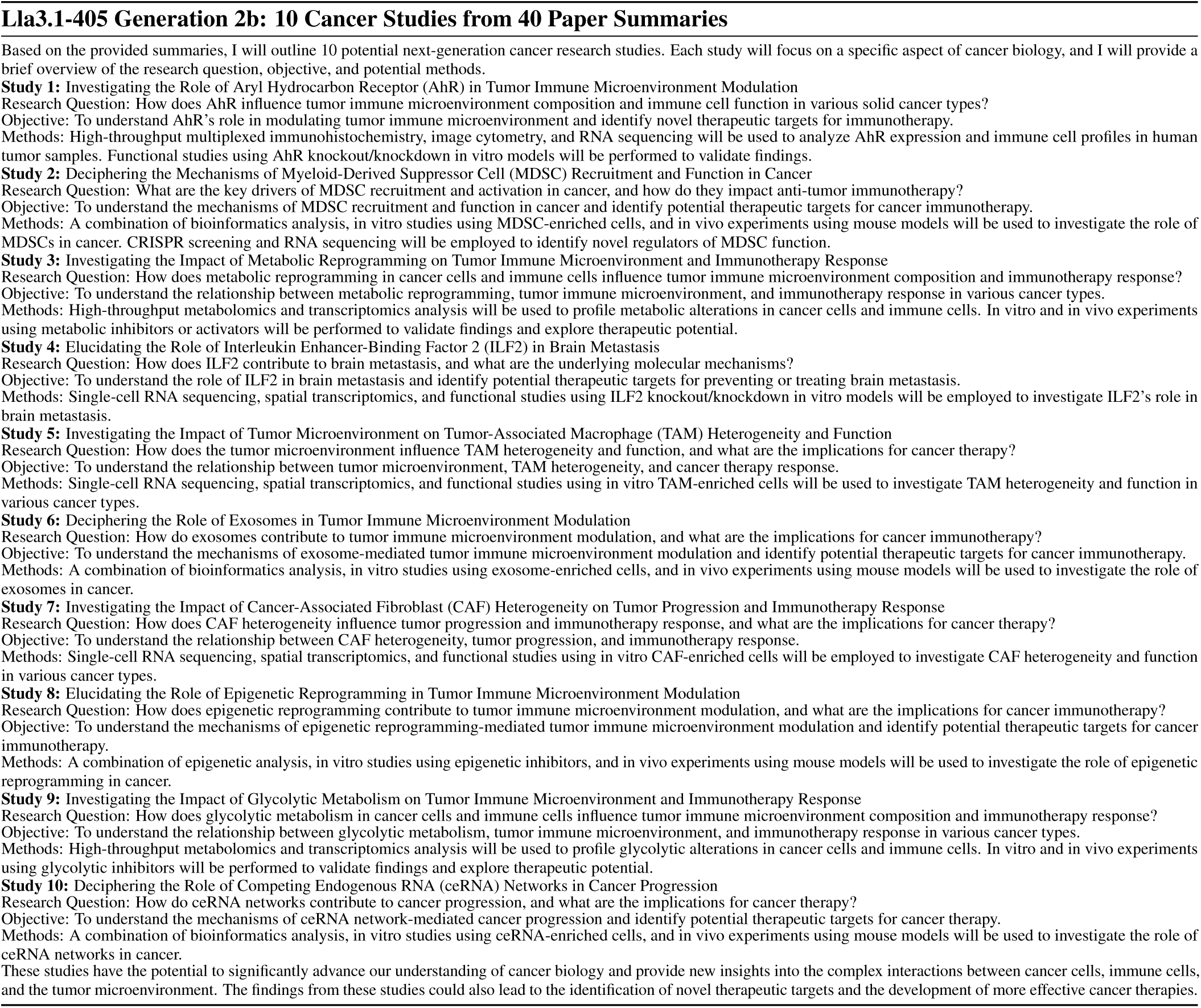
Lla3.1-405 Generation 2b: 942 words, 31.05s.

### 3.2 Part I Discussion: Information Retrieval and Cancer Insights

Clau-3Opus was chosen due to it’s ability to effectively perform document retrieval on medical literature by using the manuscript authors’ methods for each of the generations, while providing citations in quotation marks. Another model by the same manufacturer Clau-3.5So in testing tended to provide more lists, which were harder to read. OpenAI’s ChatGPT 4o also performs well with document analysis, but outputs tend to no longer utilize plain text citations and typically return a blue quote hyperlink that refers to the entire document. In addition, Clau-3Opus was utilized to compliment OpenAI’s ChatGPT o1’s strong reasoning abilities in Part II and III, as ChatGPT o1 only accepts text and images as inputs, not pdfs. The choice to request citations was due to previous success over requests for page numbers, line numbers, and word counts. The phrase “detailed report” was used over ‘report’ to increase word counts of Clau-3Opus generations. In addition, the request for AI to return authors’ names corresponding to quotations has become an important feature in many of the recent studies, [77, 78, 79]. Lla3.1-405 processed the largest text input in the study, however the generation contained moderate detail on a smaller task of generating a Top 10 research list shown in Table 4.

## 4 Part II: Tumor Immune Microenvironment

### 4.1 Part II Results: Tumor Immune Microenvironment

Table 5 depicts the two prompt sequence provided to ChatGPT o1. Prompt 3a is based on the TIME 10 Research Summaries and 10 Review Summaries generated by Clau-3Opus in Prompt 1. The AI model was tasked to its use in-context comparisons to provide how the Research Papers further complimented the Review Papers using the “Executive Summary,” “Technical Details,” “Key Insights” format. A major research objective was met between the two sets of paper reviews as the ChatGPT o1 1621 word detailed report shown in Table 6 contained many direct comparisons between several authors throughout the work such as “Tang, W. identified “SPP1+ macrophages” linked to poor prognosis in glioma, while Dai, L. found “glycolysis-high malignant B cell subsets” correlated with T cell exhaustion in DLBCL. These advances align with in Zhang, H., et al. (2024) and Han, X., et al. (2024) to understand metabolic and epitranscriptomic regulation of the TIME.”

**Table 5:**
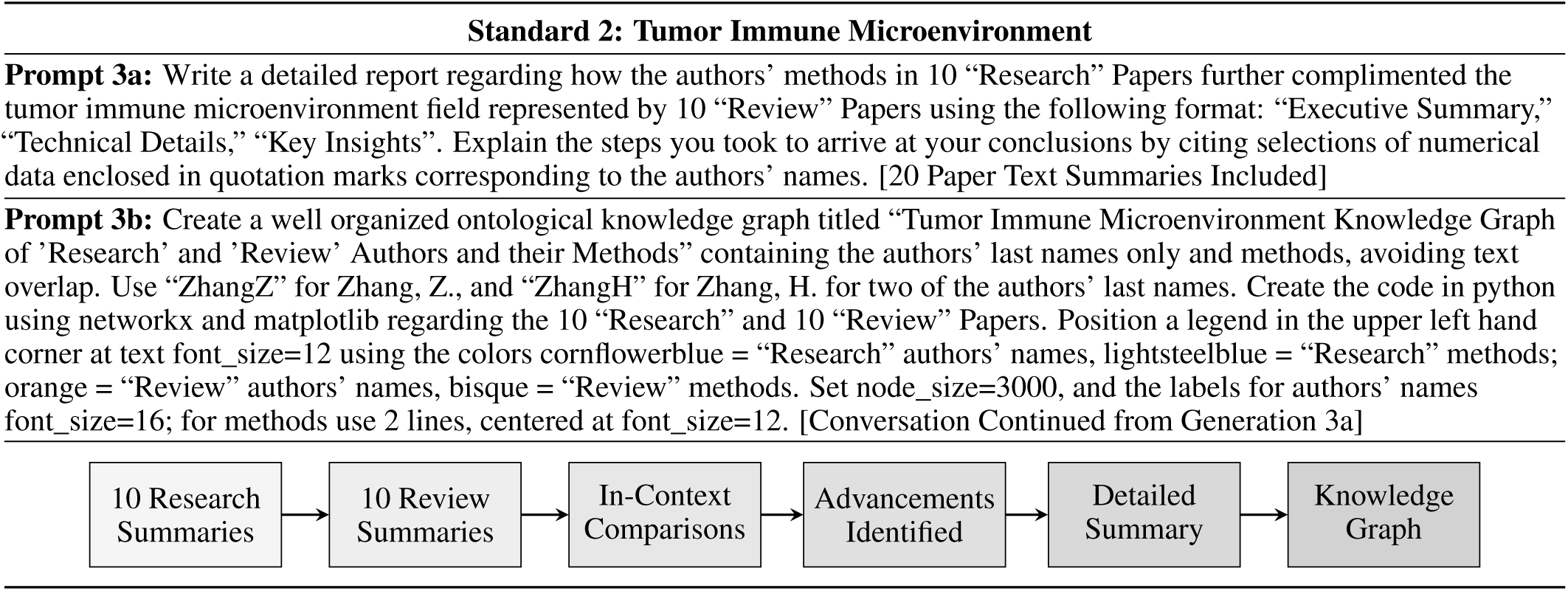
Standard 2 Process Diagram.

**Table 6:**
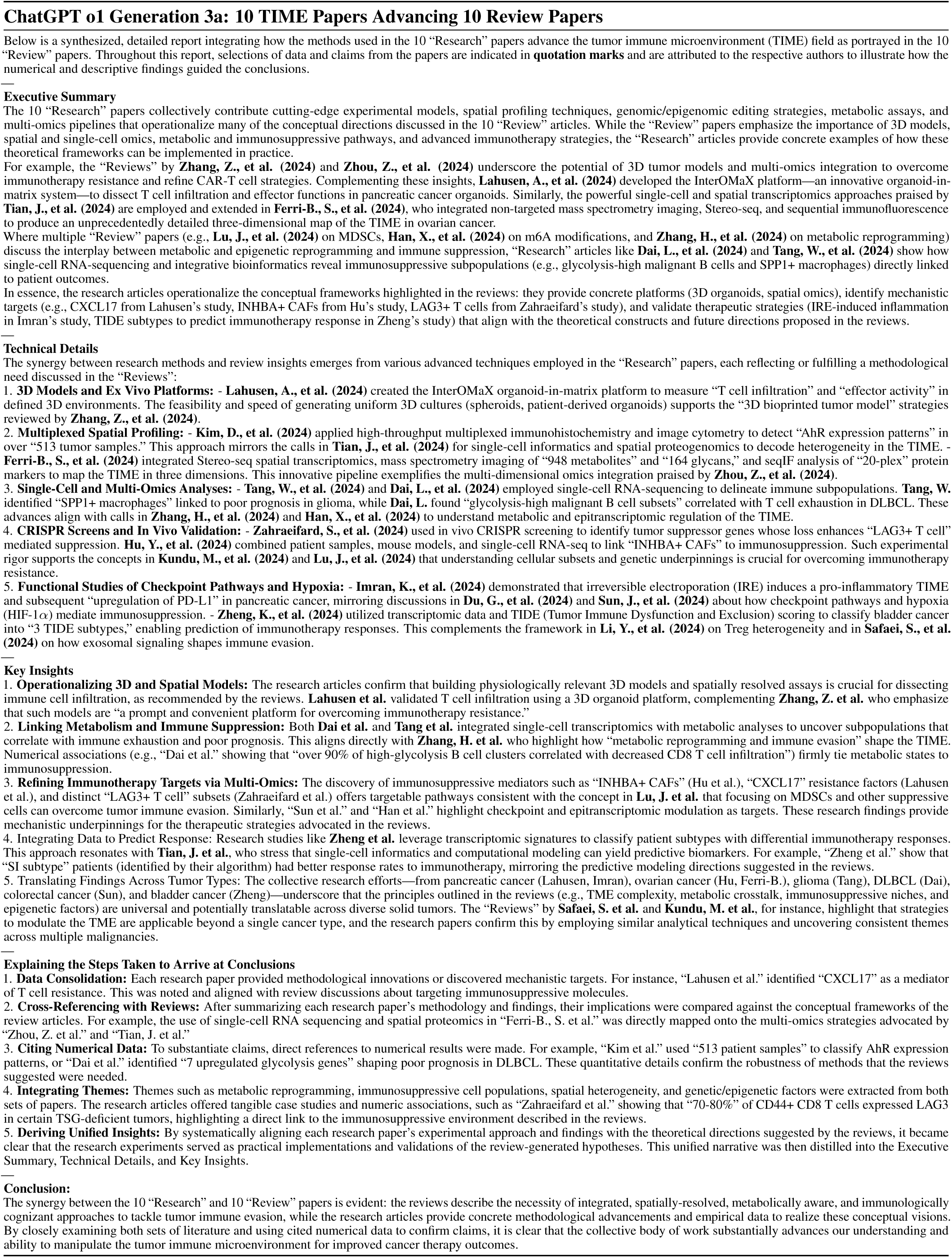
ChatGPT o1 Generation 3a: 1621 words, 57.03s.

The model also provided insights regarding how it arrived at conclusions: “Cross-Referencing with Reviews: After summarizing each research paper’s methodology and findings, their implications were compared against the conceptual frameworks of the review articles. For example, the use of single-cell RNA sequencing and spatial proteomics in “Ferri-B., S. et al.”

These correlations were directly mapped onto the multi-omics strategies advocated by “Zhou, Z. et al.” and “Tian, J. et al.””. Prompt 3b was submitted to the same conversation using ChatGPT o1 to obtain a graph visualization. The request for the AI model to provide Python code using packages networkx and matplotlib for an ontological graph was not acheived, as the model responded that it would require more information to make the graph ontological. Further attempts using ChatGPT o1 pro mode across the two prompts typically resulted in shorter outputs and less favorable graphs. Regardless, ChatGPT o1 followed several instructions to return code using 3 font size instructions, 4 color settings, a node size, text justification, and text positioning instructions shown in Table 5. Prompt 3b yielded code of a visually appealing knowledge graph, executed by ChatGPT 4o and verified in a JupyterLab Python notebook [32, 33].

### 4.2 Part II Discussion: Tumor Immune Microenvironment

Previous studies featured a maximum of 16 papers to solve problems related to drug production areas [77, 78, 79]. By relying on more input text and a recently released ChatGPT o1 for greater reasoning, an unprecedented number of comparisons and contrasts were made between authors’ cited methods from these paper summaries at over 8,800 words. The wording of Prompt 3a was optimized to obtain additional insight regarding the Research papers and Review papers when wording was modified from “advanced” and “verses” to “complimented” and “represented by”. Selections of cited numerical data in quotations were not as prevalent and robust due to a lower ability of Clau-3Opus to utilize this format, however an example includes ““70-80%” of CD44+ CD8 T cells expressed LAG3 in certain TSG-deficient tumors”. Many iterations of Prompt 3b were tested regarding Python code to obtain an appropriate graph type, color scheme, and positioning. After these settings were optimized, ChatGPT o1 outputs to Prompt 3a and 3b were run consecutively, with the final output code accurately conveying the 10 Research authors and methods separate from 10 Review authors and methods based on the prompt’s instructions represented in Figure 3.

**Figure 3:**
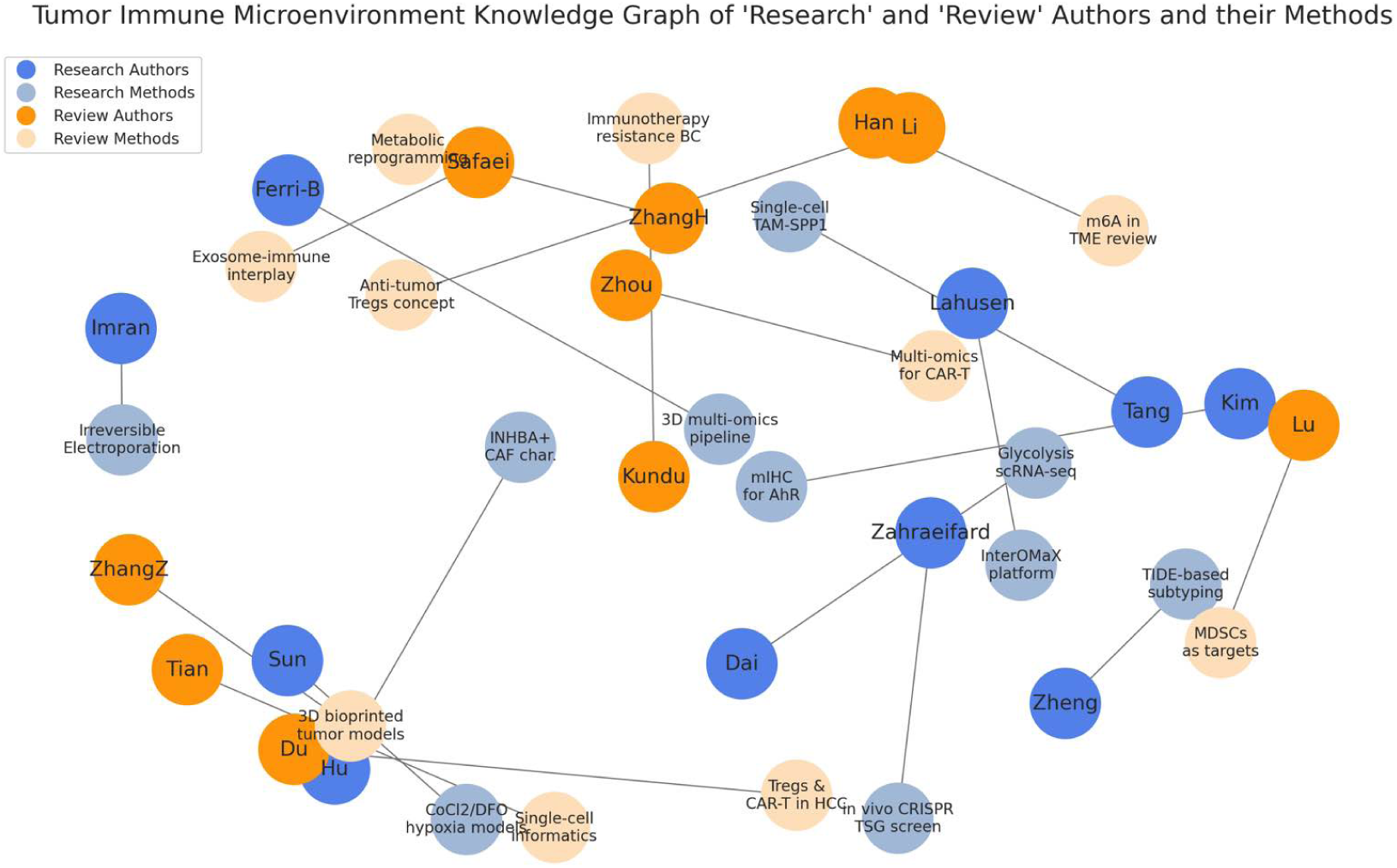
ChatGPT o1 Generation 3b

Multiple “Review” papers (e.g., **Lu, J., et al. (2024)** on MDSCs, **Han, X., et al. (2024)** on m6A modifications, and **Zhang, H., et al. (2024)** on metabolic reprogramming) discuss the interplay between metabolic and epigenetic reprogramming and immune suppression, “Research” articles like **Dai, L., et al. (2024)** and **Tang, W., et al. (2024)** show how single-cell RNA-sequencing and integrative bioinformatics reveal immunosuppressive subpopulations (e.g., glycolysis-high malignant B cells and SPP1+ macrophages) directly linked to patient outcomes. In addition, “AhR expression patterns” in over “513 tumor samples.” to 513 tumor samples vs. The authors utilized mIHC to stain tissue microarray (TMA) slides containing samples from patients with head and neck squamous cell carcinoma, bladder cancer, colorectal cancer, esophageal cancer, and non-small cell lung cancer. But corrected later: “Kim et al.” used “513 patient samples” to classify AhR expression patterns

The collective research efforts—from pancreatic cancer (Lahusen, Imran), ovarian cancer (Hu, Ferri-B.), glioma (Tang), DLBCL (Dai), colorectal cancer (Sun), and bladder cancer (Zheng)—underscore that the principles outlined in the reviews (e.g., TME complexity, metabolic crosstalk, immunosuppressive niches, and epigenetic factors) are universal and potentially translatable across diverse solid tumors. The “Reviews” by **Safaei, S. et al.** and **Kundu, M. et al.**, for instance, highlight that strategies to modulate the TME are applicable beyond a single cancer type, and the research papers confirm this by employing similar analytical techniques and uncovering consistent themes across multiple malignancies. A total of 37 citations to the 20 paper summaries totaling 1621 words were made in 57.03s. All authors in the summaries were cited at least once by ChatGPT o1, with others having two or more, with standard deviation citation per author at 0.67. The most cited authors was 3 by Lahusen, A., et al., Tang, W., et al., and Dai, L., et al.

**Figure 4:**
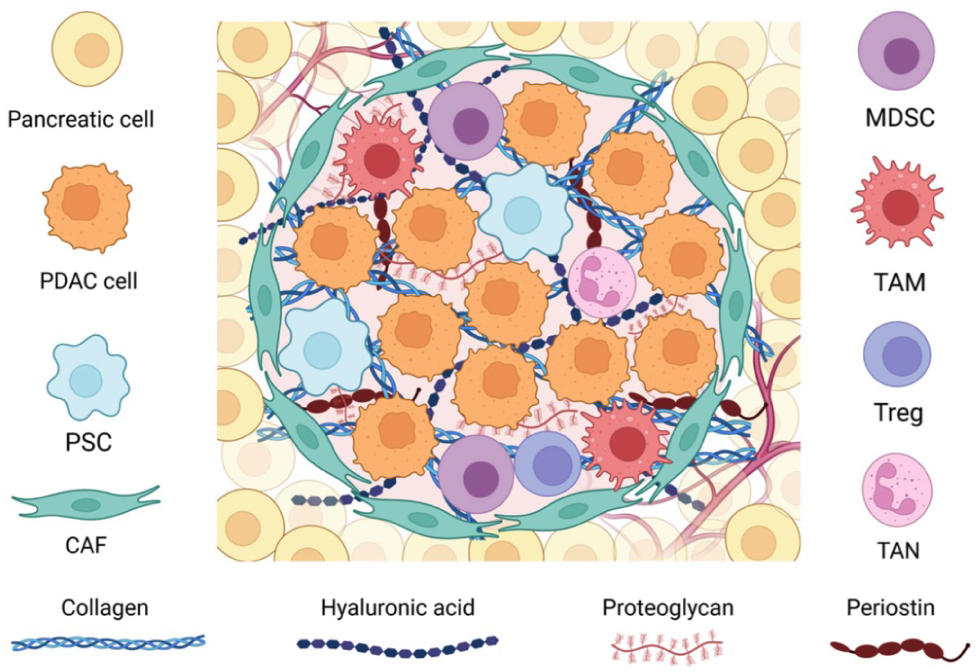
Pancreatic tumor microenvironment, Vitorakis, N., et al. 2024 [72]

## 5 Part III: Organ-Specific Cancer

### 5.1 Part III Results: Organ-Specific Cancer

The process of gaining insight using a detailed report format regarding how 10 emerging paper summaries complimented 10 historical paper summaries was similar to Part II, but this time focusing on organ-specific cancer research. Exceptional reasoning was accomplished by ChatGPT o1 in several cases as seen in Table 8. The Zhang, Y., et al. paper was copied through Data availability in HTML at 10118 words for effective processing.

a. Handling the complexities of historical papers: “Jung, H., et al. clarified chemokine pathways in lung tumors, while Garlisi, B., et al. dissected the ovarian TME, showing that “poor tumor perfusion and hypoxia” (Attane et al., cited by Garlisi, B., et al.) hindered therapy. Zhu, J., et al. and Fu, Y., et al. showed how single-cell and spatial transcriptomics in pancreatic cancer uncovered heterogeneous CAF subsets and immunosuppressive niches.”
b. Incorporating numerical findings from emerging summaries: “Emerging researchers introduced powerful methods. Zhang, Y., et al. demonstrated the generation of “approximately 200 uniform LCAs of ∼400-500 m from just 10 L of hydrogel with 106-108 cells/mL in under 1 minute,” enabling patient-specific lung cancer models. Maulana, T., et al. developed a perfused breast cancer-on-chip device, “allowing CAR-T cell infiltration and cytokine release profiling over an 8-day culture,” mirroring early organ-on-a-chip and biosensing approaches used for glioblastoma (Dornhof et al., referenced by Thenuwara, G., et al.)”
c. Setting a high standard correlating the two sets of summaries: “Historical papers identified overarching immunosuppressive players like MDSCs in GI cancers (Arshad, J., et al.) and complex immunosuppressive networks in ovarian and pancreatic cancers (Garlisi, B., et al.; Fu, Y., et al.). Emerging studies refine these findings—e.g., Wójcik, M., et al. show how LOY regulatory T cells intensify suppression in colorectal cancer, and Omilian, A., et al. highlight that CD163+ macrophages, surprisingly, can correlate with improved survival in triple-negative breast cancer.”

**Table 7:**
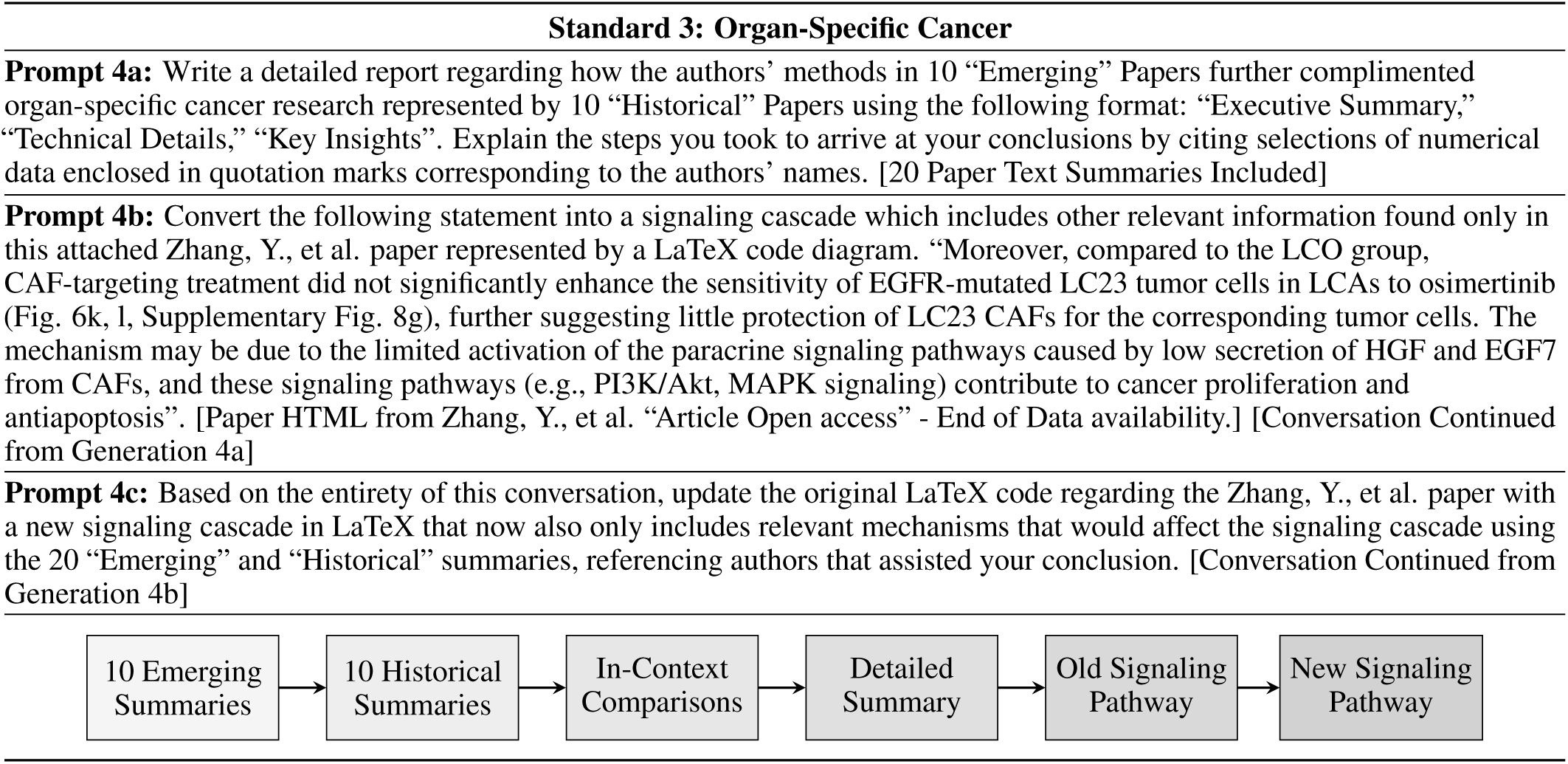
Standard 3 Process Diagram.

**Table 8:**
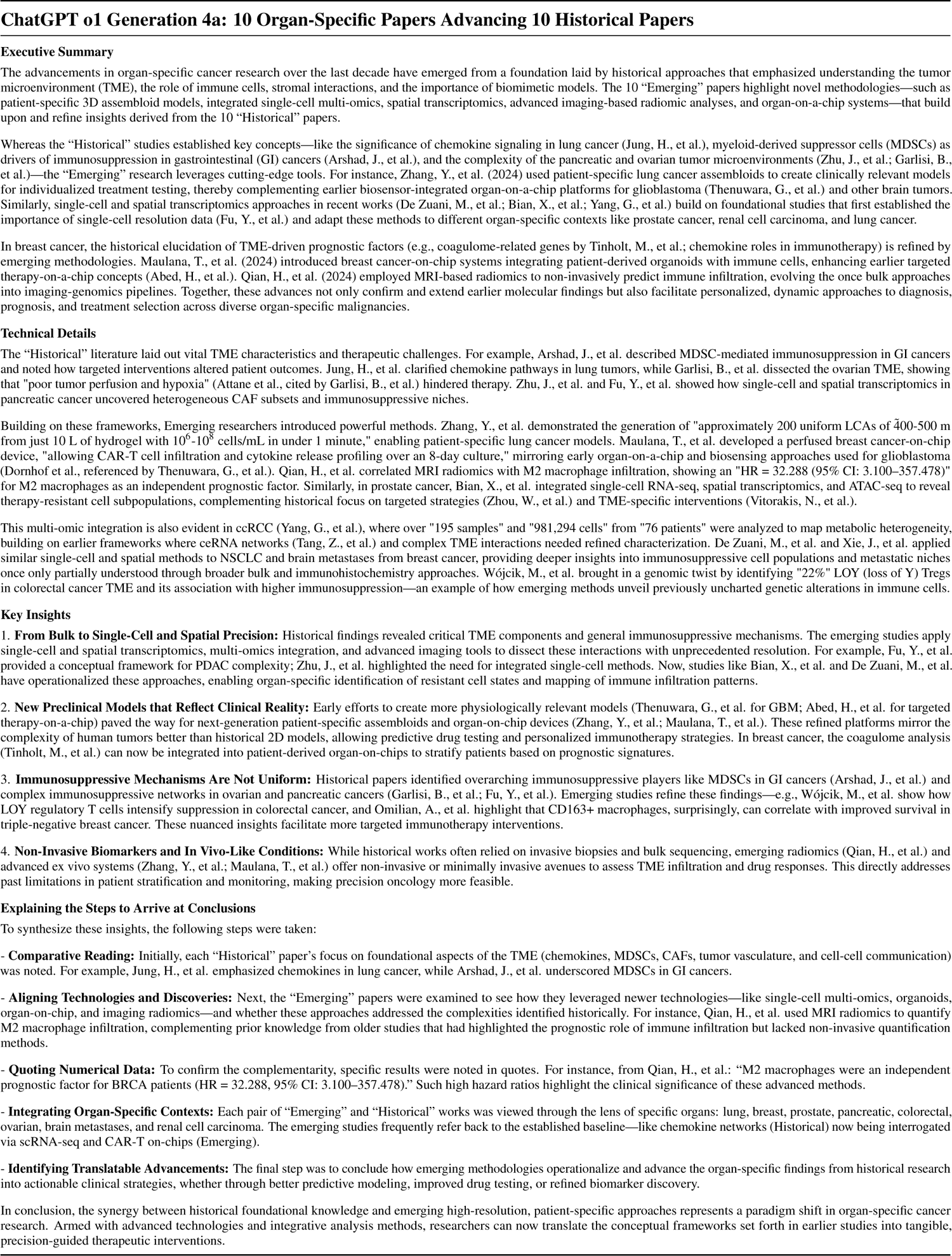
ChatGPT o1 Generation 4a: 1355 words, 68.85s.

The original “Old” 81 word signaling cascade from Zhang, Y., et al., 2024 shown in Prompt 4b was effectively coded in LaTeX, displayed natively, and scaled for readability, as the manuscript was also written in LaTeX Figure 5. The underlying mechanisms regarding the Zhang, Y., et al. excerpt in Prompt 4b appear to be properly depicted in the diagram, particularly from the sentence “The mechanism may be due to the limited activation of the paracrine signaling pathways caused by low secretion of HGF and EGF7 from CAFs, and these signaling pathways (e.g., PI3K/Akt, MAPK signaling) contribute to cancer proliferation and antiapoptosis”, however “Low HGF, Low FGF7” was mentioned twice. The overall meaning was retained as “Limited Paracrine Signaling” ultimately led to “Limited Proliferation & Antiapoptosis” as shown in the exerpt.

**Figure 5:**
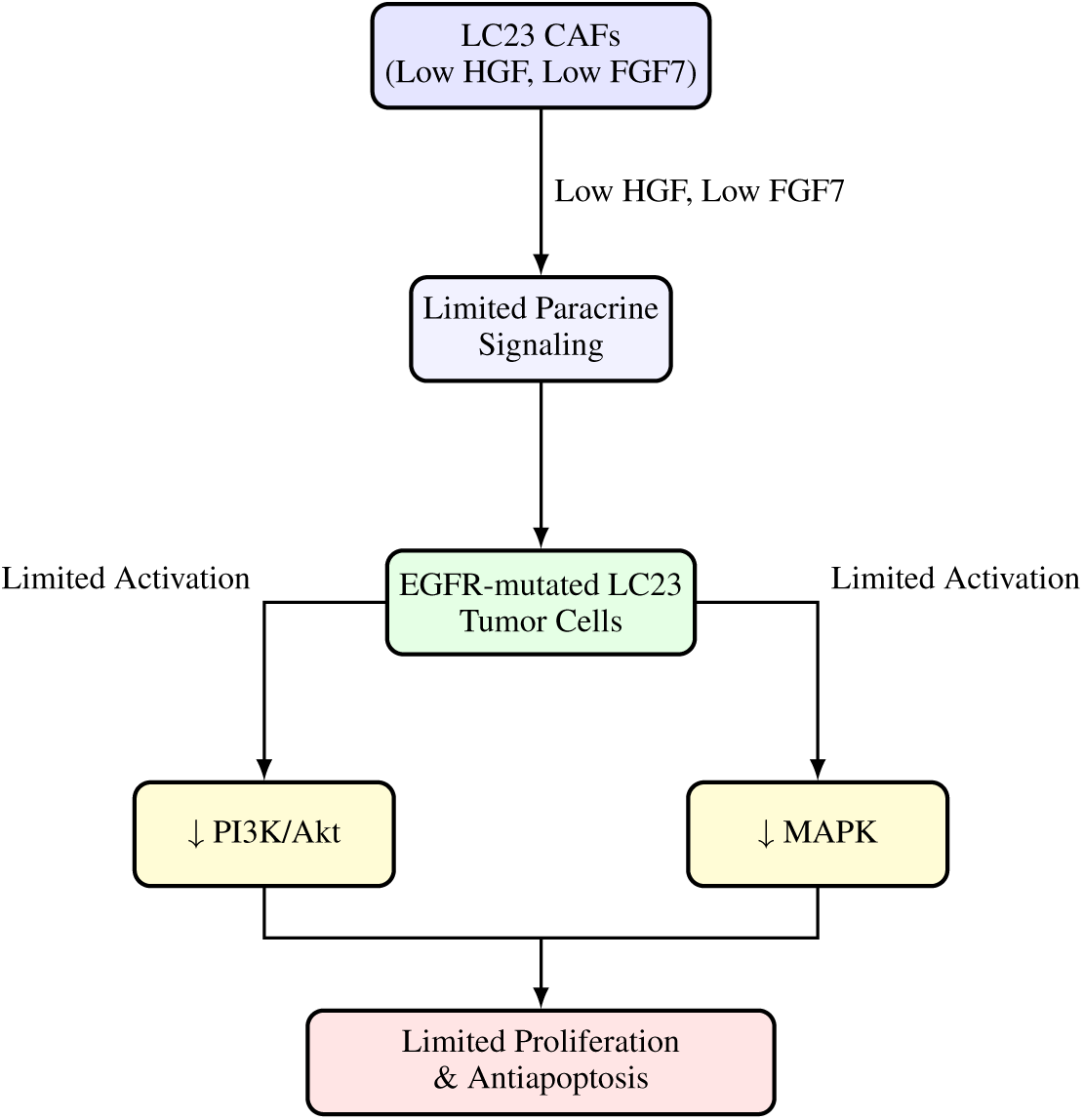
ChatGPT o1 Generation 4b Signaling Pathway

With this information now stored in the ChatGPT o1 conversation, the AI model was asked to update the original LaTeX code, this time with a new signaling pathway to include relevant mechanisms that could impact the signaling cascade using the 20 “Emerging” and “Historical” summaries, referencing authors that assisted the model’s conclusion are shown in Table 7 Prompt 4c. The resulting code is available in Supplementary SORp, which contains logical correlations between new information from authors in this study and the original cascade. However the LaTeX approach chosen by ChatGPT o1 did not display properly due to overcrowding of methods, authors, and mechanisms as shown in Figure 6. To more accurately depict the relationships from code generated by ChatGPT o1, a ChatGPT 4o conversation to manually position boxes and text yielded a new diagram, from which the manuscript author manually assigned positioning values shown in Figure 7.

**Figure 6:**
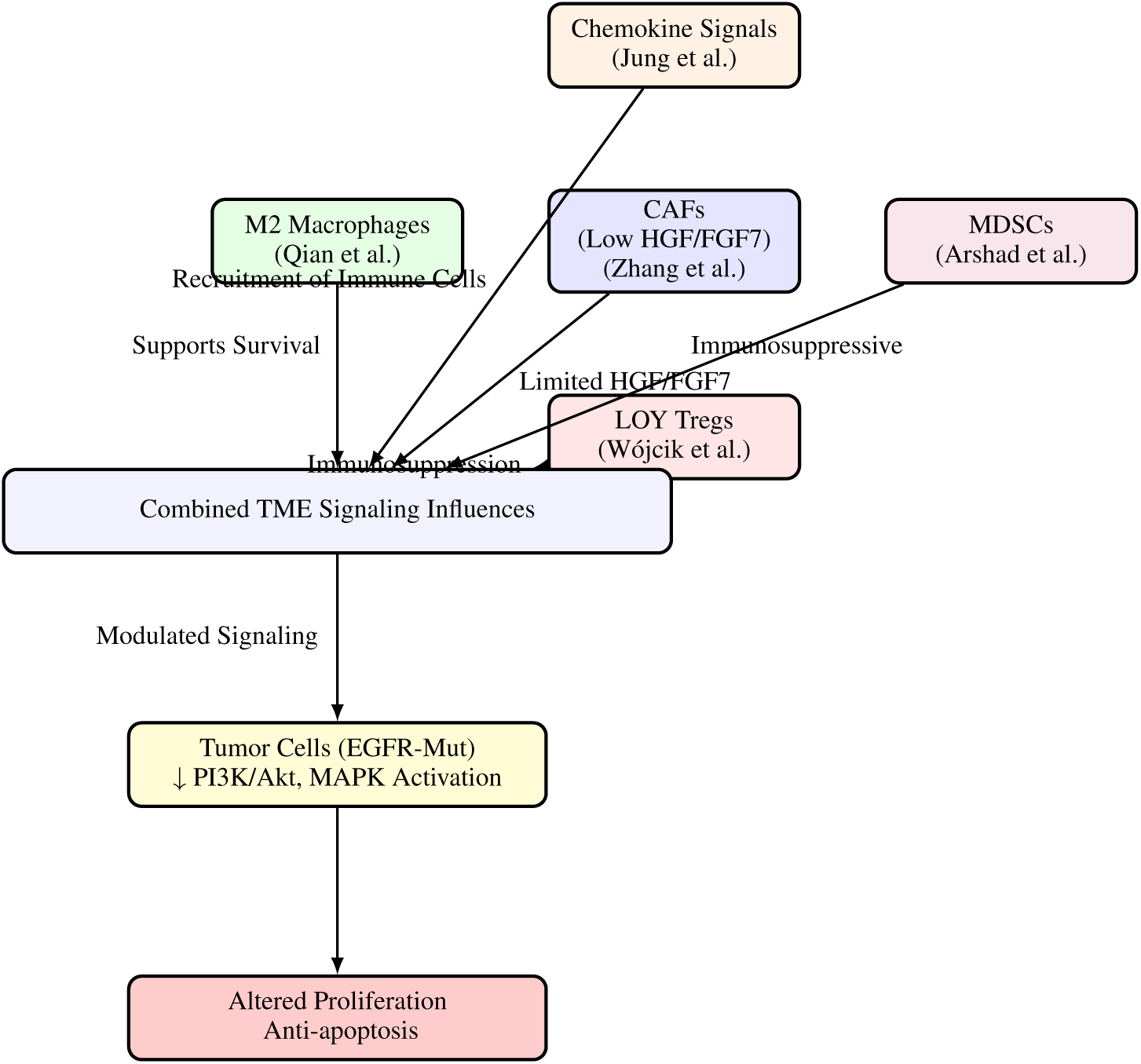
ChatGPT o1 Generation 4c Signaling Pathway

**Figure 7:**
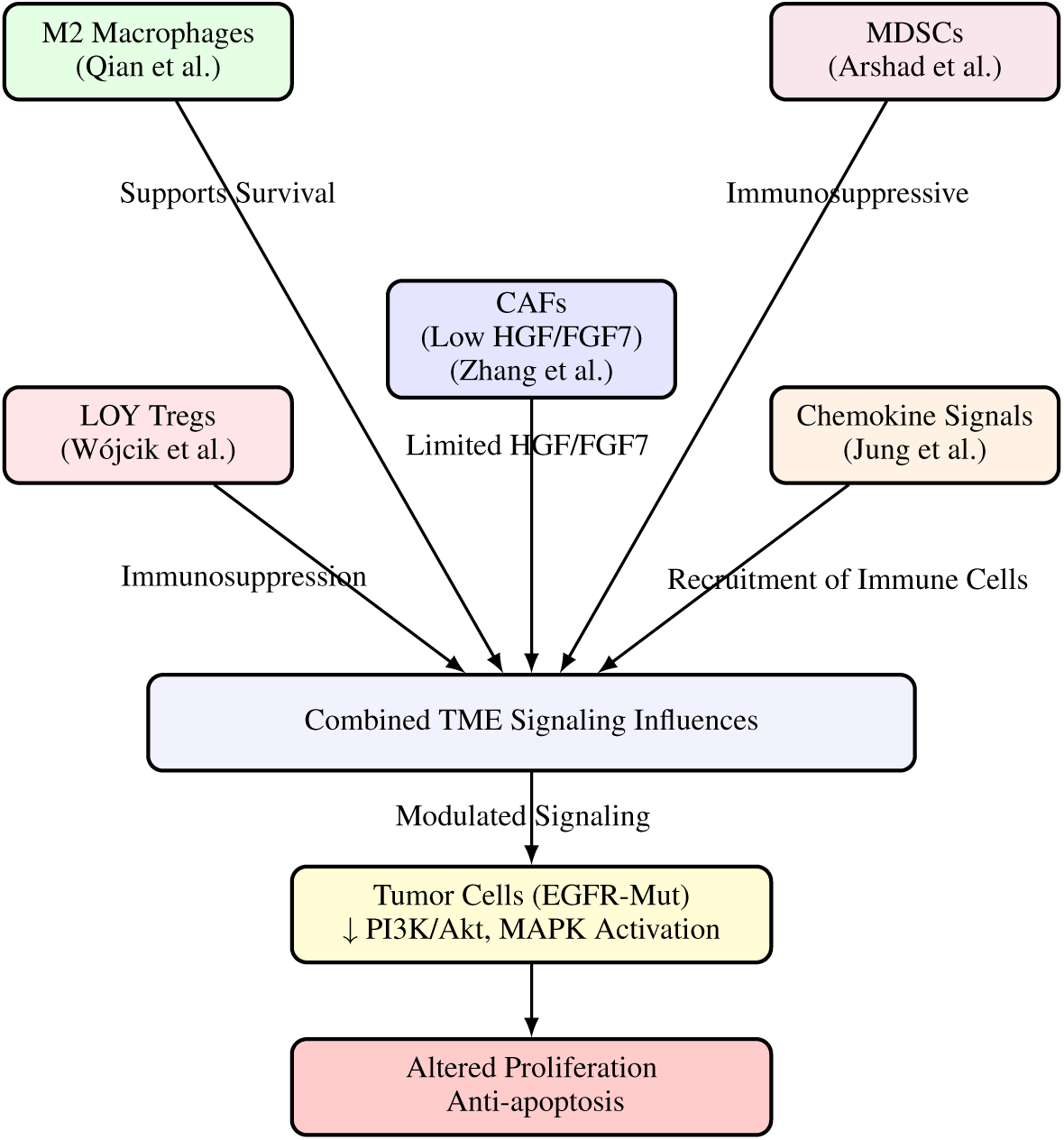
ChatGPT 4o, Author Repositioning from 4c

The overall meaning of the revised signaling pathway now leads to an “Altered Proliferation” “Anti-apoptosis” focal point with a similar mechanism from the Old pathway regarding Tumor Cells, PI3K/Akt, and MAPK, but now with authors’ new methods out of the 20 papers that were most applicable to this problem shown in Figure 7. Four of the authors depicted the specific area of research and the potential benefit provided as follows: Wójcik et al., LOY Tregs, Immunosuppression; Qian et al., M2 Macrophages, Supports Survival; Arshad et al., MDSCs, Immunosuppressive; and Jung et al., Chemokine Signals, Recruitment of Immune Cells. The original author of the mechanism, Zhang et al., maintained the CAFs/Low HGF/FGF7 relationship from the Old mechanism Figure 6. Therefore, ChatGPT o1 found that the end goal of “Altered Proliferation” “Anti-apoptosis” could be modified based on incorporating additional works regarding immunosuppression, survival longevity, and recruitment of immune cells.

### 5.2 Part III Discussion: Organ-Specific Cancer

Based on the three quotes in the Part III results section found in, the ability to convert a mechanism found in literature into LaTeX code representing a signaling pathway along with the incorporation of 20 paper summaries into a new coded diagram marked a major step towards the mainstream adoption of Conversation AI in drug discovery applications. Additional testing revealed that the quality of ChatGPT o1 generations were primarily only limited by the preciseness of author’s prompt, as triplicate studies for several key prompts were found to be reproducible shown in Supplementary SREP. Prompts 3b and 4c were more challenging to obtain effective results due to less ChatGPT o1 proficiency in for Section II in Python and Section III in LaTeX coding environments when compared to text.

53 citations representing the 20 paper summaries totaling 1,355 words were made by ChatGPT o1 in 68.85s. All authors in the summaries for Generation 4a were cited at least once, with others having two or more, with an average of 2.7 citations per author. The most cited author was 5 by Qian, H., et al. In addition, 2 authors mentioned by authors in original papers were also included: Attane et al., cited by Garlisi, B., et al., Dornhof et al., referenced by Thenuwara, G., et al.

In quoting numerical data, specific results were noted in quotes. For instance, from Qian, H., et al.: “M2 macrophages were an independent prognostic factor for BRCA patients (HR = 32.288, 95% CI: 3.100–357.478).” Such high hazard ratios highlight the clinical significance of these advanced methods. For each author citation, each fact was checked vs. summaries. Similarly, single-cell and spatial transcriptomics approaches in recent works (De Zuani, M., et al.; Bian, X., et al.; Yang, G., et al.) build on foundational studies that first established the importance of single-cell resolution data (Fu, Y., et al.) and adapt these methods to different organ-specific contexts like prostate cancer, renal cell carcinoma, and lung cancer.

Citations within citations: Maulana, T., et al. developed a perfused breast cancer-on-chip device, “allowing CAR-T cell infiltration and cytokine release profiling over an 8-day culture,” mirroring early organ-on-a-chip and biosensing approaches used for glioblastoma (Dornhof et al., referenced by Thenuwara, G., et al.). Qian, H., et al. correlated MRI radiomics with M2 macrophage infiltration, showing an “HR = 32.288 (95% CI: 3.100–357.478)” for M2 macrophages as an independent prognostic factor. An example of a LLM modification is: where over “195 samples” and “981,294 cells” from “76 patients” was actually exactly 195 samples.

## 6 Limitations and Future Work

a. Input words whose sum was less than the LLM context window, also referred to as context length, were processed effectively, but the potential negative of longer prompts has been an issue of debate [80, 81, 82]. Clau-3Opus has a limit of 200K tokens [34], while Lla3.1-405 is 128K tokens [35], and ChatGPT o1 using the “Pro” plan is 128K tokens [36], which is approximately 98K words. Based on these limits, neither the full 40 papers at 610K words or individual sets of 20 papers at 290K and 320K would come close to being accepted for processing. Therefore, only Clau-3Opus was used to process individual pdfs that were below the context window limit, while Lla3.1-405 was provided the 40 paper summaries at 17K words, and ChatGPT o1 was provided either a 8.8K word summary or 8.5K word summary. In each of the cases, context windows did not appear to pose an issue to quality, therefore future studies will likely increase the input provided to LLMs.
b. As of the time of publishing, there was no universal model available to accomplish each task in this paper with a single prompt. This was due to the fact that effective pdf processing of papers have only been identified in three models: Clau-3Opus, Clau-3.5So, and ChatGPT 4o, although HTML text can be copied into ChatGPT o1 less conveniently, followed by separate image uploads. Likewise, the ChatGPT o1 model significantly outperformed the other models’ output quality and length needed for advanced reasoning tasks in Section II, III. The Lla3.1-405 model was included due to the potential advantages of future models being open-source for industry, but the answer provided by the current model in this manuscript would likely be outdone, as seen in previous studies [83, 84]. With this being said, the high quality manual work of incorporating papers, combining summaries, and running the core experiments using code with visualizations may continue in the near-term to solve diseases.
c. Code generations in Python or LaTeX by the top ChatGPT o1 model required additional troubleshooting and typically lacked the ability to immediately convert its understandings of either a knowledge graph or signaling pathways to an attractive and convincing diagram. This was apparent in Part II Prompt 3b, where trial experiments yielded lower quality results. However, the model did properly incorporate features specified by the author. For Part III Prompt 4c, the relationships in code were appropriate, but ChatGPT o1 was not able to automatically set correct text and box positioning due to over-crowding, with further ChatGPT 4o and author modifications being required to properly display these relationships in Figure 7. Clau-3Opus also included quotations of some other phrases, and only some of the generations contained numerical text.

ChatGPT o1 quotes of passages from summaries also may include portions of surrounding text or modify a word to a different tense. Based on these results, certain aspects of general artificial intelligence using text only inputs is realistic, but would imply performing manual operations by the manuscript author in addition to generating effective final answers shown here. For instance, a two sentence prompt to obtain a breakthrough regarding cancer mechanisms would require several ChatGPT o1 instances in concert prior to generating the answer. This assumes no code or image incorporation, as processing these type of data typically requires additional testing.

## 7 Conclusions

Studies utilizing LLMs for cancer research have increased both in scope and quality of AI models. As few as 10 October 2024 articles have increased cancer research effectiveness, as apparent in recent publications such as Oh, Y., et al. published a LLM-driven oncology study in *Nature Communications* [29]. The emergence of reliable information retrieval methods present in models from Anthropic, high contextual awareness, and now advanced reasoning has opened the door for breakthroughs in cancer research by utilizing capabilities of different models combined with prompt engineering. Both immunology and organ-specific cancer are complex areas in medicine, as the systematic review of 2024 articles yielded 40 papers of over 600K words reduced to 17K words in order to run within the context length of ChatGPT o1.

Overall, the results obtained from Clau-3Opus in summarizing 40 papers were significant due to providing a consistent format and high accuracy for each summary as verified with multiple author search terms, but could have provided additional detailed numerical data quotations as specified in prompts. Lla3.1-405 provided an average detailed Top 10 Cancer Areas to research, lacking compelling evidence that was present using other models, however was the only model to process all 40 paper summaries at over 17,000 words. ChatGPT o1 provided groundbreaking and reproducible generations across two sets of 20 papers regarding advanced reasoning using text only. Additional sequential prompts in the same conversation incorporating Python or LaTeX code required additional development to achieve a professional knowledge map or signaling pathways.

The main takeaway of this work was that the quality and quantity of the data provided by Clau-3Opus to ChatGPT o1 in the form of reports was sufficient to create additional detailed reports and code used to visually represent numerous cancer research relationships. In the case of 20 summaries representing the tumor immune microenvironment, both a detailed report with an unprecedented number of relationships between author methods featured 37 citations, and a professional knowledge graph representing many of these relationships was produced in a two prompt conversation. For the 20 summaries representing organ-specific research, correlations between both author studies and specific cancer disease were made throughout many portions of the report, represented by 53 author citations. In addition, an 81 word Zhang et al. excerpt representing a cancer mechanism was converted by ChatGPT o1 into code correctly represented by a signaling pathway. Next, ChatGPT o1 updated the pathway based on the 20 summaries to yield additional code with useful relationships that now included a “Combined TME Signaling Influences” focal point with 4 additional authors, their specific area of cancer research and the benefit they could provide to the original pathway.

## Supporting information

Supplementary SIM

Supplementary SIMp

Supplementary SOR

Supplementary SORp

Supplementary SREP

## Appendix

**Figure 8:**
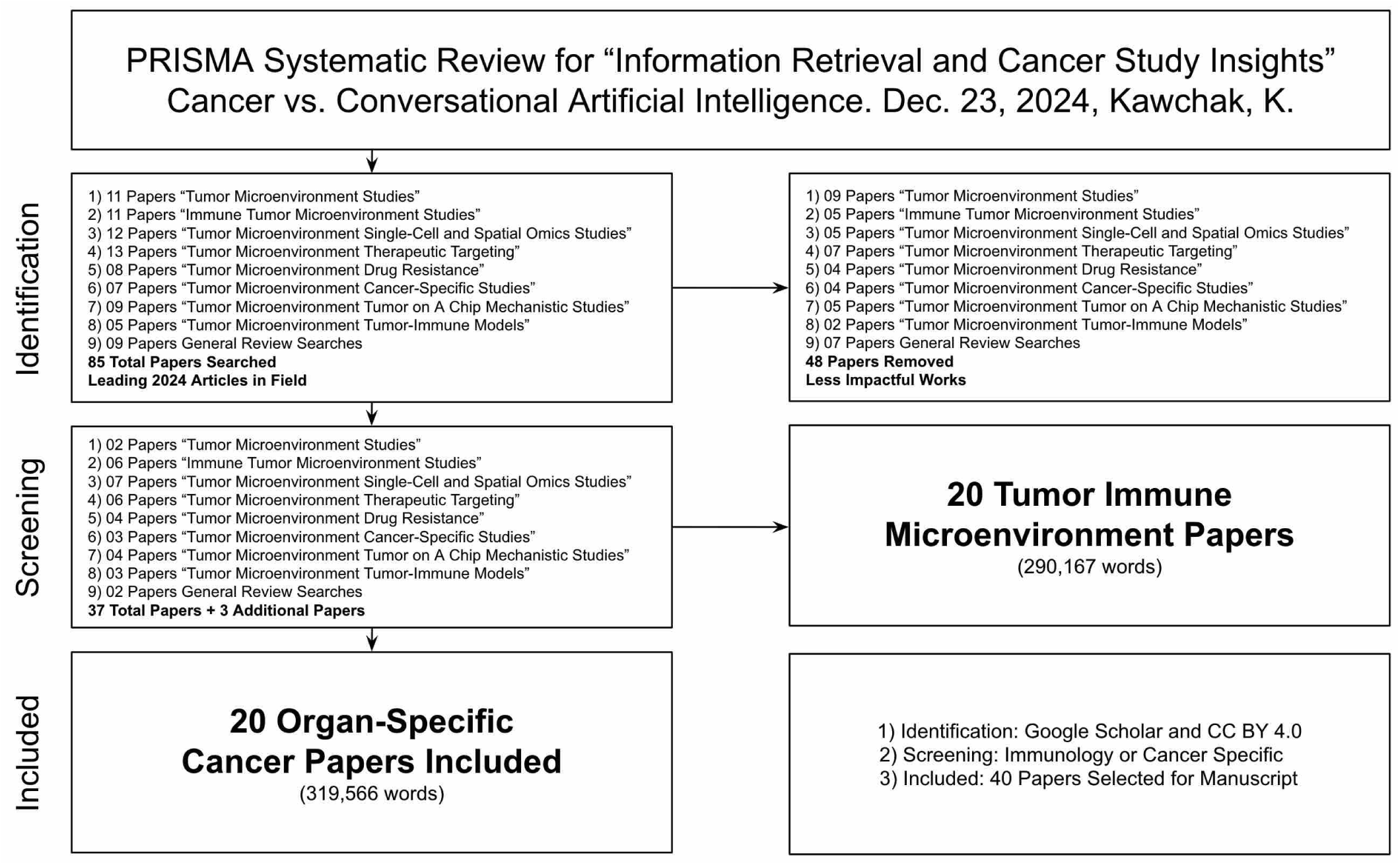
PRISMA Diagram for 40 Papers

