## Supplementary SIM for "Cancer vs. Conversational Artificial Intelligence"

### Supplementary Information SIM:

### 20 Paper Summaries

The content is available under CC BY 4.0. Refer to manuscript [Table 2] for paper references.

#### <u>Prompt 1</u>

#### Clau-3Opus: 20 Generations (Pages 01-35)

Total = 20 Generations Obtained December 7-8, 2024

**Prompt 1: (From "Write a detailed" - "the authors' names." on Page 1 were copied into LLM):**

#### <u>01 A pancreatic cancer organoid-in-matrix platform shows distinct sensitivities to T cell killing, 11803 words</u>

#### <u>Prompt 1: Write a detailed report based on the document regarding how the authors' methods advanced the tumor immune microenvironment field using the following format: "Executive Summary," "Technical Details," "Key Insights". Explain the steps you took to arrive at your conclusions by citing selections of numerical data enclosed in quotation marks corresponding to the authors' names. [1 Paper Attached]</u>

### **02 Exploring aryl hydrocarbon receptor expression and distribution in the tumor microenvironment, with a focus on immune cells, in various solid cancer types, 10012 words**

### **Prompt 1: Write a detailed report based on the document regarding how the authors' methods advanced the tumor immune microenvironment field using the following format: "Executive Summary," "Technical Details," "Key Insights". Explain the steps you took to arrive at your conclusions by citing selections of numerical data enclosed in quotation marks corresponding to the authors' names. [1 Paper Attached]**

### 03 INHBA(+) cancer-associated fibroblasts generate an immunosuppressive tumor microenvironment in ovarian cancer, 12050 words

### Prompt 1: Write a detailed report based on the document regarding how the authors' methods advanced the tumor immune microenvironment field using the following format: "Executive Summary," "Technical Details," "Key Insights". Explain the steps you took to arrive at your conclusions by citing selections of numerical data enclosed in quotation marks corresponding to the authors' names. [1 Paper Attached]

**04 Irreversible electroporation promotes a pro-inflammatory tumor microenvironment and anti-tumor immunity in a mouse pancreatic cancer model, 10174 words**
**Prompt 1: Write a detailed report based on the document regarding how the authors' methods advanced the tumor immune microenvironment field using the following format: "Executive Summary," "Technical Details," "Key Insights". Explain the steps you took to arrive at your conclusions by citing selections of numerical data enclosed in quotation marks corresponding to the authors' names. [1 Paper Attached]**
**Clau-3Opus: 43.06s, 462 words**

### **05 Loss of tumor suppressors promotes inflammatory tumor microenvironment and enhances LAG3+T cell mediated immune suppression, 12281 words**

### **Prompt 1: Write a detailed report based on the document regarding how the authors' methods advanced the tumor immune microenvironment field using the following format: "Executive Summary," "Technical Details," "Key Insights". Explain the steps you took to arrive at your conclusions by citing selections of numerical data enclosed in quotation marks corresponding to the authors' names. [1 Paper Attached]**

### **06 Molecular, Metabolic, and Subcellular Mapping of the Tumor Immune Microenvironment via 3D Targeted and Non-Targeted Multiplex Multi-Omics Analyses, 13386 words**

### **Prompt 1: Write a detailed report based on the document regarding how the authors' methods advanced the tumor immune microenvironment field using the following format: "Executive Summary," "Technical Details," "Key Insights". Explain the steps you took to arrive at your conclusions by citing selections of numerical data enclosed in quotation marks corresponding to the authors' names. [1 Paper Attached]**

**07 Revealing the role of SPP1+ macrophages in glioma prognosis and therapeutic targeting by investigating tumor-associated macrophage landscape in grade 2 and 3 gliomas, 15990 words**

**Prompt 1: Write a detailed report based on the document regarding how the authors' methods advanced the tumor immune microenvironment field using the following format: "Executive Summary," "Technical Details," "Key Insights". Explain the steps you took to arrive at your conclusions by citing selections of numerical data enclosed in quotation marks corresponding to the authors' names. [1 Paper Attached]**

### Prompt 1: Write a detailed report based on the document regarding how the authors' methods advanced the tumor immune microenvironment field using the following format: "Executive Summary," "Technical Details," "Key Insights". Explain the steps you took to arrive at your conclusions by citing selections of numerical data enclosed in quotation marks corresponding to the authors' names. [1 Paper Attached]

### 09 The Tumor Microenvironment Mediates the HIF-1α/PD-L1 Pathway to Promote Immune Escape in Colorectal Cancer, 8369 words

### Prompt 1: Write a detailed report based on the document regarding how the authors' methods advanced the tumor immune microenvironment field using the following format: "Executive Summary," "Technical Details," "Key Insights". Explain the steps you took to arrive at your conclusions by citing selections of numerical data enclosed in quotation marks corresponding to the authors' names. [1 Paper Attached]

### Clau-3Opus: 35.10s, 338 words

Executive Summary:

In this study, Sun et al. investigated the relationship between the HIF-1α/PD-L1 pathway and immune escape in colorectal cancer (CRC). They found that the hypoxic and inflammatory tumor microenvironment activates the HIF-1α/PD-L1 pathway to promote immune escape. Furthermore, they demonstrated that curcumin can inhibit this pathway, suggesting its potential as an immunomodulatory agent for CRC treatment.

Technical Details:

1. Bioinformatics analysis and Western blotting of clinical CRC samples showed HIF-1α and PD-L1 were overexpressed in CRC and positively correlated. "HIF-1α was positively correlated with PD-L1 in tumor samples ($r = 0.612$)" based on TCGA data.

2. In vitro hypoxia models using $CoCl_2$ and DFO revealed "stimulation of CT26 cells with $CoCl_2$ or DFO led to a concentration-dependent increase in HIF-1α and PD-L1 levels." The HIF-1α inhibitor YC-1 decreased PD-L1 expression in these models.

3. LPS was used to induce an inflammatory CRC microenvironment in CT26 cells. "10 μg/mL LPS was selected to build the inflammatory environment" which activated the HIF-1α/PD-L1 pathway. TLR4 inhibitor C34 suppressed HIF-1α/PD-L1 induction by LPS.

4. Curcumin inhibited proliferation, migration, and the TLR4/HIF-1α/PD-L1 pathway in the LPS-induced inflammatory model, based on MTT assay, Transwell assay, molecular docking, network pharmacology and Western blotting.

Key Insights:
1. Both hypoxic and inflammatory CRC microenvironments can activate the HIF-1α/PD-L1 pathway to enable tumor immune escape. Targeting this axis may enhance immunotherapy efficacy.

2. TLR4 positively regulates HIF-1α/PD-L1 in the inflammatory CRC setting. "TLR4, together with HIF-1α and PD-L1, was negatively expressed in adjacent normal tissue but positively expressed in tumor tissue" in clinical samples.

3. The anti-inflammatory phytochemical curcumin suppresses CRC progression by inhibiting the TLR4/HIF-1α/PD-L1 pathway, demonstrating promise as an immunomodulatory adjuvant therapy.

In summary, by employing a range of in vitro CRC models, bioinformatics, and clinical sample analyses, Sun et al. provided novel mechanistic insights into how the tumor microenvironment enables immune escape via HIF-1α/PD-L1. Their work suggests combination therapies targeting this pathway, potentially with curcumin, may improve outcomes for CRC patients.

### **10 Tumor immune dysfunction and exclusion subtypes in bladder cancer and pan-cancer- a novel molecular subtyping strategy and immunotherapeutic prediction model, 13502 words**

**Prompt 1: Write a detailed report based on the document regarding how the authors' methods advanced the tumor immune microenvironment field using the following format: "Executive Summary," "Technical Details," "Key Insights". Explain the steps you took to arrive at your conclusions by citing selections of numerical data enclosed in quotation marks corresponding to the authors' names. [1 Paper Attached]**

### 11 3D bioprinted tumor model a prompt and convenient platform for overcoming immunotherapy resistance by recapitulating the tumor microenvironment, 8226 words

### Prompt 1: Write a detailed report based on the document regarding how the authors' methods advanced the tumor immune microenvironment field using the following format: "Executive Summary," "Technical Details," "Key Insights". Explain the steps you took to arrive at your conclusions by citing selections of numerical data enclosed in quotation marks corresponding to the authors' names. [1 Paper Attached]

### 12 Deciphering the tumor immune microenvironment from a multidimensional omics perspective insight into next-generation CAR-T cell immunotherapy and beyond, 18217 words

### Prompt 1: Write a detailed report based on the document regarding how the authors' methods advanced the tumor immune microenvironment field using the following format: "Executive Summary," "Technical Details," "Key Insights". Explain the steps you took to arrive at your conclusions by citing selections of numerical data enclosed in quotation marks corresponding to the authors' names. [1 Paper Attached]

### 13 Exploring the dynamic interplay between exosomes and the immune tumor microenvironment- implications for breast cancer progression and therapeutic strategies, 10694 words

### Prompt 1: Write a detailed report based on the document regarding how the authors’ methods advanced the tumor immune microenvironment field using the following format: “Executive Summary,” “Technical Details,” “Key Insights”. Explain the steps you took to arrive at your conclusions by citing selections of

**numerical data enclosed in quotation marks corresponding to the authors' names. [1 Paper Attached]**

### Clau-3Opus: 38.55s, 421 words

Here is a report on how the methods in the reviewed paper advanced the tumor immune microenvironment field:

### 14 Metabolic reprogramming and immune evasion the interplay in the tumor microenvironment, 21561 words

**Prompt 1: Write a detailed report based on the document regarding how the authors' methods advanced the tumor immune microenvironment field using the following format: "Executive Summary," "Technical Details," "Key Insights". Explain the steps you took to arrive at your conclusions by citing selections of numerical data enclosed in quotation marks corresponding to the authors' names. [1 Paper Attached]**

### 15 Modulation of the tumor microenvironment and mechanism of immunotherapy-based drug resistance in breast cancer, 23551 words

### Prompt 1: Write a detailed report based on the document regarding how the authors' methods advanced the tumor immune microenvironment field using the following format: "Executive Summary," "Technical Details," "Key Insights". Explain the steps you took to arrive at your conclusions by citing selections of numerical data enclosed in quotation marks corresponding to the authors' names. [1 Paper Attached]

### **16 Myeloid-derived suppressor cells in cancer therapeutic targets to overcome tumor immune evasion, 18890 words**

### **Prompt 1: Write a detailed report based on the document regarding how the authors' methods advanced the tumor immune microenvironment field using the following format: "Executive Summary," "Technical Details," "Key Insights". Explain the steps you took to arrive at your conclusions by citing selections of numerical data enclosed in quotation marks corresponding to the authors' names. [1 Paper Attached]**

**17 Potential anti-tumor effects of regulatory T cells in the tumor microenvironment a review, 18241 words**

**Prompt 1: Write a detailed report based on the document regarding how the authors' methods advanced the tumor immune microenvironment field using the following format: "Executive Summary," "Technical Details," "Key Insights". Explain the steps you took to arrive at your conclusions by citing selections of numerical data enclosed in quotation marks corresponding to the authors' names. [1 Paper Attached]**

### 18 Progression of m6 A in the tumor microenvironment- hypoxia, immune and metabolic reprogramming, 21421 words

### Prompt 1: Write a detailed report based on the document regarding how the authors' methods advanced the tumor immune microenvironment field using the following format: "Executive Summary," "Technical Details," "Key Insights". Explain the steps you took to arrive at your conclusions by citing selections of numerical data enclosed in quotation marks corresponding to the authors' names. [1 Paper Attached]

**19 Regulatory T cells and immune escape in HCC understanding the tumor microenvironment and advancing CAR-T cell therapy, 14167 words**
**Prompt 1: Write a detailed report based on the document regarding how the authors' methods advanced the tumor immune microenvironment field using the following format: "Executive Summary," "Technical Details," "Key Insights". Explain the steps you took to arrive at your conclusions by citing selections of numerical data enclosed in quotation marks corresponding to the authors' names. [1 Paper Attached]**
**Clau-3Opus: 38.02s, 412 words**

### 20 Single-Cell Informatics for Tumor Microenvironment and Immunotherapy, 13727 words

### Prompt 1: Write a detailed report based on the document regarding how the authors' methods advanced the tumor immune microenvironment field using the following format: "Executive Summary," "Technical Details," "Key Insights". Explain the steps you took to arrive at your conclusions by citing selections of numerical data enclosed in quotation marks corresponding to the authors' names. [1 Paper Attached]
