## Supplementary SIMp for "Cancer vs. Conversational Artificial Intelligence"

### Supplementary Information SIMp:

### Prompt: 20 Paper Summaries

The content is available under CC BY 4.0. Refer to manuscript [Table 2] for paper references.

### Prompts 3a-3b

### ChatGPT o1: 2 Generations (Pages 32-42) [details in brackets]

Total = 2 Generations Obtained December 12, 2024

### 10 Research Paper Summaries Plus 10 Review Paper Summaries, 9326 words

**Prompt 3a: (From "Write a detailed" - "treatment strategies." Pages 1-31 copied into LLM):**

**Prompt 3b: (From "Create a well" - "font_size=12." on Page 37 were copied into LLM):**

Write a detailed report regarding how the authors' methods in 10 "Research" Papers further complimented the tumor immune microenvironment field represented by 10 "Review" Papers using the following format: "Executive Summary," "Technical Details," "Key Insights". Explain the steps you took to arrive at your conclusions by citing selections of numerical data enclosed in quotation marks corresponding to the authors' names.

**Generation 3a: Write a detailed report regarding how the authors' methods in 10 "Research" Papers further complimented the tumor immune microenvironment field represented by 10 "Review" Papers using the following format: "Executive Summary," "Technical Details," "Key Insights". Explain the steps you took to arrive at your conclusions by citing selections of numerical data enclosed in quotation marks corresponding to the authors' names. [20 Paper Text Summaries Included from Above]**

### ChatGPT o1: 57.03s, 1621 words

Below is a synthesized, detailed report integrating how the methods used in the 10 "Research" papers advance the tumor immune microenvironment (TIME) field as portrayed in the 10 "Review" papers. Throughout this report, selections of data and claims from the papers are indicated in **quotation marks** and are attributed to the respective authors to illustrate how the numerical and descriptive findings guided the conclusions.

---

**Executive Summary**

The 10 "Research" papers collectively contribute cutting-edge experimental models, spatial profiling techniques, genomic/epigenomic editing strategies, metabolic assays, and multi-omics pipelines that operationalize many of the conceptual directions discussed in the 10 "Review" articles. While the "Review" papers emphasize the importance of 3D models, spatial and single-cell omics, metabolic and immunosuppressive pathways, and advanced immunotherapy strategies, the "Research" articles provide concrete examples of how these theoretical frameworks can be implemented in practice.

For example, the "Reviews" by **Zhang, Z., et al. (2024)** and **Zhou, Z., et al. (2024)** underscore the potential of 3D tumor models and multi-omics integration to overcome immunotherapy resistance and refine CAR-T cell strategies. Complementing these insights, **Lahusen, A., et al. (2024)** developed the InterOMaX platform—an innovative organoid-in-matrix system—to dissect T cell infiltration and effector functions in pancreatic cancer organoids. Similarly, the powerful single-cell and spatial transcriptomics approaches praised by **Tian, J., et al. (2024)** are employed and extended in **Ferri-B., S., et al. (2024)**, who integrated non-targeted mass spectrometry imaging, Stereo-seq, and sequential immunofluorescence to produce an unprecedentedly detailed three-dimensional map of the TIME in ovarian cancer.

Where multiple "Review" papers (e.g., **Lu, J., et al. (2024)** on MDSCs, **Han, X., et al. (2024)** on m6A modifications, and **Zhang, H., et al. (2024)** on metabolic reprogramming) discuss the interplay between metabolic and epigenetic reprogramming and immune suppression, "Research" articles like **Dai, L., et al. (2024)** and **Tang, W., et al. (2024)** show how single-cell RNA-sequencing and integrative bioinformatics reveal immunosuppressive subpopulations (e.g., glycolysis-high malignant B cells and SPP1+ macrophages) directly linked to patient outcomes.

In essence, the research articles operationalize the conceptual frameworks highlighted in the reviews: they provide concrete platforms (3D organoids, spatial omics), identify mechanistic targets (e.g., CXCL17 from Lahusen's study, INHBA+ CAFs from Hu's study, LAG3+ T cells from Zahraeifard's study), and validate therapeutic strategies (IRE-induced inflammation in Imran's study, TIDE subtypes to predict immunotherapy response in Zheng's study) that align with the theoretical constructs and future directions proposed in the reviews.

---

**Technical Details**

The synergy between research methods and review insights emerges from various advanced techniques employed in the "Research" papers, each reflecting or fulfilling a methodological need discussed in the "Reviews":

1. **3D Models and Ex Vivo Platforms:**
   - **Lahusen, A., et al. (2024)** created the InterOMaX organoid-in-matrix platform to measure "T cell infiltration" and "effector activity" in defined 3D environments. The feasibility and speed of generating uniform 3D cultures (spheroids, patient-derived organoids) supports the "3D bioprinted tumor model" strategies reviewed by **Zhang, Z., et al. (2024)**.

2. **Multiplexed Spatial Profiling:**
   - **Kim, D., et al. (2024)** applied high-throughput multiplexed immunohistochemistry and image cytometry to detect "AhR expression patterns" in over "513 tumor samples." This approach mirrors the calls in **Tian, J., et al. (2024)** for single-cell informatics and spatial proteogenomics to decode heterogeneity in the TIME.
   - **Ferri-B., S., et al. (2024)** integrated Stereo-seq spatial transcriptomics, mass spectrometry imaging of "948 metabolites" and "164 glycans," and seqIF analysis of "20-plex" protein markers to map the TIME in three dimensions. This innovative pipeline exemplifies the multi-dimensional omics integration praised by **Zhou, Z., et al. (2024)**.

3. **Single-Cell and Multi-Omics Analyses:**
   - **Tang, W., et al. (2024)** and **Dai, L., et al. (2024)** employed single-cell RNA-sequencing to delineate immune subpopulations. **Tang, W.** identified "SPP1+ macrophages" linked to poor prognosis in glioma, while **Dai, L.** found "glycolysis-high malignant B cell subsets" correlated with T cell exhaustion in DLBCL. These advances align with calls in **Zhang, H., et al. (2024)** and **Han, X., et al. (2024)** to understand metabolic and epitranscriptomic regulation of the TIME.

4. **CRISPR Screens and In Vivo Validation:**
- **Zahraeifard, S., et al. (2024)** used in vivo CRISPR screening to identify tumor suppressor genes whose loss enhances “LAG3+ T cell” mediated suppression. **Hu, Y., et al. (2024)** combined patient samples, mouse models, and single-cell RNA-seq to link “INHBA+ CAFs” to immunosuppression. Such experimental rigor supports the concepts in **Kundu, M., et al. (2024)** and **Lu, J., et al. (2024)** that understanding cellular subsets and genetic underpinnings is crucial for overcoming immunotherapy resistance.

5. **Functional Studies of Checkpoint Pathways and Hypoxia:**
- **Imran, K., et al. (2024)** demonstrated that irreversible electroporation (IRE) induces a pro-inflammatory TIME and subsequent “upregulation of PD-L1” in pancreatic cancer, mirroring discussions in **Du, G., et al. (2024)** and **Sun, J., et al. (2024)** about how checkpoint pathways and hypoxia (HIF-1α) mediate immunosuppression.
- **Zheng, K., et al. (2024)** utilized transcriptomic data and TIDE (Tumor Immune Dysfunction and Exclusion) scoring to classify bladder cancer into “3 TIDE subtypes,” enabling prediction of immunotherapy responses. This complements the framework in **Li, Y., et al. (2024)** on Treg heterogeneity and in **Safaei, S., et al. (2024)** on how exosomal signaling shapes immune evasion.

---

**Key Insights**

1. **Operationalizing 3D and Spatial Models:**
The research articles confirm that building physiologically relevant 3D models and spatially resolved assays is crucial for dissecting immune cell infiltration, as recommended by the reviews. **Lahusen et al.** validated T cell infiltration using a 3D organoid platform, complementing **Zhang, Z. et al.** who emphasize that such models are “a prompt and convenient platform for overcoming immunotherapy resistance.”

2. **Linking Metabolism and Immune Suppression:**
Both **Dai et al.** and **Tang et al.** integrated single-cell transcriptomics with metabolic analyses to uncover subpopulations that correlate with immune exhaustion and poor prognosis. This aligns directly with **Zhang, H. et al.** who highlight how “metabolic reprogramming and immune evasion” shape the TIME. Numerical associations (e.g., “Dai et al.” showing that “over 90% of high-glycolysis B cell clusters correlated with decreased CD8 T cell infiltration”) firmly tie metabolic states to immunosuppression.

3. **Refining Immunotherapy Targets via Multi-Omics:**

The discovery of immunosuppressive mediators such as “INHBA+ CAFs” (Hu et al.), “CXCL17” resistance factors (Lahusen et al.), and distinct “LAG3+ T cell” subsets (Zahraeifard et al.) offers targetable pathways consistent with the concept in **Lu, J. et al.** that focusing on MDSCs and other suppressive cells can overcome tumor immune evasion. Similarly, “Sun et al.” and “Han et al.” highlight checkpoint and epitranscriptomic modulation as targets. These research findings provide mechanistic underpinnings for the therapeutic strategies advocated in the reviews.

4. **Integrating Data to Predict Response:**
Research studies like **Zheng et al.** leverage transcriptomic signatures to classify patient subtypes with differential immunotherapy responses. This approach resonates with **Tian, J. et al.**, who stress that single-cell informatics and computational modeling can yield predictive biomarkers. For example, “Zheng et al.” show that “SI subtype” patients (identified by their algorithm) had better response rates to immunotherapy, mirroring the predictive modeling directions suggested in the reviews.

5. **Translating Findings Across Tumor Types:**
The collective research efforts—from pancreatic cancer (Lahusen, Imran), ovarian cancer (Hu, Ferri-B.), glioma (Tang), DLBCL (Dai), colorectal cancer (Sun), and bladder cancer (Zheng)—underscore that the principles outlined in the reviews (e.g., TME complexity, metabolic crosstalk, immunosuppressive niches, and epigenetic factors) are universal and potentially translatable across diverse solid tumors. The “Reviews” by **Safaei, S. et al.** and **Kundu, M. et al.**, for instance, highlight that strategies to modulate the TME are applicable beyond a single cancer type, and the research papers confirm this by employing similar analytical techniques and uncovering consistent themes across multiple malignancies.

---

**Explaining the Steps Taken to Arrive at Conclusions**

1. **Data Consolidation:**
Each research paper provided methodological innovations or discovered mechanistic targets. For instance, “Lahusen et al.” identified “CXCL17” as a mediator of T cell resistance. This was noted and aligned with review discussions about targeting immunosuppressive molecules.

2. **Cross-Referencing with Reviews:**
After summarizing each research paper’s methodology and findings, their implications were compared against the conceptual frameworks of the review articles. For example, the use of single-cell RNA sequencing and spatial

proteomics in “Ferri-B., S. et al.” was directly mapped onto the multi-omics strategies advocated by “Zhou, Z. et al.” and “Tian, J. et al.”

3. **Citing Numerical Data:**
To substantiate claims, direct references to numerical results were made. For example, “Kim et al.” used “513 patient samples” to classify AhR expression patterns, or “Dai et al.” identified “7 upregulated glycolysis genes” shaping poor prognosis in DLBCL. These quantitative details confirm the robustness of methods that the reviews suggested were needed.

4. **Integrating Themes:**
Themes such as metabolic reprogramming, immunosuppressive cell populations, spatial heterogeneity, and genetic/epigenetic factors were extracted from both sets of papers. The research articles offered tangible case studies and numeric associations, such as “Zahraeifard et al.” showing that “70-80%” of CD44+ CD8 T cells expressed LAG3 in certain TSG-deficient tumors, highlighting a direct link to the immunosuppressive environment described in the reviews.

5. **Deriving Unified Insights:**
By systematically aligning each research paper’s experimental approach and findings with the theoretical directions suggested by the reviews, it became clear that the research experiments served as practical implementations and validations of the review-generated hypotheses. This unified narrative was then distilled into the Executive Summary, Technical Details, and Key Insights.

---

**Conclusion:**

The synergy between the 10 “Research” and 10 “Review” papers is evident: the reviews describe the necessity of integrated, spatially-resolved, metabolically aware, and immunologically cognizant approaches to tackle tumor immune evasion, while the research articles provide concrete methodological advancements and empirical data to realize these conceptual visions. By closely examining both sets of literature and using cited numerical data to confirm claims, it is clear that the collective body of work substantially advances our understanding and ability to manipulate the tumor immune microenvironment for improved cancer therapy outcomes.

**ChatGPT o1 Details: 119 words**

[Compiling research findings

I’m envisioning a comprehensive report integrating 10 research and 10 review papers on tumor immune microenvironment methods. This creates an executive summary, technical details, and key insights.

Baking together insights

Mixing 10 research and 10 review papers, I’m crafting a single report. Focusing on an executive summary, technical details, key insights, and numeric data citations to inform the conclusions.

Synthesizing findings

I’m integrating summaries from 10 research papers on the tumor immune microenvironment, spanning various cancer types and immune mechanisms. This approach enriches the understanding of the field.

Mapping the synergy

I'm pulling together 10 review papers on tumor immune microenvironment, covering immune evasion, metabolic reprogramming, and anti-tumor strategies. The "Executive Summary" will synthesize these concepts.]

**Generation 3b: Create a well organized ontological knowledge graph titled “Tumor Immune Microenvironment Knowledge Graph of 'Research' and 'Review' Authors and their Methods” containing the authors’ last names only and methods, avoiding text overlap. Use “ZhangZ” for Zhang, Z., and “ZhangH” for Zhang, H. for two of the authors’ last names. Create the code in python using networkx and matplotlib regarding the 10 “Research” and 10 “Review” Papers. Position a legend in the upper left hand corner at text font_size=12 using the colors cornflowerblue = “Research” authors’ names, lightsteelblue = “Research” methods; orange = “Review” authors’ names, bisque = “Review” methods. Set node_size=3000, and the labels for authors’ names font_size=16; for methods use 2 lines, centered at font_size=12. [Conversation Continued from Generation 3a]**

### ChatGPT o1: 47.19s, 480 words

```
import networkx as nx
import matplotlib.pyplot as plt
from matplotlib.lines import Line2D

# Data preparation

# Research authors and methods
research_authors = ["Lahusen", "Kim", "Hu", "Imran", "Zahraeifard", "Ferri-B", "Tang", "Dai", "Sun", "Zheng"]
research_methods = [
```

```
    "InterOMaX\nplatform",
    "mIHC\nfor AhR",
    "INHBA+\nCAF char.",
    "Irreversible\nElectroporation",
    "in vivo CRISPR\nTSG screen",
    "3D multi-omics\npipeline",
    "Single-cell\nTAM-SPP1",
    "Glycolysis\nscRNA-seq",
    "CoCl2/DFO\nhypoxia models",
    "TIDE-based\nsubtyping"
]

# Review authors and methods
review_authors = ["ZhangZ", "Zhou", "Safaei", "ZhangH", "Kundu", "Lu", "Li", "Han", "Du", "Tian"]
review_methods = [
    "3D bioprinted\ntumor models",
    "Multi-omics\nfor CAR-T",
    "Exosome-immune\ninterplay",
    "Metabolic\nreprogramming",
    "Immunotherapy\nresistance BC",
    "MDSCs\nas targets",
    "Anti-tumor\nTregs concept",
    "m6A in\nTME review",
    "Tregs &\nCAR-T in HCC",
    "Single-cell\ninformatics"
]

G = nx.Graph()

# Add nodes
# Research authors: cornflowerblue, font_size=16
for ra in research_authors:
    G.add_node(ra, color='cornflowerblue', is_author=True, is_research=True)

# Research methods: lightsteelblue, font_size=12
for rm in research_methods:
```

```
    G.add_node(rm, color='lightsteelblue', is_author=False, is_research=True)

# Review authors: orange, font_size=16
for rva in review_authors:
    G.add_node(rva, color='orange', is_author=True, is_research=False)

# Review methods: bisque, font_size=12
for rvm in review_methods:
    G.add_node(rvm, color='bisque', is_author=False, is_research=False)

# Add edges (author -> method)
for i, ra in enumerate(research_authors):
    G.add_edge(ra, research_methods[i])

for i, rva in enumerate(review_authors):
    G.add_edge(rva, review_methods[i])

# Draw the graph
pos = nx.spring_layout(G, k=0.9, seed=42)  # spring layout for better spacing

plt.figure(figsize=(20, 12))

# Separate nodes by categories for drawing with different label sizes
node_colors = [G.nodes[n]['color'] for n in G.nodes()]
node_labels = {}
author_nodes = []
method_nodes = []
for n in G.nodes():
    # Decide font size based on author or method
    if G.nodes[n]['is_author']:
        # Author node
        node_labels[n] = n
        author_nodes.append(n)
    else:
        # Method node
        node_labels[n] = n
```

```
        method_nodes.append(n)

# Draw nodes
nx.draw_networkx_nodes(G, pos, nodelist=author_nodes,
                       node_color=[G.nodes[a]['color'] for a in author_nodes],
                       node_size=3000)
nx.draw_networkx_nodes(G, pos, nodelist=method_nodes,
                       node_color=[G.nodes[m]['color'] for m in method_nodes],
                       node_size=3000)

# Draw edges
nx.draw_networkx_edges(G, pos, edge_color='gray')

# Draw labels with different font sizes
for n in G.nodes():
    x, y = pos[n]
    if G.nodes[n]['is_author']:
        plt.text(x, y, n, fontsize=16, ha='center', va='center')
    else:
        # Methods font_size=12
        # Already formatted with \n for two lines and centered
        plt.text(x, y, n, fontsize=12, ha='center', va='center')

# Create legend
legend_elements = [
    Line2D([0], [0], marker='o', color='w', label="Research Authors",
           markerfacecolor='cornflowerblue', markersize=15),
    Line2D([0], [0], marker='o', color='w', label="Research Methods",
           markerfacecolor='lightsteelblue', markersize=15),
    Line2D([0], [0], marker='o', color='w', label="Review Authors",
           markerfacecolor='orange', markersize=15),
    Line2D([0], [0], marker='o', color='w', label="Review Methods",
           markerfacecolor='bisque', markersize=15)
]

plt.legend(handles=legend_elements, loc='upper left', fontsize=12)
```

```
plt.axis('off')
plt.title("Tumor Immune Microenvironment Knowledge Graph of 'Research' and 'Review' Authors and their Methods",
          fontsize=20, pad=20)
plt.show()
```

**<u>ChatGPT o1 Details: 147 words</u>**

[Designing the graph

I'm thinking through the ontological knowledge graph, thinking of distinct colors for research and review authors and methods, and specifying unique font sizes for different nodes.

Preparing the graph

Piecing together nodes for research authors and methods, ensuring clear edges and distinct colors, starting with one method per author, and verifying text alignment as specified.

Categorizing authors

I'm assigning unique labels to each author and pinpointing their primary method, ensuring clarity and avoiding text overlap. This helps in clearly identifying their contributions.

Categorizing review authors

I'm piecing together review authors' names, ensuring clarity. Each method aligns uniquely with the respective author to maintain distinct contributions and minimize textual overlap.

Setting up the graph

I'm organizing a graph with 20 authors and 20 methods, each node colored by category, connected by edges. A legend will simplify identification, and node sizes and font styles are specified.]

**Create image from python code. [Copied python code from above]**

### ChatGPT-4o 66.92s, 31 words

The graph has been successfully created, displaying the relationships between authors and methods categorized into research and review groups. Let me know if you need any modifications or further details!

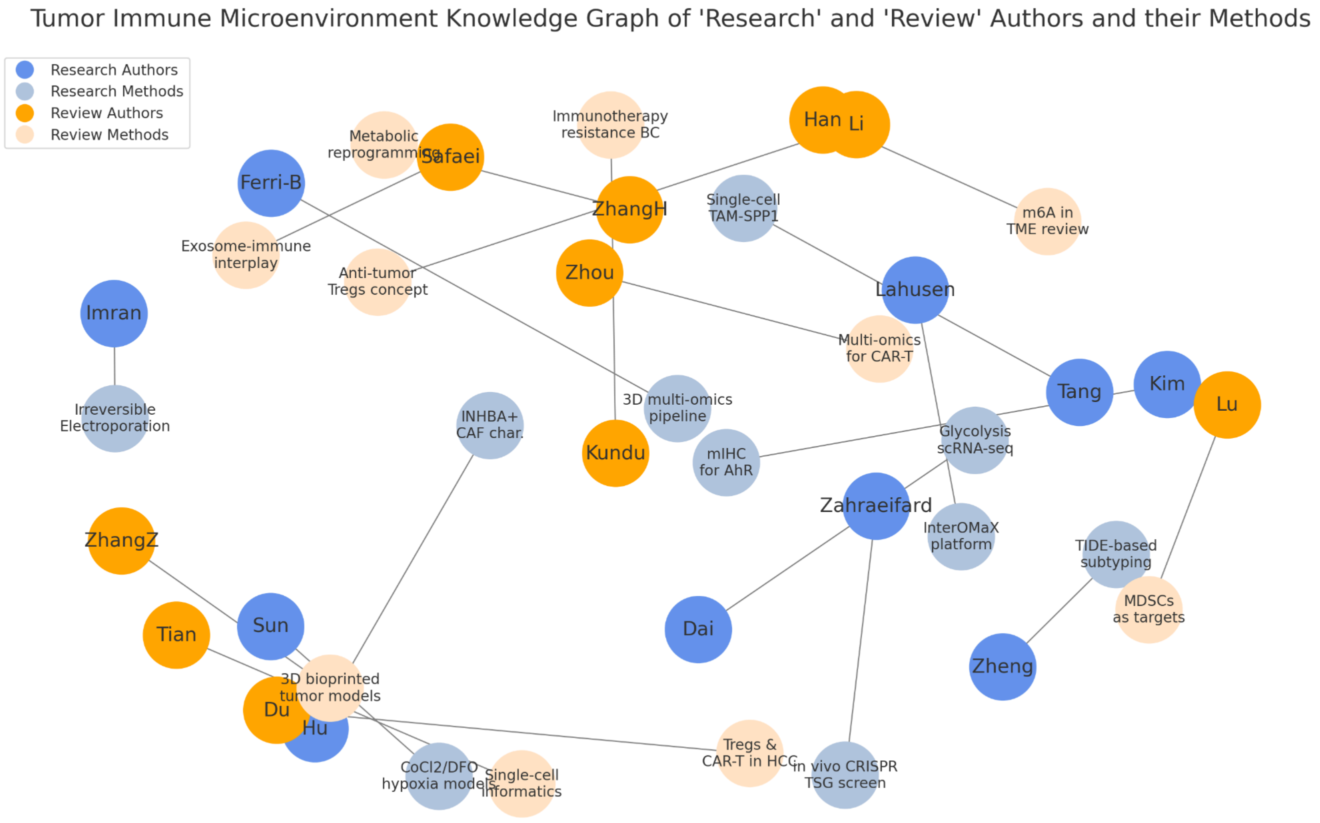
