## Supplementary SOR for "Cancer vs. Conversational Artificial Intelligence"

### Supplementary Information SOR:

#### 20 Paper Summaries

The content is available under CC BY 4.0. Refer to manuscript [Table 2] for paper references.

#### Prompt 2a

#### Clau-3Opus: 20 Generations (Pages 01-33)

Total = 20 Generations Obtained December 7, 2024

**Prompt 2a: (From "Write a detailed" - "the authors' names." on Page 1 copied into LLM):**

#### 21 A patient-specific lung cancer assembloid model with heterogeneous tumor microenvironments, 13684 words

#### Prompt 2a: Write a detailed report based on the document regarding how the authors' methods advanced organ-specific cancer research using the following format: "Executive Summary," "Technical Details," "Key Insights". Explain the steps you took to arrive at your conclusions by citing selections of numerical data enclosed in quotation marks corresponding to the authors' names. [1 Paper Attached]

### 25 Integration Analysis of Single-Cell Multi-Omics Reveals Prostate Cancer Heterogeneity, 15940 words

### Prompt 2a: Write a detailed report based on the document regarding how the authors' methods advanced organ-specific cancer research using the following format: "Executive Summary," "Technical Details," "Key Insights". Explain the steps you took to arrive at your conclusions by citing selections of numerical data

**26 Loss of Y in regulatory T lymphocytes in the tumor micro-environment of primary colorectal cancers and liver metastases, 9633 words**
**Prompt 2a: Write a detailed report based on the document regarding how the authors' methods advanced organ-specific cancer research using the following format: "Executive Summary," "Technical Details," "Key Insights". Explain the steps you took to arrive at your conclusions by citing selections of numerical data enclosed in quotation marks corresponding to the authors' names. [1 Paper Attached]**
**Clau-3Opus: 51.52s, 596 words**

### **28 Single-cell and spatial transcriptomics analysis of non-small cell lung cancer, 16852 words**
### **Prompt 2a: Write a detailed report based on the document regarding how the authors' methods advanced organ-specific cancer research using the following format: "Executive Summary," "Technical Details," "Key Insights". Explain the steps you took to arrive at your conclusions by citing selections of numerical data enclosed in quotation marks corresponding to the authors' names. [1 Paper Attached]**
### **Clau-3Opus: 41.65s, 357 words**

**29 Single-cell RNA sequencing elucidated the landscape of breast cancer brain metastases and identified ILF2 as a potential therapeutic target, 10994 words**
**Prompt 2a: Write a detailed report based on the document regarding how the authors' methods advanced organ-specific cancer research using the following format: "Executive Summary," "Technical Details," "Key Insights". Explain the steps you took to arrive at your conclusions by citing selections of numerical data enclosed in quotation marks corresponding to the authors' names. [1 Paper Attached]**
**Clau-3Opus: 39.29s, 426 words**

### 34 Novel Therapeutic Targets on the Horizon An Analysis of Clinical Trials on Therapies for Bone Metastasis in Prostate Cancer, 11800 words

### Prompt 2a: Write a detailed report based on the document regarding how the authors’ methods advanced organ-specific cancer research using the following
