## Supplementary SREP for "Cancer vs. Conversational Artificial Intelligence"

### Supplementary Information SREP:

The content is available under CC BY 4.0. Refer to manuscript [Table 2] for paper references.

#### Prompts 1, 2a, 2b, 3a, 4a
#### Clau-3Opus: 3 Generations (Pages 01-06)
#### Clau-3Opus: 3 Generations (Pages 06-10)
#### Lla3.1-405: 3 Generations (Pages 10-16)
#### ChatGPT o1: 3 Generations (Pages 16-31) [details in brackets]
#### ChatGPT o1: 3 Generations (Pages 31-45) [details in brackets]

Total = 15 generations obtained December 18, 2024. Triplicates were performed sequentially without exclusions. See Supplementary SIM, SIMp, S40p, SOR, and SORp for exact prompt details.

**Prompt 1: (From "Write a detailed" - "the authors' names." on Page 1 was copied into LLM):**

#### 01 A pancreatic cancer organoid-in-matrix platform shows distinct sensitivities to T cell killing, 11803 words [See Supplementary SIM for Full Input]
#### Clau-3Opus: 47.86s, 568 words Triplicate 1

Here is a detailed report on how the authors' methods advanced the tumor immune microenvironment field based on the provided scientific paper:

Executive Summary:
In this study, Lahusen et al. developed an advanced 3D co-culture platform called InterOMaX (INTERaction with Organoid-in-MatriX) to investigate interactions between cancer cells, T cells, and stromal cells in a customizable matrix mimicking the tumor microenvironment. This versatile imaging-transcriptomic platform enabled reliable differentiation between distinct T cell response phenotypes in pancreatic ductal adenocarcinoma (PDAC) subpopulations. The model was validated using both murine PDAC cells and patient-derived PDAC organoids. Key advantages over standard co-culture models include uniformly sized organoids, maintained cellular heterogeneity,

accurate quantification of T cell infiltration, and generation of multiple complementary readouts. The platform facilitated identification and validation of CXCL17 as a candidate gene mediating T cell resistance in PDAC.

Technical Details:
The workflow involved seeding cancer cells into agarose microwell chips to rapidly generate 3D spheroids or organoids, followed by embedding in a customizable collagen matrix. T cells, either tumor-educated from mice or pre-activated human T cells, were added and co-cultured for 72h. Multiple readouts were obtained, including qPCR of T cell markers, microscopy-based quantification of T cell infiltration and cancer cell apoptosis, and transcriptomic profiling. Key technical advances include:

- Use of microwell chips for controlled organoid size and positioning
- Matrix embedding method to completely surround organoids
- Inclusion of both murine and human PDAC models
- Quantitative imaging of T cell infiltration into each organoid
- Multicellular co-cultures with stromal cells
- Identification of T cell resistant vs sensitive PDAC phenotypes

Validation experiments showed the model reflected in vivo findings, with higher T cell infiltration into sensitive vs resistant PDAC tumors in mice (Lahusen et al: "Infiltration of Cd3 positive T cells into s.c. allograft tumors from S- and R-cells. Percentage of infiltrating T cells in relation to all cells was detected by DAB staining immunohistochemistry and quantified using the Fiji DAB Macro in 10 areas/condition. N = 4.")

Key Insights:
- The InterOMaX platform enables minute cancer cell-T cell and -stromal cell interactions through a customizable matrix, providing effective control over the PDAC microenvironment. This versatile system can help deconvolute complex in vivo tumor-niche interactions.

- Distinct T cell resistant and sensitive PDAC subpopulations were reliably identified, as determined by T cell infiltration, cancer cell apoptosis, T cell activation marker expression, and transcriptomic signatures. For example, the authors state "Number of infiltrated T cells/spheroid counted with Fiji fluorescence Macro. Means were calculated for 10 spheroids/condition. $N \geq 3$", showing significant differences between resistant and sensitive PDAC spheroids.

- The chemokine CXCL17 was found to be upregulated in T cell resistant PDAC cells and its knockdown sensitized cells to T cell killing, validating its role in mediating resistance (Lahusen et al: "Knock down of CXCL17 in resistant cells caused T cell-sensitization and increased cancer cell cytotoxicity.")

- The methods were extended to patient-derived PDAC organoids, demonstrating clinical relevance. Sensitive and resistant patient organoid phenotypes were distinguished based on T cell infiltration and activation.

In summary, the InterOMaX platform developed by Lahusen and colleagues provides an advanced preclinical model to dissect immune-cancer cell interactions in the PDAC microenvironment with multiple quantitative readouts. The ability to identify and validate regulators of T cell responsiveness, such as CXCL17, demonstrates its utility for uncovering new immunotherapy targets and resistance mechanisms in PDAC. Future applications integrating additional TME components and patient-specific cells hold promise for translating insights into personalized therapeutic strategies.

### Clau-3Opus: 38.18s, 470 words Triplicate 2

Here is a detailed report on the document:

Executive Summary:
In this study, Lahusen et al. developed an advanced 3D co-culture platform called InterOMaX (INTERaction with Organoid-in-MatriX) to model interactions between cancer cells, T cells, and stromal cells in a customizable matrix that mimics the pancreatic tumor microenvironment. This system enables reliable differentiation between distinct T cell response phenotypes in pancreatic ductal adenocarcinoma (PDAC) subpopulations through multiple complementary readouts, facilitating the identification and validation of gene candidates involved in T cell resistance.

Technical Details:
Lahusen et al. built upon their previously established agarose microwell chip array system to create the InterOMaX platform. Key features include:

1. Uniformly-sized cancer spheroids/organoids embedded in a customizable collagen I matrix
2. Co-culture with tumor antigen-specific T cells and stromal cells
3. Multiple readouts:
   - T cell infiltration and cancer cell apoptosis via fluorescence microscopy and automated quantification
   - Effector T cell activation marker expression via qPCR
   - Transcriptomic analysis of cancer cells

They validated the platform using murine KPC PDAC spheroids and patient-derived organoids (PDOs). Organoid marker expression (CK19, Ki67, E-Cadherin) was maintained in the system.

The authors also incorporated human pancreatic stellate cells (hPSCs) into PDO-T cell co-cultures, showing an hPSC-mediated inhibition of T cell functionality, demonstrating the ability to model complex multicellular interactions.

Key Insights:
1. The InterOMaX platform identified distinct T cell-sensitive (S) and -resistant (R) phenotypes in both murine KPC PDAC clones and human PDOs, as evidenced by:
   - Significantly decreased T cell infiltration and cancer cell apoptosis in R vs S cells ("Number of infiltrated T cells/spheroid" and "Casp3/7 activity - Fold change (to control)" data)
   - Lower expression of T cell activation markers (IL2, IFNG, GZMB) in T cells co-cultured with R vs S cells

2. In vivo validation: Significantly increased CD3+ T cell infiltration into subcutaneous tumors from S vs R KPC cells in mice (Lahusen et al.: "Infiltration of Cd3 positive T cells into s.c. allograft tumors from S- and R-cells. Percentage of infiltrating T cells in relation to all cells was detected by DAB staining immunohistochemistry and quantified using the Fiji DAB Macro in 10 areas/condition. N = 4.")

3. Transcriptomic analysis revealed CXCL17 as a potential mediator of T cell resistance in PDAC. Knockdown of CXCL17 in R KPC cells increased T cell infiltration, activation, and cancer cell apoptosis.

4. Multicellular co-culture with hPSCs decreased expression of T cell activation markers (IL2, IFNG, EOMES) and increased regulatory T cell marker FOXP3, indicating hPSC-mediated inhibition of T cell functionality.

In summary, the InterOMaX platform advances the tumor immune microenvironment field by providing a versatile, reproducible system to dissect complex cellular crosstalk and uncover mechanisms of T cell response in PDAC, bridging the gap between simplified in vitro models and in vivo tumor complexity. The ability to incorporate patient-derived components holds promise for clinical translation in precision immunotherapy.

### **Clau-3Opus: 32.95s, 459 words Triplicate 3**

Here is a report on how the methods in the document advanced the tumor immune microenvironment field:

Executive Summary:
In this study, Lahusen et al. developed an advanced 3D co-culture platform called InterOMaX (INTERaction with Organoid-in-MatriX) to probe the interactions between pancreatic cancer cells and T cells in a customizable matrix environment. This versatile platform enabled the authors to identify distinct T cell response phenotypes in

pancreatic cancer subpopulations and uncover gene signatures associated with T cell resistance, thus advancing our understanding of the tumor immune microenvironment.

Technical Details:
The InterOMaX platform is based on a microarray setup that allows rapid generation of uniformly-sized cancer spheroids or organoids embedded in a collagen matrix. T cells are added from outside the matrix to mimic physiological conditions. The platform generates multiple complementary readouts including T cell infiltration, T cell-mediated cancer cell apoptosis, expression of T cell activation markers, and transcriptomic analysis.

The authors first validated the platform using murine KPC pancreatic cancer spheroids. "Lin et al. utilized tumor-educated (ed)T cells isolated from tumor-draining lymph nodes of immunocompetent mice carrying subcutaneous PDAC allografts" to uncover tumor antigen-specific T cell responses. Comparing T cell-sensitive (S) and -resistant (R) KPC cells, they found significantly higher "T cell infiltration rate" and "T cell-mediated cancer cell apoptosis" in S cells.

Next, the authors established the InterOMaX platform with patient-derived pancreatic cancer organoids (PDOs). Unlike classical dome cultures, "PDOs in the InterOMaX platform are located in the same optical plane and in well-defined distances from each other," enabling accurate quantification of T cell infiltration into individual organoids. To mimic the tumor microenvironment complexity, the authors incorporated human pancreatic stellate cells (hPSCs) and found "hPSC-mediated inhibition of effector T cell functionality in co-culture with PDOs."

Key Insights:
1. The InterOMaX platform advances 3D co-culture models by allowing controlled positioning of organoids in a customizable matrix, maintaining intra-organoid heterogeneity, and enabling multiple quantitative readouts of dynamic tumor-immune interactions.

2. Using this platform, Lahusen et al. identified distinct T cell response phenotypes in pancreatic cancer subpopulations. Comparative transcriptomics revealed immunomodulatory pathways enriched in T cell-sensitive cells. The authors validated CXCL17 as a candidate gene for T cell resistance - its knockdown in resistant cells increased T cell infiltration and activation.

3. Incorporating patient-derived organoids and stromal cells, the InterOMaX platform takes a key step towards mimicking the complex tumor immune microenvironment. It provides a versatile framework for mechanistic studies to unravel tumor-niche interactions and guide personalized immunotherapy decisions.

In summary, the InterOMaX platform developed by Lahusen et al. advances our toolbox for dissecting the intricate crosstalk between cancer cells, T cells, and the microenvironment. By enabling controlled, reproducible studies that capture in vivo tumor complexity, this platform brings us closer to decoding the mechanisms underlying immunotherapy resistance in pancreatic cancer.

**Prompt 2a: (From "Write a detailed" - "the authors' names." on Page 1 copied into LLM):**

### 21 A patient-specific lung cancer assembloid model with heterogeneous tumor microenvironments, 13684 words [See Supplementary SIM for Full Input] Clau-3Opus: 43.95s, 461 words Triplicate 1

Here is a detailed report on how the authors' methods advanced organ-specific cancer research based on the provided scientific paper:

Executive Summary:
In this study, Zhang et al. developed an innovative patient-specific lung cancer assembloid (LCA) model using droplet microfluidic technology. This method enabled precise manipulation of clinical microsamples and rapid generation of uniform LCAs that recapitulated the tumor microenvironment (TME) heterogeneity and replicated clinical drug responses of the original patient tumors. The LCA model represents a promising preclinical platform for advancing personalized cancer research and drug screening.

Technical Details:
The key technical advance was the development of a microinjection strategy-based droplet microfluidic technology to generate LCAs in a high-throughput manner. This involved encapsulating patient tumor-derived TME cells and lung cancer organoids (LCOs) inside gelatin methacryloyl (GelMA)-Matrigel microgels. Optimized hydrogel formulations ensured good biocompatibility and mechanical properties similar to patient lung tumors ("The hydrogel consisting of 15% (v/v) Matrigel and 6% (w/v) GelMA (termed 6–15) exposed to 90 mW of blue light at 405 nm for 40 s showed comparable mechanical properties to those of patient lung tumors (19.1 ± 0.4 vs. 27.9 ± 3.9 kPa)"). The microfluidic platform enabled generation of "~200 uniform LCAs with sizes of 400–500 μm (~0.05 μL per LCA) within 1 min" even from limited cells derived from tiny tumor needle biopsies. LCAs could be successfully cryopreserved and thawed, facilitating potential biobanking.

Key Insights:

1. LCAs maintained the heterogeneous histology, TME cellular diversity, genomic and transcriptomic landscape of parental tumors. Histological and immunofluorescence analyses showed LCAs contained "EpCAM+ tumor cells, α-SMA+ CAFs and CD45+/CD3+ immune cells" and "replicated the heterogeneous expression of CK7 and Ki67 markers within LC14 tumors, which indicated that LCAs also recapitulated the intratumoral heterogeneity".

2. Bulk RNA-seq demonstrated high transcriptomic similarity between LCAs and parental tumors (overall correlation coefficient 0.86). Whole-exome sequencing showed most somatic variants and copy number variations in parental tumors were conserved in LCAs.

3. Single-cell RNA-seq revealed LCAs maintained diverse cell types and cell-cell interactions of parental tumors. "1 week-old and 2 week-old LCAs displayed ~88% and 84% overall similarity to parental tumors, respectively."

4. The LCA model reconstructed functionally heterogeneous CAFs and enabled evaluation of their differential impact on tumor growth and drug responses compared to cancer organoids alone.

5. LCA-based drug sensitivity tests could predict patient clinical responses to chemotherapy, targeted therapy and immunotherapy with 100% accuracy in 7 out of 7 patients, highlighting the potential of the LCA model for personalized drug screening.

In summary, the patient-specific LCA model generated by innovative microfluidic technology provides a powerful preclinical platform that captures the cellular and molecular heterogeneity of lung tumors. It shows promise for advancing personalized cancer research and predictive drug testing to guide precision oncology.

### Clau-3Opus: 39.01s, 457 words Triplicate 2

Here is a detailed report on how the authors' methods advanced organ-specific cancer research based on the provided document:

Executive Summary:
Zhang et al. developed an innovative patient-specific lung cancer assembloid (LCA) model using droplet microfluidic technology. The LCAs recapitulate the heterogeneous tumor microenvironment (TME) and molecular features of parental lung tumors. Importantly, the LCA model accurately replicated clinical treatment outcomes in patients, highlighting its potential as a powerful preclinical tool for personalized drug screening.

Technical Details:

The authors generated LCAs by evenly encapsulating patient tumor-derived TME cells and lung cancer organoids (LCOs) inside GelMA-Matrigel microgels using a microinjection-based droplet microfluidic platform. Key technical advances include:

1. Rapid generation of ~200 uniform LCAs (400-500 μm in size) within 1 min from just 10 μL of hydrogel containing 10^6-10^8 cells/mL, enabling fabrication even from tiny tumor biopsies.

2. LCAs maintained good intra-batch consistency in size and cell composition, overcoming limitations of traditional co-culture and bioprinting methods.

3. The GelMA-Matrigel hydrogel provided suitable mechanical properties (compressive modulus of 19.1 ± 0.4 kPa, comparable to patient lung tumors at 27.9 ± 3.9 kPa) while maintaining high cell viability and proliferation.

4. Successful cryopreservation of LCAs, allowing biobanking for future applications.

Key Insights:
The LCA model recapitulated key features of parental lung tumors, providing several important insights:

1. LCAs maintained inter- and intra-tumor heterogeneity, exhibiting histological and cellular diversity (EpCAM+ tumor cells, α-SMA+ CAFs, CD45+/CD3+ immune cells) similar to original patient tumors.

2. Transcriptomic analysis revealed 86% overall similarity between 1-week LCAs and parental tumors, with shared enrichment of cancer-related pathways. Single-cell RNA-seq confirmed maintenance of major cell types and their interactions in LCAs.

3. LCAs could reconstruct functionally heterogeneous CAFs. For example, "LC22 CAFs with HGFhigh FGF7high expression significantly promoted the growth of cancer organoids in LCAs and improved the resistance of LCAs to chemotherapeutic drug (Taxol) treatment," while "LC23 CAFs with HGFlow FGF7low levels did not facilitate the growth of the matched cancer cells or protect tumor cells from Taxol treatment."

4. Most notably, "LCAs could accurately predict the clinical outcomes of patients with an overall accuracy of 100% (7 of 7)." For instance, "LCA46 were more sensitive to Taxol + Carboplatin + Furmonertinib than to single-agent and double-agent chemotherapy, in agreement with the clinic outcomes" for patient LC46.

In summary, Zhang et al. developed a high-throughput method to fabricate patient-specific LCAs that recapitulate the TME heterogeneity and clinical drug responses of lung tumors with high fidelity. This demonstrates the power of the LCA model as an organ-specific preclinical tool to advance personalized cancer therapy. The quantitative comparisons between LCAs and parental tumors, along with the 100% accuracy in predicting patient responses, provide strong evidence supporting the validity and utility of this approach.

### Clau-3Opus: 32.66s, 396 words Triplicate 3

Here is a detailed report on how the authors' methods advanced organ-specific cancer research based on the provided document:

Executive Summary

In this study, Zhang, Hu, Pei et al. developed an innovative patient-specific lung cancer assembloid (LCA) model using droplet microfluidic technology. The LCAs were generated by encapsulating patient tumor-derived tumor microenvironment (TME) cells and lung cancer organoids (LCOs) inside microgels. The LCAs showed good intra-batch consistency, represented the heterogeneous TME of parental tumors, and accurately replicated patients' clinical responses to therapies. This LCA model provides a valuable tool for cancer research and personalized medicine.

Technical Details

- Droplet microfluidics based on a microinjection strategy enabled precise manipulation of clinical microsamples and rapid generation of uniform LCAs. Even "10 µL hydrogels containing 108 cells mL−1 could be successfully manipulated to generate ~200 uniform LCAs with sizes of 400–500 µm (~0.05 µL per LCA) within 1 min."
- GelMA-Matrigel composite hydrogels optimized at 15% Matrigel and 6% GelMA showed "comparable mechanical properties to those of patient lung tumors (19.1 ± 0.4 vs. 27.9 ± 3.9 kPa)" while maintaining good cell viability and proliferation.
- LCAs retained the heterogeneous histology, transcriptomic profiles, and genomic alterations of parental tumors. "The overall similarity of ligand–receptor interactions of fibroblast-epithelial cells and epithelial-fibroblasts between LCAs and parental tumors was ~78% and 68%, respectively."
- Drug testing in LCAs from 7 patients "accurately predict the clinical outcomes of patients with an overall accuracy of 100%."

Key Insights

- The LCA model overcomes limitations of conventional cancer models by rapidly generating uniform assembloids that incorporate patient-specific TME components while maintaining intra-batch consistency.
- Extensive molecular and functional characterization demonstrated LCAs faithfully recapitulate the cellular diversity, cell-cell interactions, and drug response heterogeneity of parental tumors.

- The ability of LCAs to predict patient-specific responses to chemotherapy, targeted therapy, and immunotherapy combinations highlight their potential for guiding personalized treatment decisions.
- This innovative assembloid model provides a powerful preclinical platform for studying tumor biology, screening therapeutic agents, and developing precision medicine approaches in lung cancer and potentially other cancer types.

In summary, the technical advancements and rigorous biological validation provided by Zhang, Hu, Pei and colleagues represent a major step forward in developing patient-specific ex vivo cancer models that retain the complex heterogeneity of human tumors. The LCA platform holds great promise for accelerating discovery research and enabling more predictive preclinical drug testing to ultimately improve cancer treatment outcomes.

**Prompt 2b: (From "Utilize these 40" - "cancer research.' on Pages 1-58 copied into LLM):**

### Utilize these 40 summaries to draft 10 next generation cancer research studies. Cite selections of text enclosed in quotation marks corresponding to the authors' names that helped you form your conclusion. [Supplementary S40 for Full Input] Lla3.1-405: 26.08s, 626 words Triplicate 1

Based on the provided summaries, I have identified key themes and approaches that can be combined to draft 10 next-generation cancer research studies. Each proposed study leverages advances in understanding the tumor microenvironment, single-cell analysis, and precision medicine to explore novel therapeutic targets and strategies for various cancers.

1. **Personalized neoantigen-based immunotherapy for glioblastoma**
- Integrate single-cell RNA sequencing, bulk RNA sequencing, and spatial transcriptomics to identify tumor-specific neoantigens in glioblastoma patients.
- Develop personalized neoantigen-based vaccines and combine them with immune checkpoint inhibitors to enhance treatment efficacy.
- Utilize patient-derived organoids and mice models to validate treatment responses.

2. **Targeting the HIF-1α/PD-L1 pathway in colorectal cancer**
- Investigate the role of hypoxia-inducible factor-1 alpha (HIF-1α) in regulating PD-L1 expression in colorectal cancer cells.
- Use single-cell analysis and spatial transcriptomics to study the impact of the HIF-1α/PD-L1 pathway on the tumor microenvironment.

- Develop therapeutic strategies to target this pathway, including small molecule inhibitors and RNA-based therapies.

3. **Exploring the role of SPP1+ macrophages in glioma**
- Employ single-cell RNA sequencing and spatial transcriptomics to investigate the function of SPP1+ macrophages in glioma progression.
- Analyze the interaction between SPP1+ macrophages and glioma cells using co-culture systems and patient-derived organoids.
- Develop strategies to target SPP1+ macrophages for glioma treatment, including immunotherapies and small molecule inhibitors.

4. **Modulating myeloid-derived suppressor cells in breast cancer**
- Use single-cell analysis and spatial transcriptomics to study the role of myeloid-derived suppressor cells (MDSCs) in breast cancer progression.
- Investigate the relationship between MDSCs and other immune cells in the tumor microenvironment.
- Develop therapeutic strategies to modulate MDSCs, including targeting their recruitment, differentiation, and immunosuppressive functions.

5. **Precision targeting of the pancreatic cancer tumor microenvironment**
- Employ single-cell RNA sequencing, spatial transcriptomics, and bulk RNA sequencing to identify key drivers of pancreatic cancer progression.
- Develop precision medicine approaches to target the tumor microenvironment, including vascular normalization and immune checkpoint inhibition.
- Use patient-derived organoids and mice models to validate treatment responses.

6. **Investigating the role of m6A in the tumor microenvironment**
- Study the role of N6-methyladenosine (m6A) RNA modification in regulating the tumor microenvironment in various cancers.
- Use single-cell analysis and spatial transcriptomics to investigate the impact of m6A on immune cell function and tumor progression.
- Develop therapeutic strategies to target m6A, including small molecule inhibitors and RNA-based therapies.

7. **Single-cell informatics for tumor microenvironment and immunotherapy**
- Develop computational tools to integrate single-cell data from various modalities, including RNA sequencing, ATAC sequencing, and spatial transcriptomics.

- Apply machine learning algorithms to identify biomarkers and cellular features associated with treatment response and resistance.
- Use single-cell informatics to predict patient outcomes and develop personalized treatment strategies.

8. **Targeting the chemokine network in lung cancer**
- Investigate the role of chemokines in regulating the tumor microenvironment in lung cancer.
- Use single-cell analysis and spatial transcriptomics to study the impact of chemokines on immune cell recruitment and tumor progression.
- Develop therapeutic strategies to target chemokines, including small molecule inhibitors and RNA-based therapies.

9. **Modulating the coagulome in breast cancer**
- Study the role of the coagulation-related gene set (coagulome) in breast cancer progression.
- Use single-cell analysis and spatial transcriptomics to investigate the impact of the coagulome on the tumor microenvironment.
- Develop therapeutic strategies to modulate the coagulome, including targeting coagulation factors and their receptors.

10. **Exploring the role of ILF2 in breast cancer brain metastasis**
- Investigate the role of interleukin enhancer-binding factor 2 (ILF2) in breast cancer brain metastasis.
- Use single-cell analysis and spatial transcriptomics to study the impact of ILF2 on the tumor microenvironment.
- Develop therapeutic strategies to target ILF2, including small molecule inhibitors and RNA-based therapies.

These proposed studies aim to leverage cutting-edge technologies and insights into the tumor microenvironment to develop novel therapeutic strategies for various cancers. Each study has the potential to significantly advance our understanding of cancer biology and improve patient outcomes.

### Lla3.1-405: 21.44s, 509 words Triplicate 2

Based on the provided summaries, here are 10 potential next-generation cancer research studies:

between ovarian cancer cells, immune cells, and the tumor microenvironment, aiming to identify novel therapeutic targets and strategies for ovarian cancer treatment.

10. **Investigating the Role of Competing Endogenous RNA (ceRNA) Networks in Hepatocellular Carcinoma (HCC)**: Building on the work of Tang et al. (2024), this research would employ transcriptomic analysis and functional validation to investigate the roles of ceRNA networks in HCC development and progression, aiming to identify novel biomarkers and therapeutic targets for HCC treatment.

### Lla3.1-405: 13.74s, 545 words Triplicate 3

Based on the summaries provided, here are 10 potential next-generation cancer research studies that could be conducted:

1. **Identifying and targeting tumor suppressor gene-mediated immunosuppression in triple-negative breast cancer**:

This study could utilize CRISPR screens, transcriptomics, and proteomics to identify tumor suppressor genes that contribute to immunosuppression in triple-negative breast cancer. The study could also explore targeting these genes to enhance anti-tumor immunity.

2. **Developing a 3D bioprinted model of the pancreatic cancer microenvironment to study immunotherapy resistance**:

This study could create a 3D bioprinted model of the pancreatic cancer microenvironment, incorporating tumor cells, immune cells, and stromal cells. The model could be used to study the mechanisms of immunotherapy resistance and test potential therapeutic strategies.

3. **Investigating the role of regulatory T cells in the tumor microenvironment of hepatocellular carcinoma**:

This study could utilize single-cell RNA sequencing and spatial transcriptomics to characterize regulatory T cells in the tumor microenvironment of hepatocellular carcinoma. The study could also explore targeting regulatory T cells to enhance anti-tumor immunity.

4. **Developing a biomarker-based approach to predict response to immune checkpoint inhibitors in non-small cell lung cancer**:

This study could utilize single-cell RNA sequencing, bulk RNA sequencing, and machine learning algorithms to identify biomarkers that predict response to immune checkpoint inhibitors in non-small cell lung cancer.

5. **Studying the role of myeloid-derived suppressor cells in the tumor microenvironment of ovarian cancer**:

This study could utilize single-cell RNA sequencing and spatial transcriptomics to characterize myeloid-derived suppressor cells in the tumor microenvironment of ovarian cancer. The study could also explore targeting these cells to enhance anti-tumor immunity.

6. **Investigating the impact of metabolic heterogeneity on tumor progression and treatment response in clear cell renal cell carcinoma**:

This study could utilize single-cell RNA sequencing, bulk RNA sequencing, and metabolomics to investigate the impact of metabolic heterogeneity on tumor progression and treatment response in clear cell renal cell carcinoma.

7. **Developing a CAR-T cell therapy targeting glioblastoma-specific antigens**:

This study could utilize single-cell RNA sequencing and spatial transcriptomics to identify glioblastoma-specific antigens and develop a CAR-T cell therapy targeting these antigens.

8. **Investigating the role of the tumor microenvironment in promoting immune evasion in colorectal cancer**:

This study could utilize single-cell RNA sequencing, bulk RNA sequencing, and spatial transcriptomics to investigate the role of the tumor microenvironment in promoting immune evasion in colorectal cancer.

9. **Developing a biosensor-enhanced organ-on-a-chip model of the lung cancer microenvironment**:

This study could create a biosensor-enhanced organ-on-a-chip model of the lung cancer microenvironment, incorporating tumor cells, immune cells, and stromal cells. The model could be used to study the mechanisms of tumor progression and test potential therapeutic strategies.

10. **Investigating the impact of hypoxia on tumor progression and treatment response in pancreatic cancer**:

This study could utilize single-cell RNA sequencing, bulk RNA sequencing, and metabolomics to investigate the impact of hypoxia on tumor progression and treatment response in pancreatic cancer.

These studies would require a range of technical skills and expertise, including:

* Single-cell RNA sequencing and spatial transcriptomics
* CRISPR screens and gene editing
* 3D bioprinting and organ-on-a-chip technology
* Biosensor development and application
* Machine learning and computational biology
* Metabolomics and proteomics
* Cell culture and animal models
* Immunology and cancer biology

The studies would also require collaboration between researchers from different disciplines, including cancer biology, immunology, bioengineering, and computational biology.

**Prompt 3a: (From "Write a detailed" - "treatment strategies." Pages 1-31 copied into LLM):**

### Write a detailed report regarding how the authors' methods in 10 "Research" Papers further complimented the tumor immune microenvironment field represented by 10 "Review" Papers using the following format: "Executive Summary," "Technical Details," "Key Insights". Explain the steps you took to arrive at your conclusions by citing selections of numerical data enclosed in quotation marks corresponding to the authors' names. [Supplementary SIMp for Full Input]

### ChatGPT o1: 54.18s, 1635 words Triplicate 1

Below is a consolidated, integrative report linking the advancements made by the 10 "Research" papers in studying the tumor immune microenvironment (TIME) to the conceptual frameworks, challenges, and emerging opportunities highlighted in the 10 "Review" papers. This synthesis is presented in the requested format. Throughout, we reference specific numerical data and findings from the research papers by enclosing them in quotation marks along with the authors' names. We then explain the steps taken to arrive at our conclusions.

---

**Executive Summary**

The 10 “Review” papers collectively emphasize the complexity of the tumor immune microenvironment, highlighting critical areas such as metabolic reprogramming, cellular heterogeneity, stromal-immune cell interactions, hypoxia-driven signaling, spatially resolved multi-omics analysis, and the development of novel immunotherapeutic strategies (including CAR-T cell therapies and exosome-based interventions). They note that future progress relies on physiologically relevant models, robust multi-omics platforms, single-cell resolution techniques, and approaches that integrate the metabolic, spatial, and temporal dimensions of tumor-immune dynamics.

The 10 “Research” papers complement these visions by developing and applying innovative experimental platforms and analytical methodologies that address many challenges posed by the TIME. For instance, organoid-in-matrix platforms (Lahusen et al.), multiplexed imaging (Kim et al.), single-cell RNA-seq analysis (Tang et al.), multi-omics pipelines (Ferri-B., S., et al.), and targeted interventions such as IRE (Imran, K., et al.) collectively advance the field from both a conceptual and technical standpoint. These studies not only confirm insights from the reviews—such as the importance of the extracellular matrix (ECM), immune cell infiltrate, and metabolic cues—but also identify new targets and biomarkers (e.g., INHBA(+) CAFs from Hu, Y., et al. and SPP1+ macrophages from Tang, W., et al.). In doing so, they illuminate potential therapeutic strategies that could have broad implications for overcoming immunotherapy resistance as discussed in the reviews.

---

**Technical Details**

**1. Physiologically Relevant 3D Models and Spatial Analysis**
- The reviews (e.g., Zhang, Z., et al. 2024; Zhou, Z., et al. 2024) underscore the need for advanced 3D models. Lahusen et al. (2024) developed the InterOMaX platform embedding pancreatic organoids in collagen I matrix to study T cell infiltration and killing dynamics. They showed that "T cell infiltration and tumor cell killing varied significantly across different PDAC clones" (Lahusen et al.)—a direct experimental demonstration of heterogeneity discussed in 3D modeling review papers.

- Reviews like Tian, J., et al. (2024) highlight spatial transcriptomics and multiplex imaging. Kim et al. (2024) applied multiplexed immunohistochemistry and clustering algorithms to visualize Aryl hydrocarbon receptor patterns in solid tumors. By clustering "13 features representing AhR expression across 513 patient samples" (Kim et al.), they achieved the spatial and phenotypic resolution championed by the reviews.

**2. Targeting the Tumor Stroma and Immunosuppressive Cells**
- Review articles (e.g., Kundu, M., et al. 2024; Lu, J., et al. 2024) emphasize CAFs, MDSCs, and Tregs as key therapeutic targets. Hu, Y., et al. (2024) identified INHBA(+) CAFs as pro-tumorigenic and immunosuppressive, correlating with higher Treg infiltration. Quantitative data showed a "significantly higher ratio of INHBA(+) fibroblasts in metastatic vs. primary tumors" (Hu, Y., et al.). This finding validates review assertions about fibroblast heterogeneity and immunosuppression and informs strategies to target CAF subsets, as suggested in reviews on overcoming immunotherapy resistance.

- Similarly, Zahraeifard, S., et al. (2024) used in vivo CRISPR screens and multi-omics to show that loss of certain tumor suppressors (e.g., NF1, TSC1) induces an inflammatory environment favoring LAG3+ T cell-mediated suppression. They found that "70-80% of CD8+ T effector cells were LAG3+ in TSG-deficient tumors" (Zahraeifard, S., et al.). The reviews by Li, Y., et al. (2024) and Du, G., et al. (2024) which discuss Treg and checkpoint-mediated suppression, are complemented by this mechanistic demonstration.

**3. Integrating Metabolism, Hypoxia, and Immune Regulation**
- Many reviews (e.g., Zhang, H., et al. 2024; Han, X., et al. 2024) focus on metabolic reprogramming and epitranscriptomic changes. Dai, L., et al. (2024) provide single-cell and spatial transcriptomics data linking glycolysis in diffuse large B-cell lymphoma (DLBCL) to poor prognosis and immune exhaustion. They showed that "seven glycolysis genes (STMN1, ENO1, LDHA, TPI1, CDK1, PKM, PPIA) correlated with worse outcomes" (Dai, L., et al.), mirroring the metabolic-immune interplay described in the reviews.

- Sun, J., et al. (2024) connect hypoxia-driven HIF-1α signaling to PD-L1 upregulation and immune escape in colorectal cancer. They demonstrated that "HIF-1α was positively correlated with PD-L1 ($r = 0.612$) in CRC" (Sun, J., et al.), reinforcing the concept from Han, X., et al. (2024) and Zhang, H., et al. (2024) that hypoxic and metabolic alterations drive immunosuppression.

**4. Advanced Multi-Omics and Single-Cell Characterization**
- Review papers (e.g., Zhou, Z., et al. 2024; Tian, J., et al. 2024) stress the value of multi-omics, including proteomics, metabolomics, and spatial transcriptomics. Ferri-B., S., et al. (2024) integrated non-targeted MSI, Stereo-seq transcriptomics, and seqIF to map "948 metabolites, 164 glycans, and 586 peptides" across 3D tissue sections. This multi-omics synergy exemplifies what the reviews call for—holistic, spatially resolved profiling to decode the TIME's complexity.

- Tang, W., et al. (2024) used single-cell RNA-seq to identify TAM-SPP1 populations in gliomas associated with "significantly worse overall survival ($p<0.0001$)" (Tang, W., et al.). This matches the single-cell informatics strategies outlined by Tian, J., et al. (2024) and shows how such methods identify novel prognostic markers and therapeutic targets.

**5. Interventions and Combined Modalities**
- The reviews (e.g., Zhou, Z., et al. 2024; Kundu, M., et al. 2024) highlight the importance of rational combination therapies. Imran, K., et al. (2024) demonstrated that irreversible electroporation (IRE) creates a pro-inflammatory environment but leads to IFN-γ-driven PD-L1 upregulation, suggesting that combination with checkpoint inhibitors might improve outcomes. They measured "IFN-γ increase peaking at day 9 and PD-L1 peaking at day 14 post-IRE" (Imran, K., et al.), supporting the review-driven concept that synergy between local ablation and immunotherapy can boost therapeutic effects.

- Zheng, K., et al. (2024) defined TIDE-based molecular subtypes that correlate with immunotherapy responses across cancers. Their approach aligns well with the reviews' calls for predictive biomarkers and molecular classification strategies. Similarly, introducing targeted therapies (e.g., anti-LAG3 in Zahraeifard, S., et al.) corroborates the reviews' emphasis on precision immunotherapy.

---

**Key Insights**

1. **Bridging Conceptual Gaps with Experimental Models**:
The research papers offer tangible platforms—3D organoid-in-matrix (Lahusen et al.), advanced immunohistochemistry arrays (Kim et al.), and single-cell multi-omics (Ferri-B., S., et al.; Tang, W., et al.)—that embody the principles outlined in the reviews. These methods recapitulate the TME's complexity and enable real-time interrogation of cell-cell interactions, metabolism, and spatial architecture highlighted by Zhang, Z., et al. (3D models), Zhou, Z., et al. (multi-omics), and Tian, J., et al. (single-cell informatics).

2. **Context-Dependent Immune Interactions**:
Research findings show that subsets of CAFs (Hu, Y., et al.) or tumor-associated macrophages (Tang, W., et al.; Dai, L., et al.) can drive immune suppression, aligning with the nuanced views of TME heterogeneity in reviews (Safaei, S., et al.; Li, Y., et al.). Identifying specific subpopulations (e.g., INHBA(+) CAFs, SPP1+ macrophages) transforms theoretical complexity into actionable targets.

3. **Metabolic and Hypoxic Targeting Strategies**:
By correlating gene expression signatures and metabolite distributions (Dai, L., et al.; Sun, J., et al.), the research complements review discussions about metabolic and hypoxic niches (Zhang, H., et al.; Han, X., et al.). This points

towards treatments that normalize the metabolic environment or disrupt HIF-1α/PD-L1 signaling to enhance immunotherapy.

4. **Moving Towards Precision Immunotherapy**:
The integration of CRISPR screens (Zahraeifard, S., et al.), IRE-based immunomodulation (Imran, K., et al.), TIDE subtyping (Zheng, K., et al.), and in vitro co-culture studies (Lahusen et al.) reflects the reviews' visions of more personalized immunotherapies. As suggested by Lu, J., et al. (on MDSCs) and Du, G., et al. (on Tregs in HCC), these technologies and integrative approaches lay the groundwork for overcoming resistance and tailoring treatments to individual patient tumor microenvironments.

---

**Explanation of Steps Taken to Arrive at Conclusions**

1. **Cataloging Key Themes from Reviews**:
First, we examined the 10 "Review" papers to identify recurring themes: the complexity of the TME, the need for spatial and multi-omics analyses, the importance of 3D models, the role of immunosuppressive cell types, metabolic and hypoxic influences, and strategies for improving immunotherapy.

2. **Mapping Research Papers to Review Themes**:
Next, we analyzed each of the 10 "Research" papers to identify which aspects of the TIME they addressed and which methodologies they introduced. We took note of techniques such as single-cell RNA-seq, multiplex imaging, IRE, organoid modeling, and CRISPR screens, as well as their quantifiable findings (e.g., "CXCL17" upregulation by Lahusen et al., "70-80%" LAG3+ T cells in Zahraeifard, S., et al., "513 patient samples" analyzed by Kim et al.).

3. **Integrating Data Points**:
We then correlated the detailed technical approaches of the research papers with the challenges and solutions outlined by the reviews. For instance, the review by Zhang, Z., et al. stressed 3D bioprinted models to overcome immunotherapy resistance, and Lahusen et al. provided a complementary organoid-in-matrix system. We quoted specific numerical data (e.g., "513 patient samples" from Kim et al., the percentage of LAG3+ T cells from Zahraeifard, S., et al.) to substantiate these connections.

4. **Synthesizing Into Cohesive Insights**:
Finally, we wove together the research findings with the conceptual frameworks of the reviews, showing how each research paper's methodological innovations and data-driven insights fill knowledge gaps, validate theoretical predictions, or open new translational avenues suggested by the reviews.

By following these steps and integrating empirical findings with conceptual frameworks, we demonstrated how the authors' methods in the 10 "Research" papers advance and complement the tumor immune microenvironment field as portrayed by the 10 "Review" papers.

---

In conclusion, the synergy between the empirical methodologies of the "Research" papers and the conceptual, integrative overviews offered by the "Review" papers yields a richer understanding of the TIME. This combined perspective not only validates and refines existing theories but also guides the future development of personalized, metabolism-aware, and spatially resolved immunotherapies.

**ChatGPT o1 Details: 50 Words**

[Piecing together

I'm integrating the methods from 10 research papers with 10 review papers, examining their contributions to the tumor immune microenvironment.

Showcasing alignment

I'm piecing together 10 research and review papers, showing how research advances TME insights and reveals novel therapies, all while achieving synergy with the review papers.]

### ChatGPT o1: 51.45s, 1570 words Triplicate 2

Below is a detailed integrated report showing how the 10 original "Research" Papers' methodologies build upon and complement concepts and directions highlighted by the 10 "Review" Papers in the tumor immune microenvironment (TIME) field. The format includes: "Executive Summary," "Technical Details," and "Key Insights." At the end, the steps taken to arrive at these conclusions are explained, with selected numerical data enclosed in "quotation marks" and associated with the authors' names as requested.

**Executive Summary**

The 10 “Research” Papers each employed advanced experimental, computational, and analytical techniques to dissect the tumor immune microenvironment and identify strategies to overcome immunosuppression and therapy resistance. These investigations align closely with the themes presented in the 10 “Review” Papers, which emphasize multidimensional omics characterization, metabolic reprogramming, spatial modeling, and the use of advanced 3D tissue models to recapitulate the complexity of tumor-immune interactions. By integrating cutting-edge platforms—such as organoid-in-matrix co-cultures, single-cell and spatial transcriptomics, multiplexed immunohistochemistry, CRISPR screens, and metabolite profiling—the research papers illustrate how theoretical frameworks and mechanistic hypotheses proposed in the reviews can be operationalized to yield actionable insights.

For instance, research on pancreatic cancer organoids and T cell infiltration (Lahusen et al.) and AhR expression patterns in immune cells across multiple solid cancers (Kim et al.) complements review articles emphasizing the importance of 3D models (Zhang, Z., et al.) and multi-omics integration for deciphering TIME complexities (Zhou, Z., et al.). Similarly, identifying immunosuppressive fibroblast subsets (Hu, Y., et al.), mapping tumor-associated macrophage (TAM) phenotypes (Tang, W., et al.), and examining glycolytic malignant B cell clusters (Dai, L., et al.) directly build upon review narratives that highlight how metabolic reprogramming (Zhang, H., et al.), TAM heterogeneity (Safaei, S., et al.; Lu, J., et al.), and regulatory cell subsets (Li, Y., et al.; Du, G., et al.) shape immune evasion and therapy resistance. The methods employed in the research studies are therefore both informed by and serve to advance the fields described in the review literature—particularly with respect to identifying new immunotherapeutic targets, refining prognostic biomarkers, and validating strategies for immunotherapy personalization.

**Technical Details**

In the research articles, investigators leveraged advanced techniques that the reviews propose as essential for next-generation immuno-oncology research:

1. **3D and Organoid-Based Platforms**:
   Lahusen et al. used a “3D co-culture InterOMaX platform” embedding PDAC organoids in a collagen matrix to visualize “T cell infiltration” and test gene candidates like “CXCL17” for T cell resistance. Their methods align with review insights from Zhang, Z., et al. who stress how “3D bioprinted tumor models” help recapitulate real TME conditions and overcome immunotherapy resistance.

2. **High-Throughput Spatial Proteomics and Transcriptomics**:
   Kim et al. deployed multiplexed immunohistochemistry on “513 patient samples” to cluster aryl hydrocarbon receptor (AhR) expression patterns in the TIME. Similarly, Ferri-B., S., et al. integrated non-targeted mass

spectrometry imaging and Stereo-seq transcriptomics. These approaches complement Zhou, Z., et al.'s multi-omics perspective and Tian, J., et al.'s single-cell informatics pipeline, which highlight integrating spatial, single-cell, and metabolic data as crucial next steps.

3. **Uncovering Immunosuppressive CAFs and Macrophages**:
Hu, Y., et al. identified "INHBA(+) CAFs" and demonstrated their link to Treg infiltration and shorter survival. Tang, W., et al. used single-cell RNA-seq to reveal "SPP1+ macrophages" correlating with poor glioma prognosis. These data-driven discoveries follow the logic described by Safaei, S., et al. and Kundu, M., et al., who review how stromal and macrophage subsets modulate the TIME. They also resonate with Lu, J., et al.'s discussion of MDSCs and Zhang, H., et al.'s coverage of metabolic reprogramming shaping MDSC and macrophage function.

4. **Metabolic Profiling for Therapeutic Targeting**:
Dai, L., et al. identified highly glycolytic malignant B cells and "IFN_TAMs" driving immune exhaustion in DLBCL. Metabolic studies confirm predictions from reviews by Zhang, H., et al. and Han, X., et al., who posit that metabolic reprogramming (e.g., lipid and glycolysis pathways) influences T cell dysfunction and resistance to immunotherapy.

5. **Genetic and Epigenetic Screens**:
Zahraeifard, S., et al. performed in vivo CRISPR screens to identify tumor suppressors (NF1, TSC1, TGFβRII) whose loss fosters a LAG3+ T cell-driven immunosuppressive environment. Such work takes cues from review discussions (Du, G., et al.; Li, Y., et al.) on how regulatory T cells and checkpoint pathways are shaped by underlying genetic alterations.

6. **Hypoxia and Immune Checkpoints**:
Sun, J., et al. showed that HIF-1α/PD-L1 axis promotes immune escape in CRC, a mechanistic link repeatedly emphasized in reviews like Han, X., et al. and Kundu, M., et al. regarding how environmental stressors, including hypoxia, influence epitranscriptomic modifications and checkpoint signaling.

7. **Subtyping and Prognostic Modeling**:
Zheng, K., et al. created TIDE-based molecular subtyping to predict immunotherapy response. This corresponds to the reviewers' calls (Zhou, Z., et al.; Tian, J., et al.) for integrative computational methods and prognostic modeling using single-cell and multi-omics pipelines to inform clinical decision-making.

**Key Insights**

1. **Integration of Platforms and Data Modalities**:
The research studies directly implement the multi-omics and single-cell technologies that review articles designate as critical for next-generation immunotherapy. For example, Ferri-B., S., et al. combined "948 metabolites, 164 glycans, and 586 peptides" with spatial transcriptomics, as predicted by Zhou, Z., et al., Han, X., et al., and Tian, J., et al.

2. **Validation of Hypothesized Immune Evasion Mechanisms**:
By experimentally manipulating genes like "CXCL17" (Lahusen et al.), analyzing large patient cohorts of "513 samples" for AhR expression (Kim et al.), and pinpointing metabolic bottlenecks in DLBCL (Dai, L., et al.), the research builds directly on the mechanistic frameworks from reviews addressing metabolic interplay (Zhang, H., et al.), exosome-mediated immunomodulation (Safaei, S., et al.), and Treg/MDSC control (Li, Y., et al.; Lu, J., et al.).

3. **Actionable Targets for Immunotherapy Enhancement**:
Research articles identify immune checkpoints (LAG3, PD-L1), CAF-specific factors (INHBA), and signaling molecules (HIF-1α) as tangible targets. This echoes the review findings that highlight checkpoint pathways, epigenetic modifiers, and metabolic regulators as promising intervention points. The consistency between proposed theoretical targets in reviews and experimentally validated targets in research papers underscores a cohesive evolution from theoretical knowledge to practical application.

4. **Refined Tumor Classification and Personalized Treatment**:
The novel subtyping (Zheng, K., et al.) and improved prognostic models resonate with review articles (Zhou, Z., et al.; Tian, J., et al.) that stress the importance of computational methods in stratifying patients for precision immunotherapies. By confirming that certain metabolic or immunosuppressive signatures (e.g., "TAM-SPP1" in Tang, W., et al. or the "IFN_TAMs" in Dai, L., et al.) correlate with prognosis, these research papers turn theoretical frameworks into clinical tools.

**Explanation of Steps Taken and Use of Numerical Data**

To arrive at these conclusions, the following steps were taken:

1. **Comparing Methodologies and Aims**:
Each of the 10 research papers was examined for its methodological approaches (e.g., organoid culture systems, single-cell sequencing, in vivo CRISPR screens) and linked to themes from the 10 review papers. For instance, Lahusen et al. used a 3D organoid system and "CXCL17" knockdown experiments—techniques frequently advocated in review articles (Zhang, Z., et al.) to overcome 2D culture limitations.

2. **Identifying Parallel Concepts in Reviews**:

The reviews propose integrating multi-omics, single-cell analytics, and spatial methods. Research papers like Kim et al. analyzed "513 patient samples" using multiplexed IHC and image cytometry, exactly what Zhou, Z., et al. encourage (large-scale, multi-omics profiling). Specific numeric data from Kim et al. ("513 patient samples") highlight the scale and rigor aligning with big-data approaches reviewed by Tian, J., et al.

3. **Noting Matching Findings and Themes**:

Multiple research papers identified specific immunosuppressive subsets (e.g., Hu, Y., et al. found "INHBA(+) CAFs" correlated with "significantly higher Treg ratios"), which ties directly into the review discussions on stromal cell-mediated immune escape (Kundu, M., et al.; Safaei, S., et al.). The numeric correlation "significantly higher" ratio supports the quantifiable impact Tregs have in shaping the TME as theorized in Li, Y., et al.

4. **Extracting Numerical Evidence**:

When linking methods to outcomes, numeric data offer concrete evidence. For example, Dai, L., et al. identified "7 glycolysis genes" that correlate with poor prognosis, reflecting the reviews' emphasis on metabolic targets. Zahraeifard, S., et al. performed CRISPR screening of "102 TSGs" and found that knocking down NF1, TSC1, and TGFβRII leads to altered immune infiltration, paralleling the review-driven concept that genetic alterations in tumors shape immune landscapes (Du, G., et al.).

Another numeric instance: Tang, W., et al. show that elevated TAM-SPP1 correlates with "poor survival ($p < 0.0001$)", giving quantitative weight to the review claims that specific macrophage subsets are prognostic biomarkers.

Lastly, Ferri-B., S., et al. integrating "948 metabolites," "164 glycans," and "586 peptides" is a numeric illustration of how broad omics coverage, as advocated by Zhang, H., et al. and Han, X., et al., reveals novel metabolic and immunosuppressive landscapes.

5. **Consolidating Insights**:

By systematically cross-referencing research methodologies and numeric results with review predictions, we confirm that the new experimental tools and datasets (3D models, single-cell/spatial technologies, metabolomics, CRISPR screens) provide the mechanistic and phenotypic clarity that the reviews identify as urgent needs.

In summary, the final integrated view demonstrates that the research papers do not stand alone; rather, they apply and validate the multi-omics, spatial, and mechanistic frameworks proposed in the reviews. The numeric data (e.g., "513 patient samples," "102 TSGs," "7 glycolysis genes," "948 metabolites") from the research studies confirm the scale, complexity, and impact of methods the reviews recommend. Together, these contributions enrich the field's understanding of how best to tackle immune evasion and improve cancer immunotherapy.

**ChatGPT o1 Details: 65 Words**

[Drawing conclusions

I'm pulling together "Research" and "Review" papers to highlight how the methods of the research papers enhance the field. This involves summarizing key insights and technical details to create a comprehensive integrating document.

Synthesizing data

I'm considering how to integrate quantitative data from the research and review papers to clearly illustrate the progression from methods to insights, ensuring a comprehensive and cohesive finale.]

### ChatGPT o1: 41.98s, 1704 words Triplicate 3

Below is an integrated, detailed report that connects how the methods employed in the 10 selected **Research Papers** build upon and complement the theoretical and conceptual frameworks presented by the 10 selected **Review Papers** in the field of tumor immune microenvironment (TIME). The report is structured into "Executive Summary," "Technical Details," and "Key Insights," as requested. Following the report, an explanation of the steps taken to arrive at these conclusions is provided, including selected numerical data in quotation marks that correspond to the authors' names.

--------------------------------------

**Executive Summary**

The 10 **Review Papers** collectively underscore critical themes in the evolving field of the tumor immune microenvironment (TIME), emphasizing the importance of 3D models, single-cell multi-omics technologies, metabolic reprogramming, immunosuppressive cell populations, immune checkpoints, and the role of RNA modifications (e.g., m6A) in shaping immune evasion. They provide theoretical frameworks suggesting that comprehensive and spatially resolved profiling, along with advanced bioengineering and immunomodulatory strategies, are crucial to understanding and overcoming resistance to immunotherapies.

The 10 **Research Papers** translate these conceptual advancements into actionable methodologies. Through platforms such as organoid-in-matrix (InterOMaX) co-cultures, single-cell and spatial profiling (scRNA-seq, multiplex imaging, spatial transcriptomics), selective genetic screens (CRISPR-based approaches), and integrated multi-omics pipelines, these studies deliver new insights into how specific cell populations, metabolic pathways, and molecular signals drive immune evasion. For instance, authors have identified distinct T cell-resistant cancer cell subpopulations, validated tumor-associated fibroblast subsets that promote immunosuppression, uncovered metabolic signatures linked to dysfunctional antitumor immunity, and demonstrated that interventions like irreversible electroporation (IRE) or knockdown of immunosuppressive genes can remodel the TIME.

Together, these research methods closely align with and expand on the conceptual scaffolding presented in the review literature. By bridging theory with cutting-edge experimental platforms, the research studies confirm the importance of integrating 3D modeling, multi-omics, and spatial analysis approaches to dissect tumor-immune interactions and guide new immunotherapeutic strategies.

--------------------------------------

**Technical Details**

1. **3D Model Systems & Bioprinting:**
   - The **Review Papers** (e.g., Zhang, Z., et al. 2024) highlight 3D bioprinted tumor models as next-generation platforms for simulating the authentic tumor microenvironment.
   - The **Research Paper** by Lahusen, A., et al. 2024 uses the InterOMaX platform with organoids in collagen to measure T cell infiltration and apoptosis. Such a system mirrors the recommendations of bioprinting reviews by enabling "uniformly-sized 3D cultures" and "customizable matrix" composition, directly validating the importance of spatial complexity discussed in the review literature.

2. **Advanced Spatial and Single-Cell Profiling:**
   - Multiple reviews (e.g., Zhou, Z., et al. 2024; Tian, J., et al. 2024) emphasize single-cell and spatial multi-omics to decode cellular heterogeneity and cell-cell communication networks.
   - In the research realm, Kim, D., et al. 2024 employ multiplexed immunohistochemistry and image cytometry to classify "513 patient samples" into distinct AhR expression clusters. Similarly, Ferri-B., S., et al. 2024 integrate non-targeted mass spectrometry imaging (MSI), Stereo-seq transcriptomics, and seqIF to achieve "3D targeted and non-targeted multiplex multi-omics analyses," directly operationalizing the multi-omics approach praised in reviews.

3. **Immunosuppressive Subsets and Immune Evasion Mechanisms:**

- Reviews like Lu, J., et al. 2024 and Li, Y., et al. 2024 focus on immunosuppressive cell types (MDSCs, Tregs) and their intricate roles. Kundu, M., et al. 2024 further detail how CAFs and TAMs shape the immunosuppressive niche.
- In line with these concepts, Hu, Y., et al. 2024 identify "INHBA(+)" CAFs that create an immunosuppressive environment in ovarian cancer, while Tang, W., et al. 2024 pinpoint SPP1+ TAMs as key drivers of poor prognosis in glioma. Both findings confirm review-based predictions that fibroblasts and macrophages have specialized subtypes linked to immune escape.

4. **Metabolic Reprogramming:**
   - Review articles like Zhang, H., et al. 2024 discuss how altered tumor and immune cell metabolism contributes to immune evasion, suggesting that metabolic inhibitors could improve immunotherapy.
   - Research by Dai, L., et al. 2024 identifies glycolysis-driven B cells and IFN_TAMs in diffuse large B-cell lymphoma (DLBCL) that correlate with an exhausted immune microenvironment. Such data provide empirical support for the review-based premise that metabolic states can define immune infiltration patterns and therapy responses.

5. **Hypoxia and RNA Modifications:**
   - Several reviews (Han, X., et al. 2024) highlight the role of m6A RNA modifications and hypoxia in shaping immunosuppression and drug resistance.
   - Research by Sun, J., et al. 2024 shows that the HIF-1α/PD-L1 pathway is activated under hypoxic/inflammatory conditions to promote immune escape in colorectal cancer. This aligns with the review perspective that RNA modifications and hypoxic signaling are key molecular handles influencing TIME and therapeutic outcomes.

6. **Therapeutic Interventions and Resistance Mechanisms:**
   - Reviews like Du, G., et al. 2024 and Zhou, Z., et al. 2024 emphasize that understanding the TIME at cellular and molecular levels can suggest improvements to CAR-T cell therapy and checkpoint blockade strategies.
   - Research by Zahraeifard, S., et al. 2024, through CRISPR-based TSG inactivation, reveals that LAG3+ T cell-mediated immunosuppression can be overcome by combined immunotherapies. Similarly, Imran, K., et al. 2024 demonstrate that irreversible electroporation can induce a pro-inflammatory environment, albeit transiently, suggesting combination therapies to maintain antitumor immune responses.

7. **Predictive Biomarkers and Personalized Approaches:**
   - The TIDE subtyping strategies discussed in Zheng, K., et al. 2024's research expand on the review themes of patient stratification and biomarker development (seen in Tian, J., et al. 2024). This refined classification approach helps identify which patients respond to immunotherapies, echoing the review-based call for integrative and predictive modeling of TIME features.

-------------------------------------
**Key Insights**

1. **Multi-Dimensional Modeling is Critical:**
The combined insights show that 3D models, single-cell, and spatial multi-omics platforms are not just theoretical recommendations (from reviews) but practical tools (from research) that yield robust, physiologically relevant data on immune cell infiltration, cellular states, and microenvironmental cues.

2. **Targeting Specific Cell Subsets Can Overcome Resistance:**
Both reviews and research highlight that identifying and targeting unique immune-suppressive populations—such as INHBA(+) CAFs, SPP1+ TAMs, and LAG3+ T cells—can break immunotherapy resistance. Studies identify "CXCL17" (Lahusen, A., et al. 2024) and "Activin A/INHBA" (Hu, Y., et al. 2024) as actionable targets, confirming the review suggestion that intervention at the level of critical TME regulators is a key strategy.

3. **Metabolic and Hypoxic Niches Influence Immune Function:**
Research findings on glycolytic B cells (Dai, L., et al. 2024) and HIF-1α/PD-L1 induction (Sun, J., et al. 2024) map directly onto review-based predictions that nutrient competition, lactate accumulation, and hypoxia modify immune cell function. Intervening in these metabolic pathways may restore T cell functionality and sensitize tumors to immunotherapy.

4. **Integration of Technologies Guides Clinical Translation:**
The synergy between the technical methods (like multiplex immunohistochemistry, CRISPR screens, and multi-omics) and conceptual frameworks (highlighted in the reviews) guides the rational design of combination therapies (PD-L1 blockade with metabolic modulators, CAF reprogramming, or MDSC inhibition). The TIDE-based subtyping (Zheng, K., et al. 2024) exemplifies how integrating multi-layered data can produce clinically actionable predictive models.

5. **Personalized Immunotherapy is the Future:**
Reviews predict a paradigm shift toward personalized immunotherapies based on thorough TME characterization. The research studies validate this shift: patient-derived organoids (Lahusen, A., et al. 2024) and predictive biomarkers (Zheng, K., et al. 2024) point toward tailored strategies that consider unique TME features and genetic backgrounds.

-------------------------------------
**Explanation of Steps and Use of Numerical Data**

To derive these conclusions, the following steps were taken:

1. **Synthesis of Themes from Review Articles:**
The review papers outlined key conceptual frameworks: importance of 3D modeling (Zhang, Z., et al.), single-cell and multi-omics profiling (Zhou, Z., et al.; Tian, J., et al.), metabolic and immunosuppressive states (Zhang, H., et al.; Lu, J., et al.), the role of RNA modifications (Han, X., et al.), Treg and CAF/TAM interplay (Kundu, M., et al.; Li, Y., et al.), and predictive biomarkers (Du, G., et al.).

2. **Matching Research Methodologies to Review Concepts:**
Each research paper was examined to identify how their methodologies support or extend the review-based concepts:
- Lahusen, A., et al. leveraged organoid-in-matrix platforms, validating the 3D model emphasis. They report "distinct T cell response phenotypes" and the identification of "CXCL17" as a T cell resistance mediator.
- Kim, D., et al. profiled "513 patient samples" using multiplex IHC to classify AhR patterns, aligning with single-cell and spatial profiling approaches suggested in reviews.
- Hu, Y., et al. showed how INHBA(+) CAFs shape an immunosuppressive environment, confirming fibroblast heterogeneity and its functional importance as discussed in Kundu, M., et al.
- Imran, K., et al. used IRE and documented temporal changes in MDSCs and T cells, highlighting transient immunostimulatory windows. They measured "100 pulses at 800 V" and observed "peaking IFN-γ at day 9," providing quantitative support for dynamic TIME remodeling.
- Zahraeifard, S., et al. combined CRISPR screening with multi-omics to find LAG3+ T cell-mediated immune suppression linked to tumor suppressor gene loss. "TS-Imm signature" was associated with "decreased OS and RFS ($p = 0.008$)"—numeric data confirm the clinical relevance.
- Ferri-B., S., et al. integrated MSI, Stereo-seq, and seqIF, using "17 serial 5 μm FFPE tissue sections" and measuring "948 metabolites" to map the TME in 3D, directly implementing recommendations for multi-omics and spatial resolution.
- Tang, W., et al. performed single-cell analysis in gliomas to identify SPP1+ macrophages. "SPP1+ TAMs were significantly correlated with poor OS ($p < 0.0001$)"—this quantitative result validates prognostic utility suggested by reviews.
- Dai, L., et al. uncovered glycolysis-driven B cells associated with immune exhaustion in DLBCL. They identified "7 glycolysis genes" with prognostic significance, echoing metabolic reprogramming discussions in Zhang, H., et al.
- Sun, J., et al. showed HIF-1α/PD-L1 correlation in CRC: "$r = 0.612$" and the effect of hypoxia on immune escape, confirming hypotheses about hypoxia's immunosuppressive role.

- Zheng, K., et al. validated TIDE-based subtyping in bladder cancer, identifying "three TIDE subtypes" and linking them to immunotherapy response patterns. "Hypergeometric test" results demonstrated meaningful subtype distinctions.

3. **Integrating Data to Form a Unified Narrative:**
By comparing these detailed approaches and results to the review-based concepts, a coherent narrative emerges. Numerical data, such as "513 patient samples" (Kim, D., et al.), "day 9 IFN-γ peak" (Imran, K., et al.), "p = 0.0001" correlation (Tang, W., et al.), and "decreased OS and RFS (p = 0.008)" (Zahraeifard, S., et al.), were used as evidence of robust, statistically significant findings reinforcing the conceptual frameworks from the reviews.

This careful matching of experimental evidence with theoretical insights from the reviews illustrates how the research methods validate, refine, and advance the field's understanding of the TIME.

**ChatGPT o1 Details: 31 Words**

[Synthesizing insights

I'm gathering insights from 10 research and 10 review papers on the tumor immune microenvironment. This involves highlighting key findings, methodologies, and data to create a comprehensive integrated report.]

**Prompt 4a: (From "Write a detailed" - "cancer research." on Page 1-29 copied into LLM):**

### Write a detailed report regarding how the authors' methods in 10 "Emerging" Papers further complimented organ-specific cancer research represented by 10 "Historical" Papers using the following format: "Executive Summary," "Technical Details," "Key Insights". Explain the steps you took to arrive at your conclusions by citing selections of numerical data enclosed in quotation marks corresponding to the authors' names. [Supplementary SORp for Full Input]

### ChatGPT o1: 28.13s, 1528 words Triplicate 1

Below is a comprehensive report integrating insights from 10 "Emerging" and 10 "Historical" papers. The "Emerging" papers introduce cutting-edge models, technologies, and analytical approaches, while the "Historical" papers provide foundational context and earlier methodologies. Together, they illustrate how recent methodological breakthroughs complement and extend established organ-specific cancer research.

**Executive Summary**
Organ-specific cancer research has traditionally relied on model systems, imaging, and bulk sequencing methods to understand tumor biology. The 10 “Historical” studies provided foundational insights into tumor microenvironments (TMEs), immunological dynamics, and the efficacy of various therapies, but often faced limitations in recapitulating human tumor complexity, spatial context, and intratumoral heterogeneity.

In contrast, the 10 “Emerging” studies leverage next-generation models—such as patient-specific assembloids, cancer-on-chip platforms, single-cell multi-omics, and spatial transcriptomics—to more accurately model patient tumors and dissect the TME at unprecedented resolution. These newer methods build directly on the needs and knowledge gaps identified by the earlier studies, enabling researchers to:
1. Better mimic native tumor conditions with 3D organ-on-chip and assembloid models.
2. Achieve single-cell and spatial resolution to identify distinct cell populations and metabolic states.
3. Validate and improve on prognostic biomarkers and therapeutic targets identified through bulk or lower-resolution methods.
4. Tailor immunotherapies and targeted treatments based on patient-specific tumor features and dynamic responses.

By bridging earlier foundational work (the “Historical” papers) with the advanced tools and models described in the “Emerging” papers, researchers can move toward more precise, personalized, and effective treatments for a range of organ-specific cancers.

**Technical Details**
The “Historical” research often centered on understanding tumor biology through traditional tissue cultures, immunohistochemistry, bulk RNA sequencing, and targeted biomarker analyses. For example:

- **Glioblastoma Models & Biosensors (Thenuwara, G., et al., 2024)**: The historical perspective described integrating biosensors in simpler organ-on-a-chip models. These early studies revealed the complexity of the glioblastoma microenvironment, paving the way for more complex organoid-based models and integrated sensors in the Emerging literature.

- **Chemokine Role in Lung TME (Jung, H., et al., 2024)** and **Immunosuppressive Cells in GI Cancers (Arshad, J., et al., 2024)**: Historical analyses used bulk tumor tissues and immunostaining to identify immunosuppressive cell types (e.g., MDSCs) and chemokine networks modulating immune infiltration. Emerging approaches like Zhang, Y., et al. (2024)’s lung cancer assembloids or De Zuani, M., et al. (2024)’s single-cell/spatial datasets in NSCLC build upon these findings. Now, microfluidic devices and single-cell RNA-seq (scRNA-seq) allow precise manipulation and tracking of these immune cells in engineered 3D environments, verifying the earlier chemokine and MDSC pathways identified historically.

- **Prostate Cancer Bone Metastases and MDSC Targeting (Zhou, W., et al., 2024)**: Historically, survival and therapy response data were collected at the bulk tissue or clinical endpoints. Emerging studies like Bian, X., et al. (2024) integrate single-cell RNA, ATAC-seq, and spatial transcriptomics to uncover cell-type-specific genetic and epigenetic alterations. This complements earlier clinical trial-based insights by mapping how metastatic niches evolve at single-cell resolution.

- **Pancreatic Cancer TME and Precision Targeting (Vitorakis, N., et al., 2024; Zhu, J., et al., 2024; Fu, Y., et al., 2024)**: Earlier works emphasized the complexity of the pancreatic TME, often limited to bulk expression analyses and immunostaining. The Emerging studies (e.g., Yang, G., et al., 2024 and Xie, J., et al., 2024) apply single-cell and spatial transcriptomics, and deep-learning-driven data integration, enabling "unbiased" decoding of complex cellular states. Now it is possible to quantify metabolic heterogeneity or pinpoint ILF2-driven metastatic niches in brain metastases of breast cancer using scRNA-seq and functional assays.

- **Renal and Colorectal Cancer Studies (Yang, G., et al., 2024; Wójcik, M., et al., 2024)** align with earlier findings from ovarian and hepatocellular carcinoma research (Garlisi, B., et al., 2024; Tang, Z., et al., 2024) that identified prognostic markers and coagulome gene sets. Emerging approaches confirm and refine these markers at single-cell resolution. For instance, the historical identification of immunosuppressive cell infiltration is now validated by single-cell analyses that show specific subsets of macrophages or loss-of-Y Tregs with unique immunosuppressive signatures.

- **Cancer-on-a-Chip and Coagulome Insights (Abed, H., et al., 2024; Tinholt, M., et al., 2024)**: Historical papers characterizing biomarkers and prognostic factors in ovarian and breast cancer now inform the design of microfluidic chips loaded with patient-derived cells. Emerging technologies like Maulana, T., et al. (2024)'s breast cancer-on-chip or Qian, H., et al. (2024)'s MRI-based radiomics now measure "~3,000 pg/mL" cytokine concentrations and correlate them with immune cell infiltration, improving upon earlier bulk-level prognostic data. Similarly, integration of imaging, computational modeling, and patient-specific assembloids (Zhang, Y., et al., 2024) allows for predictive drug testing that historical studies could only approximate.

**Key Insights**
1. **Refining Biomarkers and Prognostic Factors**:
Historical studies laid the groundwork by identifying prognostic factors—such as "M2 macrophage infiltration" (Qian, H., et al., 2024) or certain coagulome genes (Tinholt, M., et al., 2024)—but lacked the resolution to fully understand their spatial and cellular context. Emerging single-cell and spatial platforms bridge this gap by pinpointing which cell subsets express these genes and how their localization impacts prognosis. For instance,

Omilian, A., et al. (2024) found CD163+ macrophages linked to survival in triple-negative breast cancer; emerging technologies can now place these cells spatially and functionally in patient-matched assembloids.

2. **Validating and Improving Preclinical Models**:
The Historical research underscored the need for more physiologically relevant models. Emerging methods address this: Zhang, Y., et al. (2024) can generate "~200 uniform lung cancer assembloids in 1 minute" from tiny biopsies, a technical feat not previously possible. Maulana, T., et al. (2024) accurately model CAR-T cell infiltration in perfused breast cancer chips. These physiologically relevant organ-on-chip and assembloid models validate the earlier biomarker-driven hypotheses in controlled conditions that replicate in vivo complexity.

3. **Uncovering Novel Therapeutic Targets and Mechanisms**:
While Historical work identified general cellular players (e.g., MDSCs, CAFs, TAMs) and pathways implicated in multiple cancers, Emerging approaches delineate their functional heterogeneity. For example, Bian, X., et al. (2024)'s single-cell multi-omics in prostate cancer clarifies how distinct epithelial and immune cell subsets evolve, complementing earlier clinical trial results (Zhou, W., et al., 2024). Xie, J., et al. (2024) identified ILF2 in breast cancer brain metastases, a new target that may have been masked in bulk data. Such advances were enabled by the nuanced resolution and integrative computational methods not widely available at the time of the Historical studies.

4. **Personalized and Predictive Medicine**:
The Historical papers provided statistical correlations and generalizable patterns. Now, Emerging studies show patient-specific modeling. For example, lung cancer assembloids predicting clinical drug response (Zhang, Y., et al., 2024) directly complement earlier knowledge about chemokine-driven immunosuppression (Jung, H., et al., 2024) and MDSC-mediated resistance (Arshad, J., et al., 2024). Personalized organ-on-chip models can test how individual patient tumors respond to immunotherapies or chemotherapies, bridging the translational gap and moving toward precision oncology.

5. **Quantitative Validation and Multi-Modal Integration**:
Historical research identified key pathways but often lacked the tools to integrate multi-omic data. Emerging approaches use single-cell multi-omics, spatial transcriptomics, and deep learning (e.g., scMet in Yang, G., et al., 2024) to convert bulk data into single-cell resolution "over 15 billion RNA expression values." Such integration transforms historical macro-level observations into mechanistic insights at the single-cell level, enabling rational combination therapies targeting multiple components of the TME simultaneously.

**Explanation of Steps and Numerical Citations**:
To arrive at these conclusions, the analysis involved:

1. **Comparing Models**:
Historical papers often discussed limitations of traditional models. Emerging studies like Zhang, Y., et al. (2024) solved the sample size problem by producing "~200 uniform assembloids" from "10 µL hydrogel" in "1 minute."

2. **Linking Past Biomarkers with Present Technology**:
Historical insights into MDSC infiltration rates or chemokine expression patterns (Jung, H., et al., 2024; Arshad, J., et al., 2024) are now dissected using single-cell resolution. For example, Qian, H., et al. (2024) identified M2 macrophages as "HR = 32.288" risk factors for breast cancer patients. Emerging single-cell methods can specifically assign these cells to distinct niches, validating and explaining these large hazard ratios mechanistically.

3. **Spatial and Molecular Integration**:
Historical approaches lacked high spatial resolution. Emerging spatial transcriptomics (De Zuani, M., et al., 2024; Yang, G., et al., 2024) overcome this, making it possible to confirm previously identified prognostic factors at the exact tumor location.

4. **Functional Validation**:
Historical observations of therapy resistance and immunosuppression are now functionally tested on chips. For instance, Maulana, T., et al. (2024) demonstrated CAR-T infiltration "13-fold higher" than controls in a perfused breast cancer-on-chip model, validating earlier predictions of T cell exhaustion and infiltration rates.

5. **Citing Numerical Data from Authors**:
Each time a value is mentioned (e.g., "~200 uniform LCAs" from Zhang, Y., et al., or hazard ratios from Qian, H., et al.), it is placed in quotes corresponding to the authors' names, providing numerical anchoring to their claims. These numerical details confirm the improvements in precision and relevance of emerging methods compared to historical baselines.

---

In summary, the cutting-edge methods introduced in the 10 "Emerging" papers—such as patient-derived assembloids, single-cell and spatial multi-omics, cancer-on-chip platforms, and novel computational tools—build directly on and significantly enhance the foundational insights obtained from the 10 "Historical" studies. By providing finer resolution, physiological relevance, and patient-specific data, these emerging techniques complement and advance the field of organ-specific cancer research established by earlier work.

**ChatGPT o1 Details: 54 Words**

[Blending methods

I'm weaving together insights from 10 Emerging and 10 Historical papers to highlight how their different methods complement each other in organ-specific cancer research.

Structured synthesis

I'm mapping out a detailed report that blends insights from 10 Historical and 10 Emerging papers, showing the ways their complementary methods enhance organ-specific cancer research.]

### ChatGPT o1: 34.17s, 1432 words Triplicate 2

Below is a comprehensive, integrated report demonstrating how the innovative methodologies and approaches described in 10 "Emerging" Papers complement and advance the organ-specific cancer research laid out in 10 "Historical" Papers. The report is organized into "Executive Summary," "Technical Details," and "Key Insights." Throughout the explanation, we cite numerical data from authors' works using direct quotations with the authors' names, showcasing how these "Emerging" methods build upon and enhance the foundational knowledge established by the "Historical" papers.

**Executive Summary**
The 10 "Historical" Papers established critical frameworks for understanding organ-specific cancer microenvironments, including glioblastoma (GBM), lung cancer, gastrointestinal (GI) cancers, prostate cancer, pancreatic cancer, ovarian cancer, hepatocellular carcinoma (HCC), renal cell carcinoma, non-small cell lung cancer (NSCLC), and breast cancer. These earlier works focused on identifying key molecular drivers, immunosuppressive cells, angiogenic pathways, and therapeutic response markers within the tumor microenvironment (TME). They highlighted the complexity of each organ-specific niche and the need for improved in vitro and in vivo models, better biomarkers, and targeted therapy strategies.

The 10 "Emerging" Papers build directly on these foundations by integrating cutting-edge single-cell and spatial transcriptomics, advanced organoid and organ-on-a-chip platforms, patient-derived assembloids, biosensors, and radiomic analyses. These new methods enable finer resolution of cellular heterogeneity, maintain patient-specific tumor features, and provide non-invasive ways to predict therapeutic responses. By translating historical insights into actionable, high-fidelity experimental systems and predictive tools, the "Emerging" Papers push organ-specific cancer research toward precision oncology.

**Technical Details**

1. **Advanced Patient-Derived Models and Platforms**:

- Historical lung cancer studies (e.g., Jung et al. 2024; Thenuwara et al. 2024) identified key chemokine axes and complex tumor-immune interactions. Building upon this, “Emerging” methods such as the patient-specific lung cancer assembloid model developed by Zhang, Y. et al. (2024) generated “~200 uniform LCAs” from minimal clinical samples. These assembloids retain the “88% and 84% overall transcriptomic similarity” to parental tumors, surpassing historical 2D cultures by maintaining cellular diversity and TME complexity.
- Organ-on-a-chip systems historically proposed for GBM and other tumors are now integrated with biosensors (Thenuwara et al. 2024 “Historical”). In Maulana, T. et al. (2024 “Emerging”), a breast cancer-on-chip model allowed “8 days” of CAR-T cell perfusion and infiltration tracking, closely mimicking early clinical events and extending beyond simple static culture models.

2. **Multi-Omics and Spatial Profiling**:

- Early single-cell analyses in pancreatic and prostate cancers (Fu, Y. et al. 2024; Vitorakis, N. et al. 2024 “Historical”) revealed cell-type heterogeneity but lacked spatial context. Now, “Emerging” studies such as Bian, X. et al. (2024) for prostate cancer integrate single-cell RNA-seq, spatial transcriptomics, and bulk ATAC-seq to characterize “77,661 cells” from “7 PCa tissues” and delineate novel resistant cell states. Similarly, Yang, G. et al. (2024 “Emerging”) used “195 samples” and “981,294 cells” from “76 patients” to profile clear cell renal cell carcinoma, mapping metabolic states with unprecedented resolution.
- De Zuani, M. et al. (2024 “Emerging”) applied single-cell and spatial transcriptomics to nearly “900,000 cells” from NSCLC patients, identifying unique macrophage and epithelial subsets. This directly complements earlier NSCLC studies (Jung, H. et al. 2024 “Historical”) that highlighted chemokine-driven immunosuppression but lacked spatial and cell-type resolution.

3. **Immunoprofiling and Immune Cell Dynamics**:

- Prior research underscored immunosuppressive myeloid cell subsets and their impact on gastrointestinal (Arshad, J. et al. 2024 “Historical”) and ovarian cancers (Garlisi, B. et al. 2024 “Historical”). Emerging studies, such as Omilian, A. et al. (2024), used digital quantification of CD163+ macrophages in triple-negative breast cancer and found “HR = 0.30” improved survival, refining our understanding of tumor-associated macrophages first proposed historically.
- Wójcik, M. et al. (2024 “Emerging”) built on the concept of myeloid cell dysregulation by identifying loss of Y chromosome in Tregs within colorectal cancer and liver metastases, connecting genetic mosaicism to immunosuppression. This complements earlier GI cancer immunology frameworks set forth by Arshad, J. et al. (2024 “Historical”).

4. **Non-Invasive Biomarkers and Radiomics**:

- Historical studies on breast and prostate cancers stressed the need for better prognostic biomarkers (Zhou, W. et al. 2024; Tinholt, M. et al. 2024). Now, Qian, H. et al. (2024 “Emerging”) applied MRI-based radiomics to estimate M2 macrophage infiltration in breast tumors. With “HR = 32.288” for poor prognosis linked to M2 infiltration, radiomics offers a non-invasive extension to previous biomarker-driven research.
- Similarly, historical discussions of HCC biomarkers (Tang, Z. et al. 2024 “Historical”) are complemented by “Emerging” methods from Xie, J. et al. (2024), who used single-cell transcriptomics to identify ILF2 as a potential therapeutic target in breast cancer brain metastases. Though focused on a different organ, the approach parallels strategies for decoding HCC ceRNA networks.

5. **Refined Therapeutic Screening and Personalized Medicine**:
- Historical papers highlighted the complexity of targeting PDAC, with few effective therapeutics (Zhu, J. et al. 2024; Fu, Y. et al. 2024 “Historical”). Emerging works like Maulana, T. et al. (2024) and Zhang, Y. et al. (2024) develop personalized microfluidic models and assembloids that predict patient drug responses and immunotherapy effectiveness. Zhang et al.’s LCAs predicted clinical outcomes in “7 out of 7 patients,” bridging the gap between historical findings of drug resistance and practical personalized testing platforms.

**Key Insights**
1. **Enhanced Fidelity of Disease Models**:
“Emerging” patient-derived assembloids, organ-on-a-chip, and integrated multi-omic profiling tools refine the complexity observed in historical studies. Instead of relying on 2D cultures or animal models, these new techniques recapitulate patient-specific TME heterogeneity, genetic alterations, and immune dynamics with unprecedented realism.

2. **Spatial and Single-Cell Resolution**:
Historical recognition of the complexity of TMEs lacked spatial cellular context. The “Emerging” adoption of spatial transcriptomics and single-cell multiomics now maps cell subtypes, metabolic states, and immunosuppressive niches in situ, offering mechanistic insights into tumor evolution and therapy resistance directly corresponding to organ-specific microenvironments.

3. **Non-Invasive Prognostic Tools**:
While earlier studies identified molecular and immune markers of prognosis, “Emerging” radiomic and biosensor-based approaches provide new ways to monitor tumor status and predict therapeutic responses non-invasively. This dramatically increases clinical utility by integrating imaging and molecular profiling—offering earlier intervention and personalized treatment adjustments.

4. **Targeted Therapies Informed by TME Insights**:

Building on historical frameworks that identified key immunosuppressive cells and pathways, “Emerging” studies characterize novel gene targets (e.g., ILF2 in brain metastases), cell populations (LOY Tregs in CRC), and metabolic vulnerabilities. These refined targets emerge from a deeper understanding of the tumor’s cellular ecosystem, paving the way for more effective, context-specific therapies.

5. **Bridging Bench to Bedside**:
Historical findings often remained in preclinical conceptual frameworks. In contrast, the “Emerging” methodologies have begun validating these concepts in clinically relevant models, including patient-derived assembloids that mirror actual patient responses (Zhang, Y. et al. 2024), or breast cancer-on-a-chip models that simulate CAR-T therapy (Maulana, T. et al. 2024). This pipeline improvement accelerates translation from basic research to clinical trials and personalized medicine.

---

**Explanation of Steps and Data Citation**:
To arrive at these conclusions, we first reviewed the 10 “Historical” Papers, noting their principal findings: the identification of key immunosuppressive cell types (e.g., MDSCs in GI cancers), relevant signaling pathways (e.g., CXCR2 in lung cancer), and biomarker-based prognoses (e.g., coagulome genes in breast cancer). We then examined the 10 “Emerging” Papers, focusing on their methodologies—such as single-cell and spatial transcriptomics (Bian, X. et al. 2024; De Zuani, M. et al. 2024), patient-derived assembloids (Zhang, Y. et al. 2024), on-chip immunotherapy modeling (Maulana, T. et al. 2024), and radiomic analyses (Qian, H. et al. 2024).

By comparing these sets of studies:

- We noted that “Zhang, Y. et al.” generated “~200 uniform LCAs” with “88% and 84%” similarity to parental tumors, which addresses historical limitations where traditional models lost tumor heterogeneity.
- “Maulana, T. et al.” integrated CAR-T cells into a breast cancer-on-chip system, observing a “13-fold” increase in CAR-T infiltration compared to control T cells, a direct improvement over older systems lacking dynamic immune cell tracking.
- “Bian, X. et al.” analyzed “77,661 cells” and “Yang, G. et al.” integrated data from “981,294 cells” to reveal cell-state transitions in prostate and kidney cancers, surpassing historical population-level analyses by providing granular, single-cell insights.
- “Qian, H. et al.” correlated M2 macrophage infiltration with prognosis (HR = “32.288”), providing a predictive capability that complements historically established but less quantifiable prognostic biomarkers in breast cancer.

In each case, we enclosed numerical data in quotation marks and attributed them directly to authors, ensuring that the synergy between historical context and emerging techniques is firmly grounded in quantifiable, cited results.

In conclusion, the “Emerging” methods significantly enhance the foundational organ-specific cancer research from the “Historical” papers by providing more nuanced models, in-depth spatial and single-cell analyses, non-invasive predictive tools, and pathways to personalized therapy—ultimately accelerating the translation of bench research to clinical applications in precision oncology.

**ChatGPT o1 Details: 6 Words**

[Authors’ methods and organ-specific Cancer Research]

### ChatGPT o1: 75.46s, 1353 words Triplicate 3

Below is a comprehensive integrated report that shows how the cutting-edge techniques (“Emerging” Papers) build upon and further complement organ-specific cancer research approaches introduced in the “Historical” Papers. The format includes an **Executive Summary**, **Technical Details**, and **Key Insights**, followed by an explanation of the steps taken to reach these conclusions. Throughout, we cite specific numerical data and findings in **quotation marks** accompanied by authors’ names, as requested.

---

#### Executive Summary

Recent “Emerging” studies have introduced sophisticated patient-specific tumor models, advanced single-cell and spatial multi-omics technologies, novel organ-on-chip platforms, and non-invasive radiomic approaches. These innovations significantly enhance the foundational concepts developed in the “Historical” literature on organ-specific cancer biology and the tumor microenvironment (TME).

While the “Historical” Papers (e.g., Thenuwara et al. on glioblastoma-on-a-chip, Jung et al. on chemokines in lung cancer, and Arshad et al. on myeloid-derived suppressor cells in gastrointestinal cancers) established a deep understanding of cancer complexity and preliminary strategies for intervention, the “Emerging” Papers have elevated these concepts by:

- Achieving patient-specific, physiologically relevant tumor models that capture the full heterogeneity of the TME (Zhang et al., Maulana et al.).

- Leveraging single-cell multi-omics and spatial transcriptomics to decode intricate cell-cell interactions, metabolic states, and immune landscapes within organ-specific tumors (Bian et al., Yang et al., De Zuani et al., Xie et al.).
- Adopting radiomics and imaging-based biomarkers for non-invasive immune profiling and prognosis assessment (Qian et al.).
- Introducing novel therapeutic testing platforms (e.g., CAR-T efficacy on chip, assembloid models) and identifying new biomarkers (ILF2 in brain metastases, CD163+ macrophages in triple-negative breast cancer) to refine precision oncology strategies (Omilian et al., Xie et al.).

By integrating the robust conceptual frameworks provided by the “Historical” research—such as the importance of immunosuppressive cells (Arshad et al.), chemokine signaling (Jung et al.), and complex TME interactions (Garlisi et al., Tang et al.)—the “Emerging” studies now offer powerful experimental and computational methods. These methods enable refined, personalized approaches to patient care and drug discovery in organ-specific cancers, including lung, breast, prostate, colorectal, renal, pancreatic, and ovarian malignancies.

---

#### Technical Details

**Integration of Advanced 3D Models and Biosensor Platforms:**
The “Historical” literature introduced organ-on-chip models incorporating biosensors, as seen in Thenuwara et al.’s glioblastoma-on-chip, which monitored “energy metabolites such as oxygen, lactate, and glucose.” Building on this, “Emerging” work by Zhang et al. engineered lung cancer assembloids (LCAs) that can be generated in ~“1 minute” to produce “~200 uniform LCAs” from minimal biopsy material. Similarly, Maulana et al.’s breast cancer-on-chip integrated patient-derived organoids and CAR-T cells, enabling direct assessment of immunotherapy efficacy.

**From Bulk to Single-Cell and Spatial Omics:**
“Historical” Papers such as Fu et al. and Zhu et al. discussed how single-cell multiomics and spatial transcriptomics could deepen understanding of the TME in pancreatic cancer. The “Emerging” Papers like Bian et al. and Yang et al. fully realized this potential by integrating single-cell RNA-seq, spatial transcriptomics, and epigenomics. For instance, Bian et al. analyzed “77,661 cells” from “7 PCa tissues” and performed pseudotime trajectory analysis to uncover lineage relationships in prostate cancer. Yang et al. integrated “9 major scRNA-seq databases” with “981,294 cells” from “76 patients” with ccRCC to map metabolic heterogeneity. This surpasses earlier work limited to bulk analyses, enabling precise identification of cellular subpopulations and their metabolic states.

**Imaging and Non-Invasive Biomarkers:**

Historical studies often emphasized the complexity of the TME but lacked robust, non-invasive methods to assess it clinically. Qian et al. (Emerging) addressed this gap in breast cancer by using MRI-based radiomics to predict M2 macrophage infiltration, reporting it as an independent prognostic factor with hazard ratios “HR = 32.288” for poor survival. This complements earlier conceptual frameworks (e.g., Tang et al. on ceRNA networks in HCC, Garlisi et al. on ovarian TME complexity) by providing a quantifiable, translational imaging tool.

**Immune and Stromal Complexity Mapped at Single-Cell Resolution:**
Historical work by Jung et al. and Arshad et al. showed how chemokines and MDSCs shape immunosuppression in lung and GI cancers. Emerging studies like De Zuani et al. and Xie et al. leveraged single-cell and spatial methods to pinpoint immune cell subsets and their functional states. De Zuani et al. profiled “~900,000 cells” across 25 NSCLC patients, revealing subtype-specific immune checkpoints like TIGIT in LUAD and CD96 in LUSC. Meanwhile, Xie et al. identified ILF2 as a novel target in breast cancer brain metastases, validated by “ILF2-KD” models that showed significant reductions in metastatic burden.

**Personalized Therapy Testing:**
While older literature suggested the importance of TME-driven therapy resistance (Zhou et al., Vitorakis et al.), Emerging work now offers patient-specific models to test and predict therapy responses. Zhang et al.’s LCAs “accurately replicated clinical drug responses” for 7 out of 7 patients treated with various therapies. Maulana et al. showed CAR-T cell infiltration and lysis of breast cancer organoids correlated strongly with the expression of target antigens, enabling patient-specific immunotherapy screening.

---

#### Key Insights

1. **Refined Patient-Specific Models:**
   Emerging methods provide robust platforms—such as lung cancer assembloids and breast cancer-on-chip systems—to mirror patient tumor behavior, validating and extending historical organ-on-a-chip frameworks. This move from generic models to patient-tailored constructs enables testing of therapies more closely aligned with individual TME complexities.

2. **Comprehensive Molecular Profiling:**
   Single-cell multiomics and spatial transcriptomics, now widely applied, build upon the foundational understanding of TME cell heterogeneity established in historical literature. These new technologies can pinpoint specific cell types (e.g., CD163+ macrophages, ILF2+ malignant cells, unique CAF subsets) and metabolic states, offering actionable targets identified only conceptually before.

3. **Synergy Between Imaging and Molecular Data:**
   Emerging radiomic analyses complement historical insights into the TME by non-invasively quantifying prognostic immune infiltration patterns. Such synergy allows clinicians to track TME changes in real-time, bridging the gap between experimental complexity and clinical applicability.

4. **Therapeutic Advancement and Resistance Overcoming:**
   Building on the historical identification of immunosuppressive cells and signaling pathways (like MDSCs, chemokine axes), emerging single-cell and organ-on-chip studies can now test strategies—e.g., CAR-T cell infiltration and metabolic targeting—that directly address resistance mechanisms identified conceptually in older literature.

In sum, the “Emerging” Papers translate the foundational, organ-specific cancer biology insights from the “Historical” Papers into powerful experimental and computational platforms. This synergy accelerates the path to personalized oncology, where therapy decisions are guided by robust, patient-specific data and models.

---

#### Steps Taken to Arrive at These Conclusions

1. **Comparative Reading:**
   We first reviewed the “Historical” Papers, noting their emphasis on the complexity of the TME, the importance of stromal and immune cells, and the early development of organ-on-chip and multi-omics frameworks. For example, Thenuwara et al. discussed glioblastoma-on-chip with integrated sensors; Jung et al. highlighted chemokine-driven TME interactions in lung cancer; Arshad et al. focused on MDSCs in GI cancers.

2. **Extracting Numerical Data and Key Methods:**
   We then examined each “Emerging” Paper, extracting specific data points, such as Zhang et al.’s ability to produce “~200 uniform LCAs” in “~1 minute,” or De Zuani et al.’s profiling of “~900,000 cells” in NSCLC. We aligned these numbers with the earlier qualitative findings to show a leap from conceptual complexity to high-throughput, quantitative modeling.

3. **Identifying Complementary Themes:**
   Historical works often laid the conceptual groundwork (e.g., tumor-stromal-immune crosstalk), while Emerging studies provided precise, patient-specific, and integrative methods to test and apply these concepts. For instance, the

immunosuppressive role of macrophages identified historically (Omilian et al.) is now quantifiable and targetable with radiomics and scRNA-seq platforms (Qian et al., Xie et al.).

4. **Synthesizing Insights and Highlighting Synergy:**
By comparing technical achievements—like the single-cell resolution data from Bian et al. (Emerging) with earlier multi-omics perspectives suggested in Fu et al. and Zhu et al. (Historical)—we illustrated how multi-omic integration, patient-derived models, and advanced imaging tools directly build upon and enhance historical understandings.

5. **Final Integration into a Unified Framework:**
The final report weaves together these studies, showing a clear progression from broad conceptual and mechanistic insights in the "Historical" literature to refined, personalized methodologies in the "Emerging" literature that enable actionable, patient-centered cancer therapy development.

---

**In conclusion**, the synergy between the "Historical" and "Emerging" sets of papers propels organ-specific cancer research forward by bridging conceptual knowledge with innovative, patient-focused methodologies.

**ChatGPT o1 Details: 158 Words**

[Aligning work

I'm working through how methods in 10 "Emerging Papers" enrich organ-specific cancer research from 10 "Historical Papers," following a structured format.

Synthesizing insights

I'm combining insights from 10 "Emerging Papers" to enhance organ-specific cancer research, building upon the foundational insights from 10 "Historical Papers" to create a unified and enriching overview.

Combining insights

I'm compiling insights from 20 papers, merging "Emerging" methods with "Historical" research, and integrating authors' numerical data for a comprehensive overview.

Highlighting advancements

I’m comparing "Emerging" and "Historical" research, showcasing patient-specific models, novel immunotherapy methods, and advanced multi-omic analysis, integrating and building upon foundational organ-specific cancer research.

Creating the report

I’m crafting an integrated report on how "Emerging" methods enhance "Historical" cancer research. Highlighting patient-specific models, novel immunotherapy techniques, and advanced multi-omic analysis. Progress is evident.

Drawing connections

Revealing how "Emerging" techniques, like Zhang's droplet microfluidics, align with "Historical" biosensor OOC approaches, leading to enhanced patient-specific preclinical platforms and more accurate immunotherapy predictions.]
